# Cholesterol biosynthesis inhibition synergizes with AKT inhibitors in triple-negative breast cancer

**DOI:** 10.1101/2024.01.16.575899

**Authors:** Alissandra L. Hillis, Timothy D. Martin, Haley E. Manchester, Jenny M. Hogstrom, Na Zhang, Emmalyn Lecky, Nina Kozlova, Nicole S. Persky, David E. Root, Myles Brown, Karen Cichowski, Stephen J. Elledge, Taru Muranen, David A. Fruman, Simon T. Barry, John G. Clohessy, Ralitsa R. Madsen, Alex Toker

## Abstract

Triple-negative breast cancer (TNBC) is responsible for a disproportionate number of breast cancer deaths due to its molecular heterogeneity, high recurrence rate and lack of targeted therapies. Dysregulation of the phosphoinositide 3-kinase (PI3K)/AKT pathway occurs in approximately 50% of TNBC patients. We performed a genome-wide CRISPR/Cas9 screen with PI3Kα and AKT inhibitors to find targetable synthetic lethalities in TNBC. We identified cholesterol homeostasis as a collateral vulnerability with AKT inhibition. Disruption of cholesterol homeostasis with pitavastatin synergized with AKT inhibition to induce TNBC cytotoxicity *in vitro*, in mouse TNBC xenografts and in patient-derived, estrogen receptor (ER)-negative breast cancer organoids. Neither ER-positive breast cancer cell lines nor ER-positive organoids were sensitive to combined AKT inhibitor and pitavastatin. Mechanistically, TNBC cells showed impaired sterol regulatory element-binding protein 2 (SREBP-2) activation in response to single agent or combination treatment with AKT inhibitor and pitavastatin. This was rescued by inhibition of the cholesterol trafficking protein Niemann-Pick C1 (NPC1). NPC1 loss caused lysosomal cholesterol accumulation, decreased endoplasmic reticulum cholesterol levels and promoted SREBP-2 activation. Taken together, these data identify a TNBC-specific vulnerability to the combination of AKT inhibitors and pitavastatin mediated by dysregulated cholesterol trafficking. Our work motivates combining AKT inhibitors with pitavastatin as a therapeutic modality in TNBC.

## Main

Triple-negative breast cancer (TNBC) is an aggressive disease with the worst five-year survival rate of all breast cancer subtypes^1^. TNBC is characterized by lack of expression of the estrogen receptor (ER), progesterone receptor (PR) and human epidermal growth factor receptor 2 (HER2). TNBC often recurs on standard of care chemotherapy, so there is a pressing need to identify targetable vulnerabilities^2,3^. The phosphoinositide 3-kinase (PI3K)/AKT signaling pathway is hyperactivated in nearly 50% of TNBC cases and promotes cell growth, survival and metabolism^2–5^. Because of the high prevalence of mutations in the PI3K/AKT pathway across cancer types, inhibitors of multiple nodes of the pathway have been developed, including PI3K, AKT and mTOR inhibitors^6,7^. In 2023, the catalytic AKT inhibitor AZD5363 (capivasertib, Truqap) was FDA-approved in combination with endocrine therapy (fulvestrant, Faslodex) to treat subsets of patients with hormone receptor (HR)-positive breast cancer^8–10^. Yet, the clinical use of PI3K/AKT inhibitors across distinct cancer lineages remains limited, and their efficacy depends on identifying combination therapies to limit on-target toxicities and acquired resistance^6,11^. Thus far, attempts to combine PI3K/AKT inhibitors with other drugs in the clinic have been mostly limited to standard of care regimens^6,11,12^. Here, we present an unbiased approach to characterize synergistic drug combinations with PI3K/AKT inhibitors in TNBC.

### Genome-wide CRISPR/Cas9 screen identifies synergy with combined AKT inhibition and cholesterol homeostasis genes

To identify synthetic lethal combinations with PI3K/AKT inhibitors in TNBC, we performed a genome-wide CRISPR/Cas9 screen in *PIK3CA*-mutant (H1047L) TNBC cells (SUM159). Cells were treated for 72 hours with DMSO (vehicle) or cytostatic doses of the PI3Kα-selective inhibitor BYL719 (0.4 µM) or the catalytic AKT inhibitor GDC-0068 (3 µM) (Fig. 1a). As expected, *PTEN* knockout provided a growth advantage to cells treated with BYL719, but not to cells treated with GDC-0068. *TSC2* knockout conferred a growth advantage to cells treated with either BYL719 or GDC-0068. Conversely, *FOXM1* knockout synergized to impair cell growth in both drug arms (Fig. 1b). Gene set enrichment analysis (GSEA) identified cholesterol homeostasis as one of the top perturbed pathways in the AKT inhibitor arm of the screen (Fig. 1c). We performed a second CRISPR/Cas9 screen utilizing a minipool library targeting 197 hits from the top perturbed pathways in the AKT inhibitor arm of the genome-wide screen. We identified synergy with AKT inhibition and knockout of cholesterol homeostasis genes across three TNBC cell lines (Extended Data Fig. 1a-c). Specifically, knockout of the master lipogenic transcriptional regulators, *SREBF1* and *SREBF2*, synergized with GDC-0068, suggesting that AKT inhibition sensitizes TNBC cells to inhibition of cholesterol biosynthesis (Fig. 1d).

**Fig. 1.**
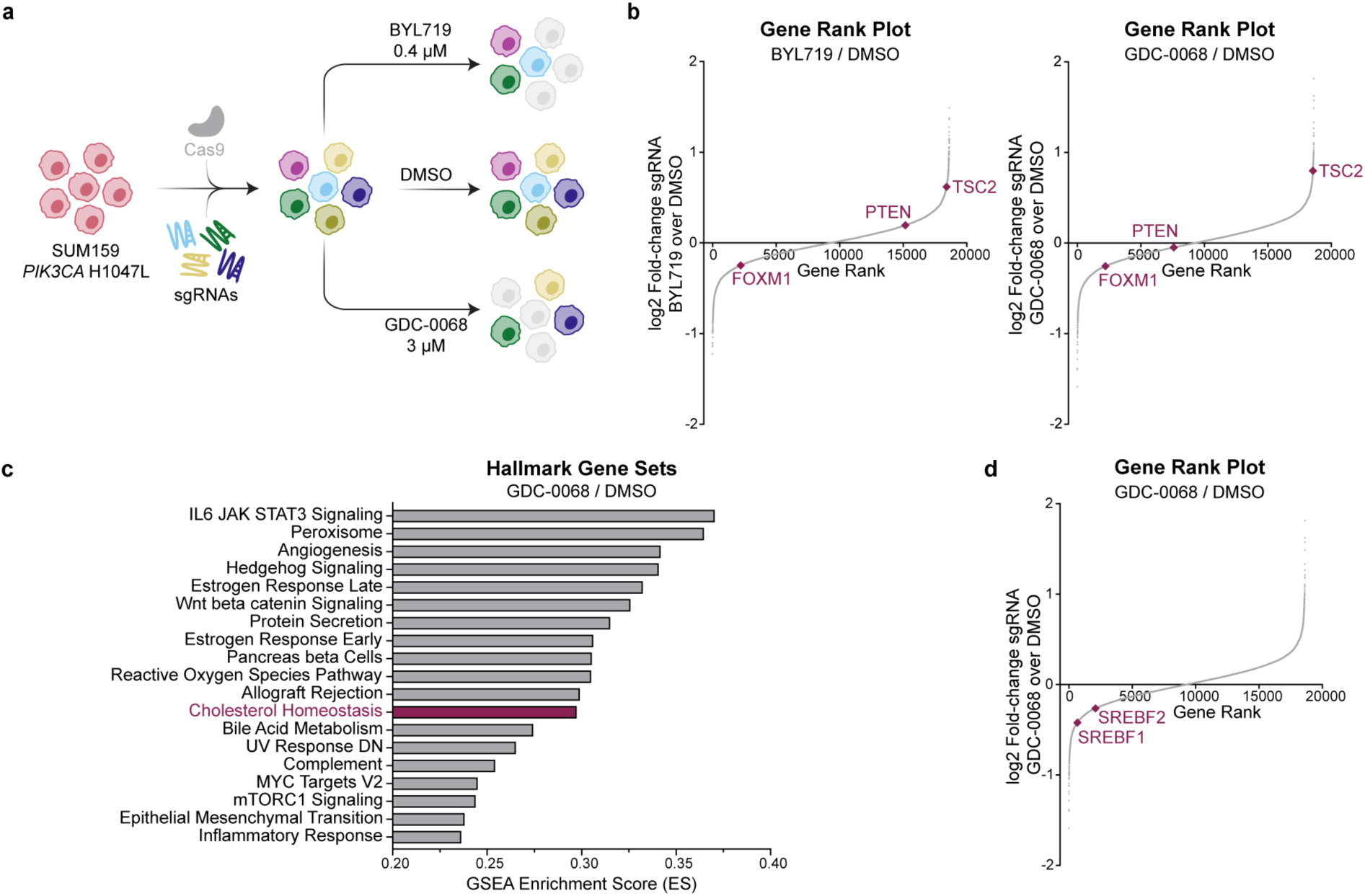
Genome-wide CRISPR/Cas9 screen identifies synergy with combined AKT inhibition and cholesterol homeostasis gene knockout in TNBC cells. **a,** Schematic of CRISPR/Cas9 screen. SUM159 cells were transduced with a Cas9-expressing lentivirus containing 94,495 sgRNAs with 3-4 sgRNAs per gene. Infected cells were allowed to grow for approximately 1 week before seeding the treatment arms. Cells were treated with DMSO, BYL719 (PI3Kα-selective inhibitor, 0.4 µM) or GDC-0068 (catalytic AKT inhibitor, 3 µM) for 72 hours (N=3 technical replicates for each treatment arm). **b,** Rank plots showing the log2 fold-change of each gene plotted against the dropout gene rank for the BYL719 and GDC-0068 treatments arms compared to the DMSO arm. Expected changes in PI3K/AKT signaling genes are highlighted, including *TSC2*, *PTEN* and *FOXM1*. Plots were generated using MAGeCK with a read count cutoff of 50 (N=3 technical replicates). **c,** Plot of the top pathways selectively perturbed in the GDC-0068 arm of the CRISPR/Cas9 screen. Analysis was performed via Gene Set Enrichment Analysis. **d,** Rank plot showing the log2 fold-change of each gene plotted against the dropout gene rank for the GDC-0068 treatment arm of the CRISPR/Cas9 screen compared to the DMSO arm. The transcription factors *SREBF1* and *SREBF2* are highlighted. The plot was generated using MAGeCK with a read count cutoff of 50.

### Disruption of cholesterol homeostasis synergizes with AKT inhibition in TNBC

Statins are a family of drugs that inhibit 3-hydroxy-3-methylglutaryl coenzyme A reductase (HMGCR), the first rate-limiting enzyme in cholesterol biosynthesis (Fig. 2a)^13^. In a panel of TNBC cell lines, the lipophilic statins pitavastatin and lovastatin and the hydrophilic statin rosuvastatin synergized with AKT inhibition (Fig. 2b-c, Extended Data Fig. 2a,c). Highest single agent (HSA) synergy score analysis for the combination of AKT inhibitors (GDC-0068, AZD5363) and statins showed significant synergy (Extended Data Fig. 2b,d, Extended Data Fig. 3a-b). Pitavastatin was more potent than lovastatin or rosuvastatin, synergizing with AKT inhibitors at 500-2000 nM concentrations, but all three statins showed similar degrees of synergy (Extended Data Fig. 2b,d, Extended Data Fig. 3a-b). In a 4-day growth curve, the combination of AZD5363 and pitavastatin outperformed DMSO or single agent at impairing proliferation and inducing cell death (Fig. 2d).

**Fig. 2.**
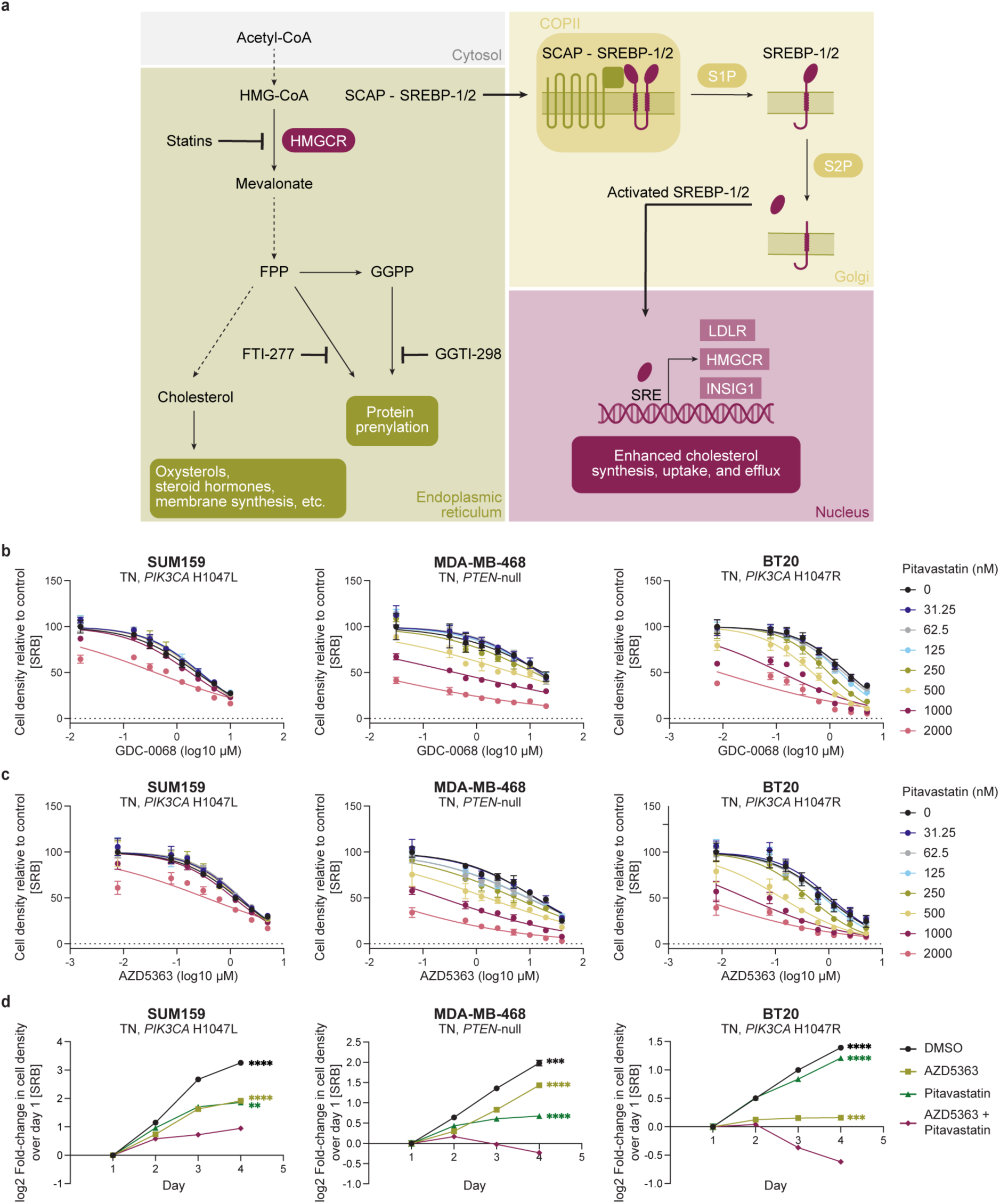
Disruption of cholesterol homeostasis synergizes with AKT inhibition in TNBC cells. **a,** Cholesterol is synthesized in multiple steps from acetyl-CoA. HMG-CoA reductase (HMGCR) catalyzes the rate-limiting step of cholesterol biosynthesis. This pathway also generates the prenylation substrates FPP and GGPP. SREBP-1/2 sense low endoplasmic reticulum cholesterol levels and translocate from the endoplasmic reticulum to the Golgi where they are cleaved and activated. N-terminal active SREBP-1/2 enter the nucleus to regulate the transcription of target genes. Drugs targeting this pathway are highlighted, including HMGCR inhibitors (statins) and inhibitors of protein farnesylation (FTI-277) and geranylgeranylation (GGTI-298). **b-c,** TNBC cell lines (SUM159, MDA-MB-468, BT20) were treated with increasing doses of GDC-0068 (SUM159: 0-10 µM, MDA-MB-468: 0-20 µM, BT20: 0-5 µM) (**b**) or AZD5363 (SUM159: 0-5 µM, MDA-MB-468: 0-40 µM, BT20: 0-5 µM) (**c**) and pitavastatin (0-2000 nM) for 72 hours, and cell density was measured by SRB assay. Data are represented as mean ± SD (N=3 technical replicates). **d,** TNBC cell lines (SUM159, MDA-MB-468, BT20) were treated with DMSO, AZD5363 (SUM159: 2.5 µM, MDA-MB-468: 10 µM, BT20: 1.25 µM), pitavastatin (SUM159: 4 µM, MDA-MB-468: 2 µM, BT20: 0.5 µM) or a combination of AZD5363 and pitavastatin for 72 hours, and cell density was measured daily by SRB assay. Data are represented as mean ± SD (N=3 technical replicates). Statistical analysis was performed using two-way analysis of variance (ANOVA) with Dunnett’s multiple comparison test; asterisks (*) indicate significant differences compared to the AZD5363 and pitavastatin combination treatment on day 4 (**, p = 0.0021, ***, p = 0.0002, ****, p < 0.0001).

We hypothesized that statins would also synergize with PI3K inhibitors, since PI3K acts upstream of AKT, and as expected, knockout of *SREBF1* and *SREBF2* synergized with BYL719 in the genome-wide CRISPR/Cas9 screen (Extended Data Fig. 4a). In a panel of TNBC cell lines, BYL719 synergized with pitavastatin (Extended Data Fig. 4b-c). Furthermore, Torin 1-mediated inhibition of mTORC1/2, which regulate sterol regulatory element-binding protein 1/2 (SREBP-1/2) activation downstream of PI3K/AKT, synergized with pitavastatin, suggesting that synergy between PI3K/AKT inhibitors and statins is mTORC1/2-dependent (Extended Data Fig. 4d-e). Non-tumorigenic mammary epithelial cells (MCF10A) and liver adenocarcinoma cells (HepG2) were less sensitive to the combination of AKT inhibitor and pitavastatin than TNBC cells, suggesting that this combination is not toxic to all cells (Extended Data Fig. 5a-b). Together, these data show that PI3K/AKT pathway inhibition synergizes with inhibition of cholesterol biosynthesis in TNBC.

### AKT inhibition synergizes with pitavastatin to induce TNBC cytotoxicity *in vitro*, in xenografts and in patient-derived organoids (PDOs)

Combination AZD5363 and pitavastatin treatment for 72 hours induced significant TNBC cell death compared to either single agent alone (Fig. 3a). To evaluate the efficacy of this combination in preclinical models, we used HCC70 TNBC xenografts. *In vitro*, AKT inhibitors synergized with pitavastatin in HCC70 cells (Extended Data Fig. 6a). HCC70 tumor-bearing mice were maintained on a low geranylgeraniol chow diet to limit the rescue of geranylgeraniol pyrophosphate (GGPP) levels after pitavastatin treatment^14,15^. Mice were treated with 100 mg/kg AZD5363 (4 days on, 3 days off) and 100 mg/kg pitavastatin (daily) by oral gavage. Single agent AZD5363 or pitavastatin did not significantly affect tumor growth. Combination AZD5363 and pitavastatin significantly impaired tumor growth and decreased tumor size and tumor percent body weight at endpoint (Fig. 3b-d, Extended Data Fig. 6b-c). Both drugs showed on-target efficacy as detected using markers of PI3K/AKT pathway activation (pAKT^Ser473^, pPRAS40^Thr246^, pAKT^Thr308^, pGSK3β^Ser9^, pS6^Ser240/244^), protein prenylation (RHEB, unprenylated RAP1A, HDJ2) and HMGCR (Extended Data Fig. 6d). In a panel of breast cancer PDOs, the combination of AZD5363 and pitavastatin impaired the growth of all PDOs, but induced cytotoxicity primarily in TN/ER-low PDOs (Fig. 3e, Extended Data Fig. 7a-d). Consistent with these results, pitavastatin-induced HMGCR upregulation was impaired in an ER-low PDO compared to an ER-positive PDO, and this was associated with a greater accumulation of unprenylated RAP1A (Fig. 3f). These data show that the combination of AZD5363 and pitavastatin induces cytotoxicity in preclinical models of TNBC.

**Fig. 3.**
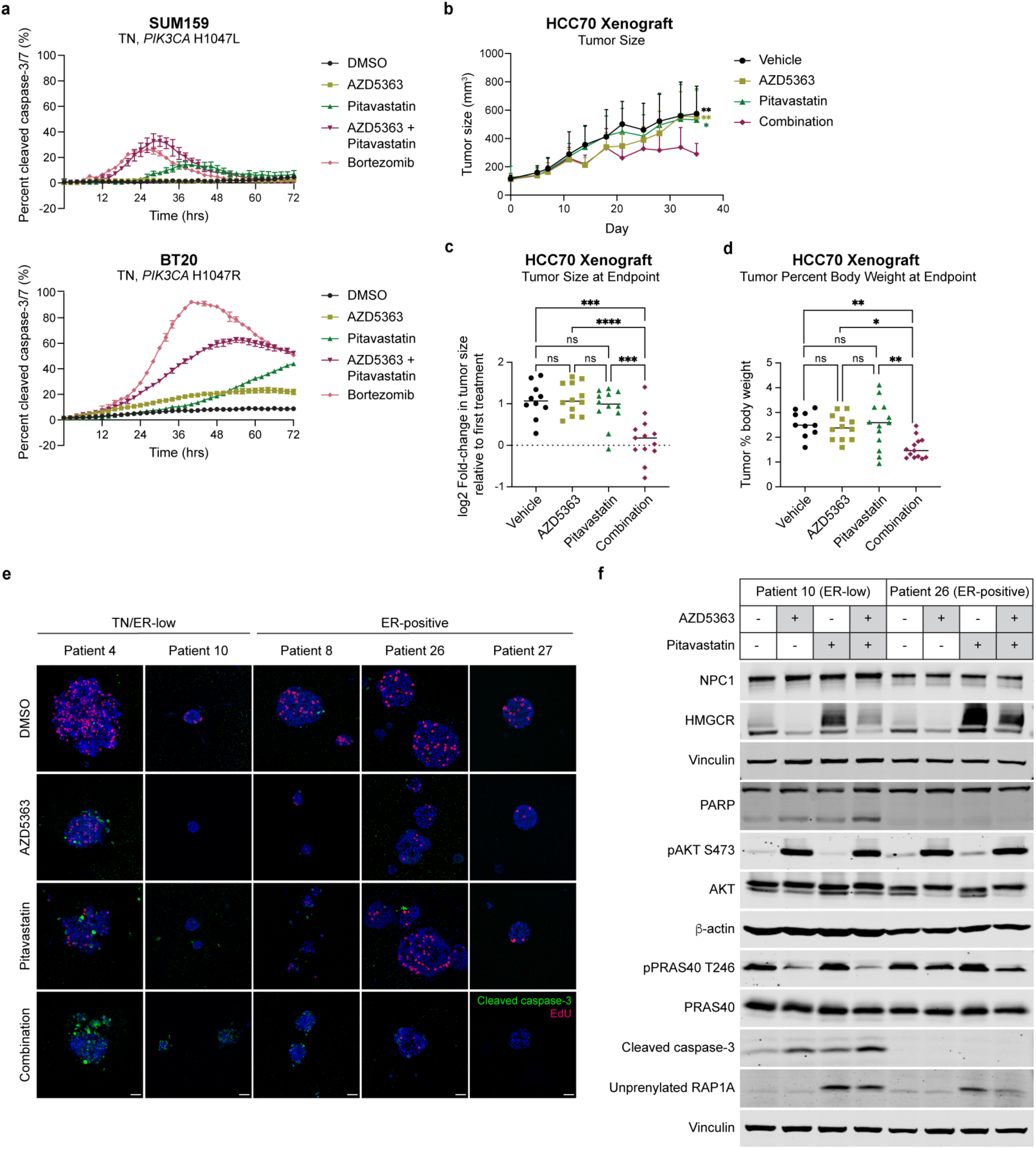
AKT inhibitors synergize with pitavastatin to induce TNBC cytotoxicity. **a,** TNBC cell lines (SUM159, BT20) were treated with DMSO, AZD5363 (SUM159: 5 µM, BT20: 1.25 µM), pitavastatin (SUM159: 4 µM, BT20: 2 µM) or a combination of AZD5363 and pitavastatin, and total cell number (rapid red nuclear dye) and number of dead cells (cleaved caspase-3/7 dye) were measured every 2 hours for 72 hours by Incucyte live-cell analysis. Data are represented as mean ± SD of percent cleaved caspase-3/7 signal (N=4 technical replicates). **b-d,** HCC70 cells were injected subcutaneously into NSG mice and tumors were allowed to grow for 21 days before starting treatments. Mice were switched to a low geranylgeraniol chow diet 3 days before starting treatments and were treated once daily with vehicle (0.5% carboxymethylcellulose, N=10), 100 mg/kg AZD5363 (4 days on, 3 days off, N=12), 100 mg/kg pitavastatin (daily, N=12) or both (N=13) for 24 days. **b,** Tumor size (mm^3^) was measured every 3-4 days. **c,** Tumor size (mm^3^) was measured at the endpoint. **d,** Tumor weight and mouse body weight were measured at the endpoint, and the tumor percent body weight was calculated by dividing tumor weight by mouse body weight. For **b-d**, statistical analysis was performed using two-way analysis of variance (ANOVA) with Tukey’s multiple comparison test; asterisks (*) indicate significant differences. For **b**, asterisks (*) indicate significant differences compared to the AZD5363 and pitavastatin combination treatment at the endpoint (*, p = 0.0332, **, p = 0.0021, ***, p = 0.0002, ****, p < 0.0001). **e,** A panel of breast cancer PDOs were treated with DMSO, 1 µM AZD5363, 5 µM pitavastatin or the combination of AZD5363 and pitavastatin for 96 hours and then pulsed with EdU and stained with a cleaved caspase-3 antibody. A representative image for each PDO in each treatment condition is shown. Scale bars are 40 µm. **f,** Immunoblots of NPC1, HMGCR, PARP, pAKT^Ser473^, pPRAS40^Thr246^, cleaved caspase-3, unprenylated RAP1A, β-actin and vinculin in an ER-low (patient 10) and ER-positive (patient 26) organoid treated with DMSO, 1 µM AZD5363, 5 µM pitavastatin or the combination for 24 hours.

### AKT inhibition does not synergize with pitavastatin in ER-positive breast cancer cells

We further tested the combination in a panel of PI3K/AKT pathway-mutant ER-positive breast cancer cell lines. ER-positive breast cancer cells were completely insensitive to the combination of AKT inhibitor and pitavastatin (Fig. 4a, Extended Data Fig. 8a-b). Similarly, ER-positive breast cancer cells were insensitive to the combination of BYL719 or Torin 1 plus pitavastatin (Extended Data Fig. 9a-d). Endocrine therapy (fulvestrant)-resistant T47D cells were exquisitely sensitive to AKT inhibitor and pitavastatin combination therapy, compared to the matched parental T47D cells, and this coincided with a loss of ER expression in fulvestrant-resistant cells (Fig. 4b, Extended Data Fig. 10a-b). In a panel of six TN and five ER-positive breast cancer cell lines, all TNBC cell lines were sensitive to combination catalytic (GDC-0068, AZD5363) or allosteric AKT (ARQ-092, MK-2206) inhibitor and pitavastatin, whereas none of the ER-positive breast cancer cell lines were sensitive (Fig. 4c). We hypothesized that ER expression contributes to statin resistance, however overexpression of ER in SUM159 and fulvestrant-resistant T47D cells did not affect sensitivity to pitavastatin or to the combination of AKT inhibitor plus pitavastatin (Extended Data Fig. 11a-b). Similarly, degradation of ER with fulvestrant did not sensitize ER-positive breast cancer cells to pitavastatin, indicating that ER expression is not sufficient to mediate resistance to the antiproliferative effects of statins (Extended Data Fig. 11c). This demonstrates that HR-negative breast cancer cells either have a unique dependency on the cholesterol biosynthesis pathway or have dysregulated cholesterol homeostasis.

**Fig. 4.**
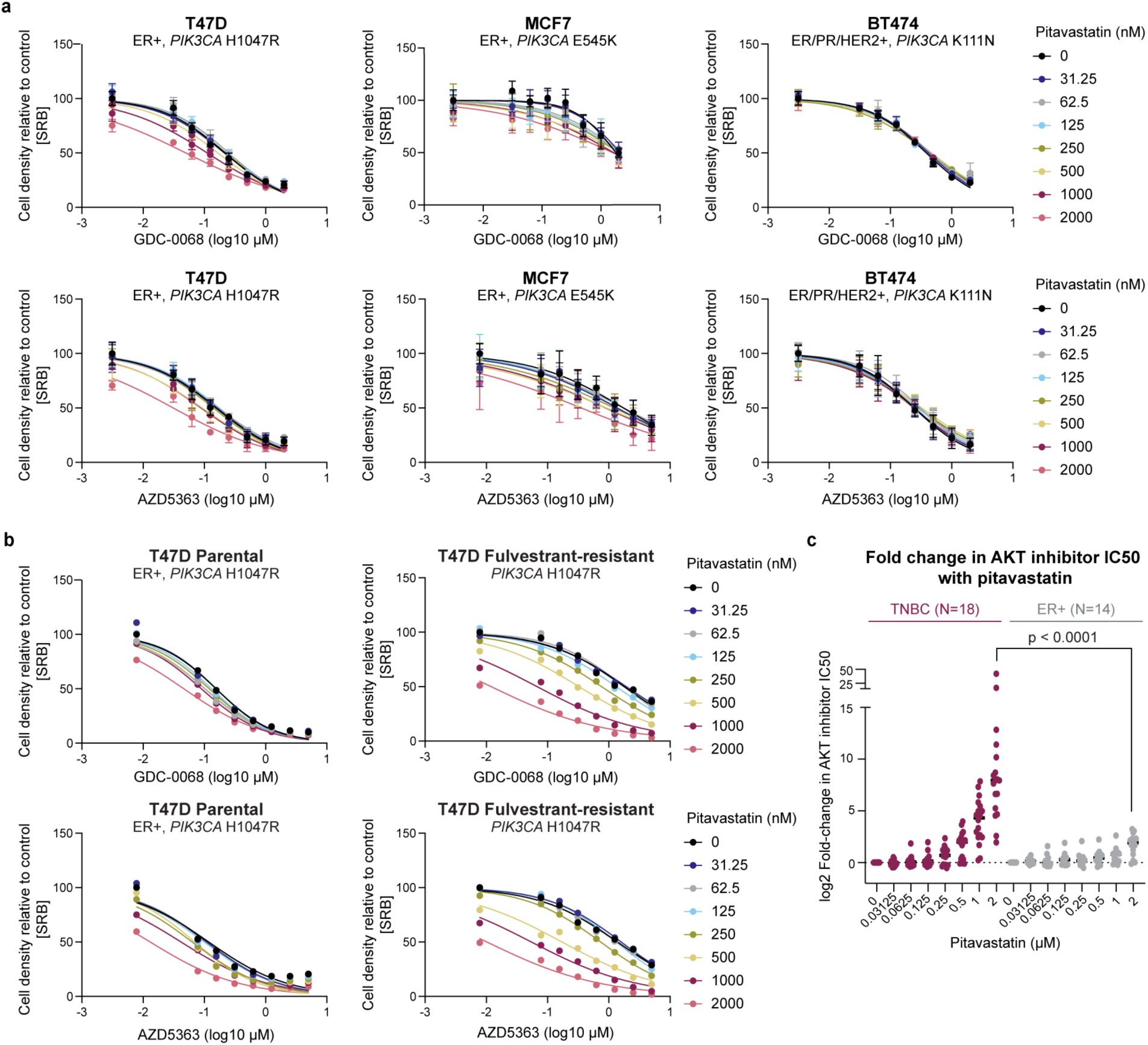
Pitavastatin does not synergize with AKT inhibition in ER-positive breast cancer cells. **a,** ER-positive breast cancer cell lines (T47D, MCF7, BT474) were treated with increasing doses of GDC-0068 (0-2 µM) or AZD5363 (T47D: 0-2 µM, MCF7: 0-5 µM, BT474: 0-2 µM) and pitavastatin (0-2000 nM) for 72 hours, and cell density was measured by SRB assay. Data are represented as mean ± SD (N=3 technical replicates). **b,** Parental and fulvestrant-resistant T47D cells were treated with increasing doses of GDC-0068 (0-5 µM) or AZD5363 (0-5 µM) and pitavastatin (0-2000 nM) for 72 hours, and cell density was measured by SRB assay (N=1 technical replicate). **c,** Log2 fold-change in AKT inhibitor IC50 (GDC-0068, AZD5363, MK-2206, ARQ-092) with 2 vs. 0 µM pitavastatin was calculated for six TNBC and five ER-positive breast cancer cell lines. Data are represented around the median (N = the number of cell line and AKT inhibitor combinations tested). Statistical analysis was performed for the 2 µM pitavastatin conditions using an unpaired, non-parametric Mann-Whitney test (p < 0.0001).

To measure the dependency of breast cancer cells on the cholesterol biosynthesis pathway, we treated a panel of TN and ER-positive breast cancer cells with the geranylgeranyltransferase-I (GGTase-I) inhibitor GGTI-298, which inhibits protein geranylgeranylation downstream of statins. Both TN and ER-positive breast cancer cells were sensitive to GGTI-298 and to the combination of AZD5363 and GGTI-298, suggesting that all breast cancer cells depend on the cholesterol biosynthesis pathway for survival through the generation of the prenylation substrate GGPP (Extended Data Fig. 12a-b). After statin treatment, but not GGTI-298 treatment, ER-positive breast cancer cells can maintain sufficient GGPP levels. By contrast, only RAS-altered breast cancer cells were sensitive to the farnesyltransferase (FTase) inhibitor, FTI-277 (Extended Data Fig. 13a-b). AZD5363 and pitavastatin-induced TNBC cell death was rescued by the addition of mevalonate, the product of HMGCR, or GGPP, further suggesting that loss of GGPP contributes to AKT inhibitor and pitavastatin-mediated cell death (Extended Data Fig. 12c-d). Cholesterol did not rescue cell death, but rather potentiated the cytotoxicity of AZD5363 and pitavastatin, likely by further inhibiting cholesterol biosynthesis through negative feedback (Extended Data Fig. 12c, Extended Data Fig. 14a)^16,17^. Consistent with this hypothesis, supplementing ER-positive breast cancer cells with exogenous cholesterol sensitized these cells to pitavastatin (Extended Data Fig. 14b). Altogether, these data show that the combination of AKT inhibition and pitavastatin induces apoptotic cell death by critically depleting GGPP in TNBC cells, which display dysregulated cholesterol homeostasis.

### Impaired SREBP-2 activation sensitizes TNBC cells to statins

To characterize the differential sensitivity of TN and ER-positive breast cancer cells to combined AKT inhibitor and pitavastatin, we performed bulk RNA-sequencing in statin-sensitive TNBC cells (MDA-MB-468) and statin-resistant ER-positive breast cancer cells (T47D) treated with single agent AKT inhibitor (AZD5363), pitavastatin or the combination for 24 or 48 hours. Statin-resistant ER-positive cells rapidly upregulated and sustained expression of *SREBF2* target genes, including *HMGCR*, in response to pitavastatin or combination AZD5363 and pitavastatin (Fig. 5a, Extended Data Fig. 15a-d, Extended Data Fig. 16c-d). This is the canonical response to low sterol conditions, which is mediated by transport of the SREBP cleavage-activating protein (SCAP)-SREBP-2 complex to the Golgi, SREBP-2 cleavage and subsequent translocation of active SREBP-2 to the nucleus where it promotes the transcription of target genes, including *LDLR*, *INSIG1* and *HMGCR*^17–19^. By contrast, statin-sensitive TNBC cells were unable to increase the expression of *SREBF2* target genes as potently or rapidly as ER-positive cells in response to pitavastatin or combination AZD5363 and pitavastatin (Fig. 5a, Extended Data Fig. 15a-d, Extended Data Fig. 16a-b). Depletion of *SREBF2*, but not *SREBF1*, further sensitized TNBC cells to pitavastatin and sensitized ER-positive breast cancer cells to pitavastatin and combination AKT inhibitor and pitavastatin by impairing pitavastatin-induced HMGCR upregulation (Fig. 5b, Extended Data Fig. 17a). Similarly, SREBP-1/2 inhibition with fatostatin, which binds SCAP and inhibits the transport of SREBP-1/2 from the endoplasmic reticulum to the Golgi, impaired HMGCR upregulation after combination AZD5363 and pitavastatin treatment and sensitized ER-positive breast cancer cells to pitavastatin (Extended Data Fig. 17b-c). Over a 24-hour time course of pitavastatin treatment, ER-positive breast cancer cells upregulated HMGCR mRNA and protein levels more significantly than TNBC cells, and higher HMGCR expression was associated with reduced unprenylated RAP1A levels (Fig. 5c, Extended Data Fig. 18a).

**Fig. 5.**
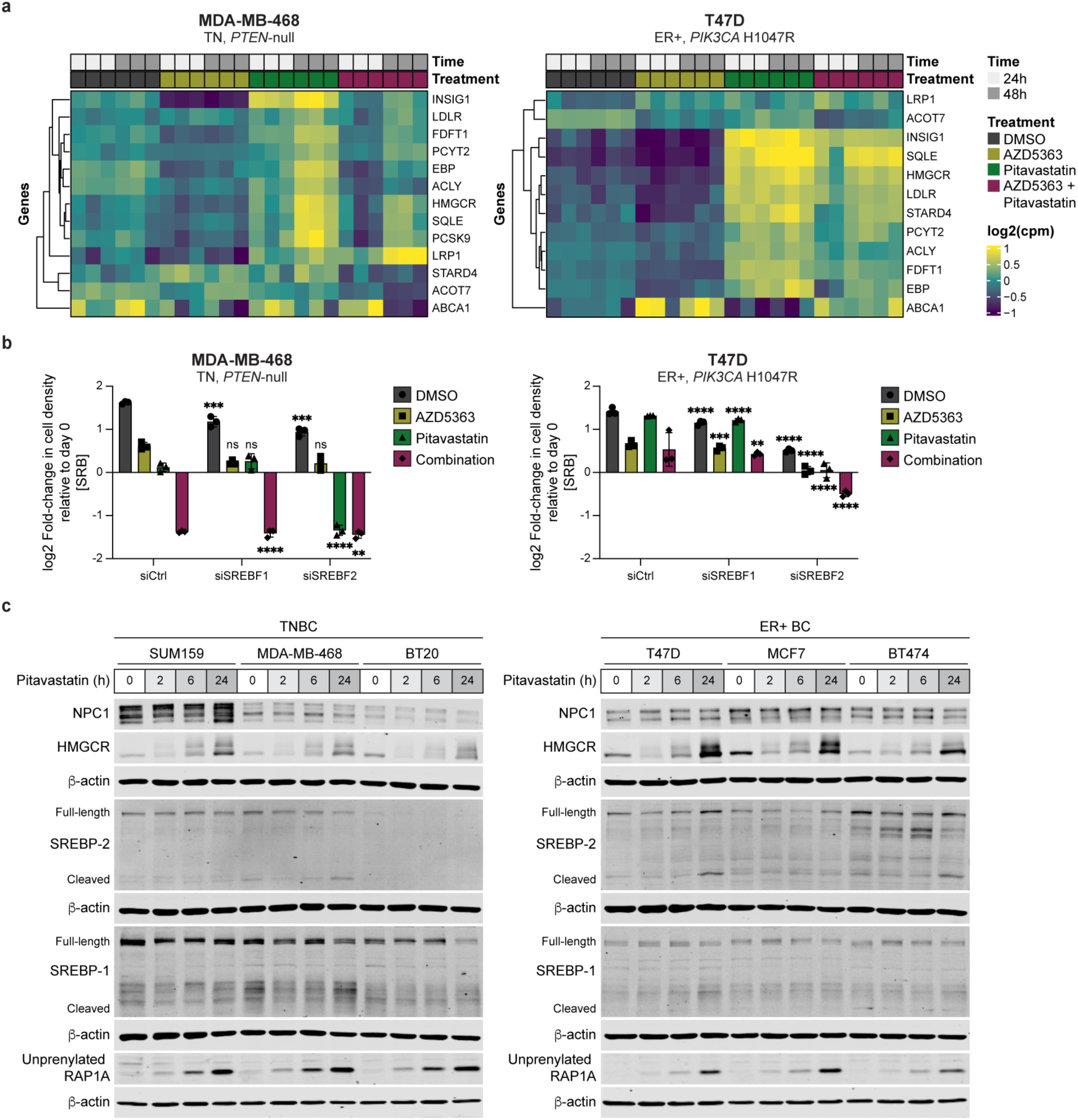
TNBC cells have impaired pitavastatin-induced SREBP-2 activation. **a,** RNA-sequencing was performed in TN (MDA-MB-468) and ER-positive (T47D) breast cancer cells treated with DMSO, AZD5363 (MDA-MB-468: 10 µM, T47D: 0.25 µM), pitavastatin (1 µM) or a combination of AZD5363 and pitavastatin for 24 or 48 hours. Data for *SREBF2* target genes are represented as log2 counts per million (cpm) for each replicate (N=3 biological replicates per condition). **b,** TN (MDA-MB-468) and ER-positive (T47D) breast cancer cells were transfected with siControl (siCtrl), siSREBF1 or siSREBF2 for 24 hours and then treated with DMSO, AZD5363 (MDA-MB-468: 15 µM, T47D: 0.25 µM), pitavastatin (2 µM) or a combination of AZD5363 and pitavastatin for 72 hours, and cell density was measured by SRB assay. Data are represented as mean ± SD (N=3 technical replicates). Statistical analysis was performed using two-way analysis of variance (ANOVA) with Tukey’s multiple comparison test; asterisks (*) indicate significant differences compared to the matched treatment condition in the siCtrl cells (**, p = 0.0021, ***, p = 0.0002, ****, p < 0.0001). **c,** Immunoblots of NPC1, HMGCR, SREBP-1/2, unprenylated RAP1A and β-actin in TN (SUM159, MDA-MB-468, BT20) and ER-positive breast cancer (T47D, MCF7, BT474) cell lines treated with DMSO or 2 µM pitavastatin for 2, 6 or 24 hours.

To test whether HMGCR expression is sufficient to confer resistance to pitavastatin, we attempted to overexpress wild-type (WT) or catalytically inactive HMGCR in TN or ER-positive breast cancer cells. However, intracellular cholesterol levels are tightly regulated, thereby significantly limiting HMGCR overexpression, to the extent that expression of WT HMGCR did not desensitize TNBC cells to pitavastatin (Extended Data Fig. 18b-c). To determine whether HMGCR expression is necessary to confer resistance to pitavastatin, we depleted *HMGCR* in a panel of TN and ER-positive breast cancer cells and treated them with AZD5363, pitavastatin or the combination. *HMGCR* depletion further sensitized TNBC cells to single agent pitavastatin but did not potentiate the cytotoxic effects of combination AZD5363 and pitavastatin (Extended Data Fig. 18d). By contrast, *HMGCR* depletion significantly sensitized ER-positive breast cancer cells to pitavastatin and induced cytotoxicity to combination AZD5363 and pitavastatin (Extended Data Fig. 18e).

Since AKT is known to regulate SREBP-2 activation, we hypothesized that AKT inhibitors synergize with pitavastatin in TNBC through enhanced suppression of pitavastatin-induced HMGCR upregulation^20–22^. In a panel of TNBC cell lines, AZD5363 decreased HMGCR expression and pitavastatin increased HMGCR expression. The combination of AZD5363 and pitavastatin impaired pitavastatin-induced HMGCR expression, consistent with impaired SREBP-2 activation (Extended Data Fig. 19a). These data suggest that statin-induced HMGCR upregulation occurs through SREBP-2 activation, however HMGCR is also regulated by proteasomal degradation. Pre-treatment of TN and ER-positive breast cancer cells with the proteasome inhibitor MG132 before a 24-hour treatment with pitavastatin or combination AZD5363 plus pitavastatin did not increase pitavastatin-induced HMGCR upregulation. Pre-treatment with the translation inhibitor cycloheximide (CHX) abolished pitavastatin-induced HMGCR upregulation, indicating that new synthesis accounts for increased HMGCR expression after statin treatment (Extended Data Fig. 19b-c). Collectively, these data show that AKT inhibitors synergize with pitavastatin in TNBC by potently suppressing SREBP-2 activation, resulting in decreased flux through the cholesterol biosynthesis pathway.

### NPC1 inhibition causes lysosomal cholesterol accumulation and rescues pitavastatin sensitivity

SREBP-2 activity is primarily regulated by cholesterol levels in the endoplasmic reticulum. We reasoned that perturbing intracellular cholesterol trafficking would alter SREBP-2 activity and pitavastatin sensitivity. Treatment of TN and ER-positive breast cancer cells with chloroquine, an autophagy inhibitor that causes lysosomal cholesterol accumulation, increased HMGCR expression, and *SREBF2* depletion abolished this increase (Extended Data Fig. 19d). Increased HMGCR expression induced by chloroquine and pitavastatin rescued the antiproliferative effects of pitavastatin in TNBC cells but did not alter pitavastatin sensitivity in ER-positive breast cancer cells (Extended Data Fig. 19e).

We next treated cells with OSW-1, a potent inhibitor of the oxysterol-binding protein (OSBP), which transports cholesterol out of the endoplasmic reticulum. Pitavastatin-sensitive breast cancer cells were more sensitive to OSW-1 than ER-positive, pitavastatin-resistant cells. Moreover, treatment with low dose OSW-1 (0.1 nM) sensitized ER-positive breast cancer cells to pitavastatin (Fig. 6a, Extended Data Fig. 20a). OSW-1 does not induce endoplasmic reticulum stress as a single agent or in combination with pitavastatin, suggesting that the antiproliferative effects are specifically due to endoplasmic reticulum cholesterol accumulation (Extended Data Fig. 20b). To standardize baseline endoplasmic reticulum cholesterol levels across TN and ER-positive breast cancer cells, we cultured cells in media with 10% lipid-depleted serum, containing no cholesterol, thereby limiting the cellular source of cholesterol to *de novo* synthesis. Concomitant addition of high dose pitavastatin (10 µM) for 1 hour inhibits cholesterol synthesis, resulting in selective depletion of cholesterol from the endoplasmic reticulum. After standardizing endoplasmic reticulum cholesterol levels, cells were treated with low dose pitavastatin for 72 hours in media supplemented with 10% lipid-depleted serum or cholesterol-rich 10% fetal bovine serum (complete serum). TNBC cells maintained in lipid-depleted serum were desensitized to pitavastatin, indicating that endoplasmic reticulum cholesterol levels determine statin sensitivity (Fig. 6b). By contrast, TNBC cells treated with low dose pitavastatin in complete serum remained pitavastatin sensitive, suggesting that these cells took up and trafficked exogenous cholesterol to the endoplasmic reticulum, thereby limiting SREBP-2 activation (Extended Data Fig. 20c).

**Fig. 6.**
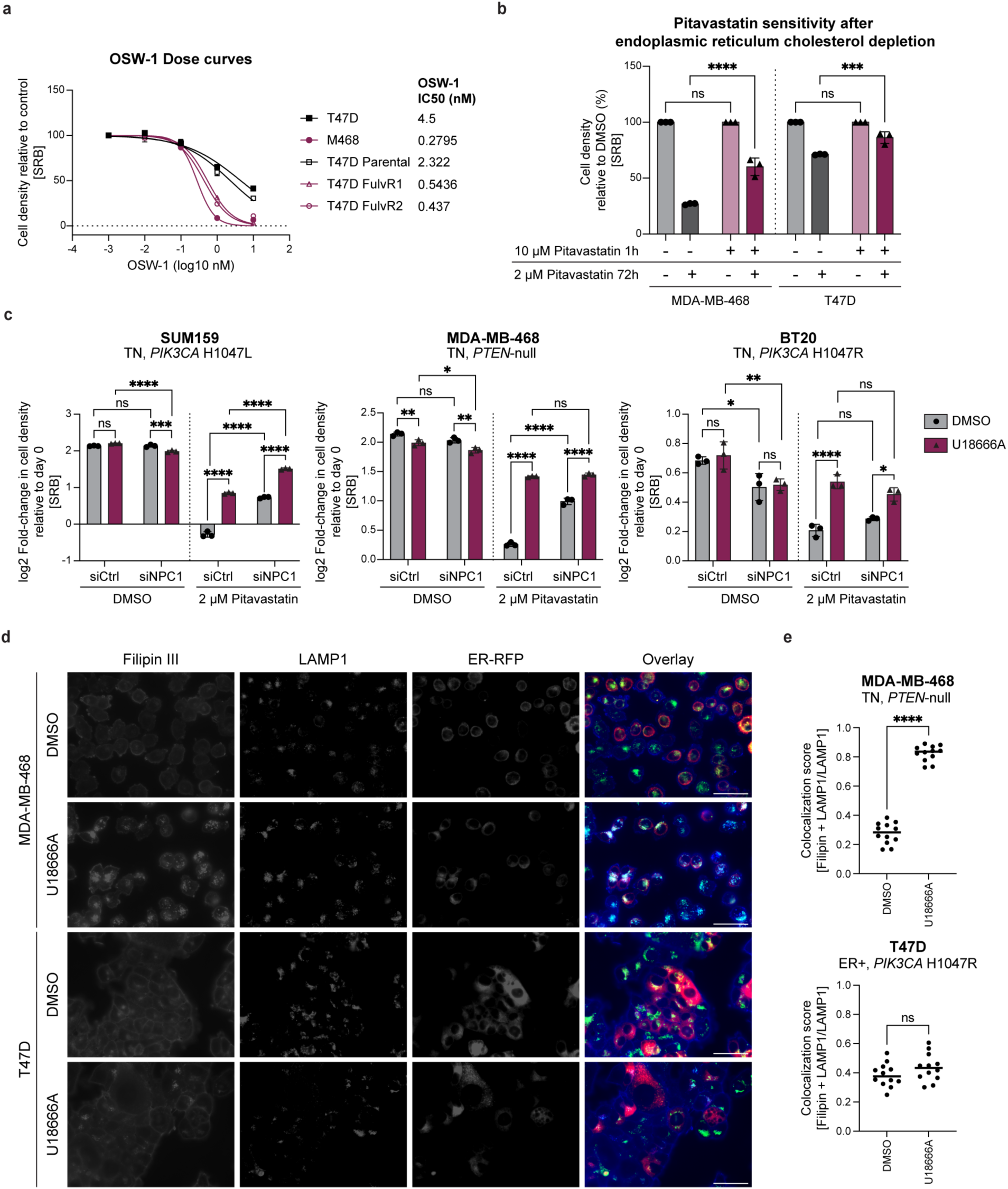
NPC1 inhibition causes lysosomal cholesterol accumulation and rescues pitavastatin sensitivity. **a,** ER-negative (MDA-MB-468, T47D fulvestrant-resistant clone 1 and 2) and ER-positive (T47D, parental T47D) breast cancer cell lines were treated with a range of concentrations of OSW-1 (0-10 nM) for 72 hours, and cell density was measured by SRB assay. Data are represented as mean ± SD (N=3 technical replicates). IC50 values for each cell line are reported. **b,** TN (MDA-MB-468) and ER-positive (T47D) breast cancer cells were seeded into media supplemented with 10% lipid-depleted serum and treated for 1 hour with vehicle or high dose pitavastatin (10 µM). Media was removed and replaced with media supplemented with 10% lipid-depleted serum and vehicle or low dose pitavastatin (2 µM) for 72 hours, and cell density was measured by SRB assay. Data are represented as mean ± SD (N=3 technical replicates). Statistical analysis was performed using two-way analysis of variance (ANOVA) with Šidák’s multiple comparison test (***, p = 0.0002, ****, p < 0.0001). **c,** TNBC cells (SUM159, MDA-MB-468, BT20) were transfected with siControl (siCtrl) or siNPC1 and then treated with DMSO or 1 µM U18666A and DMSO or 2 µM pitavastatin for 72 hours, and cell density was measured by SRB assay. Data are represented as mean ± SD (N=3 technical replicates). Statistical analysis was performed using two-way analysis of variance (ANOVA) with Šidák’s multiple comparison test (*, p = 0.0332, **, p = 0.0021, ***, p = 0.0002, ****, p < 0.0001). **d,** TN (MDA-MB-468) and ER-positive (T47D) breast cancer cells expressing endoplasmic reticulum RFP (ER-RFP) were treated with DMSO or 1 µM U18666A for 24 hours. Cells were fixed with 4% formaldehyde and stained with Filipin III and a LAMP1 antibody. Representative images are shown. Scale bars are 50 µm. **e,** Quantification of Filipin III and LAMP1 co-localization normalized to total LAMP1 from 10 non-overlapping fields. Statistical analysis was performed using an unpaired, two-tailed parametric t-test (****, p < 0.0001).

The cholesterol trafficking protein Niemann-Pick C1 (NPC1) transports cholesterol from the lysosome to the endoplasmic reticulum. Depletion or inhibition of NPC1 results in lysosomal cholesterol accumulation and decreased endoplasmic reticulum cholesterol levels. In a panel of TNBC cell lines, siRNA-mediated depletion of NPC1 or treatment with the NPC1 inhibitor U18666A rescued the antiproliferative effects of pitavastatin (Fig. 6c). Since U18666A can inhibit other cholesterol trafficking proteins beyond NPC1, we depleted a panel of genes reported to be inhibited by U18666A and evaluated pitavastatin sensitivity. Depletion of *GRAMD1A/B/C* or *OSBPL9* did not rescue pitavastatin sensitivity, indicating that loss of NPC1 alone is sufficient to rescue the antiproliferative effects of pitavastatin (Extended Data Fig. 21a). Combined depletion of NPC1 and treatment with U18666A did not significantly outperform depletion or inhibitor alone, further demonstrating that NPC1 alone can mediate pitavastatin sensitivity in TNBC (Fig. 6c). Depletion of *SREBF2* abrogated the NPC1 inhibitor-mediated rescue of pitavastatin in TNBC, suggesting that NPC1 inhibition activates SREBP-2 (Extended Data Fig. 21b). NPC1 depletion or inhibition in combination with pitavastatin increased HMGCR upregulation, and this increase was abolished upon depletion of *SREBF2* (Extended Data Fig. 21c). These data indicate that NPC1 inhibition rescues pitavastatin sensitivity by decreasing endoplasmic reticulum cholesterol levels to promote SREBP-2 activation and subsequent HMGCR upregulation.

Finally, we visualized subcellular cholesterol localization in TN and ER-positive breast cancer cells expressing RFP in the endoplasmic reticulum and stained with Filipin III (a cholesterol stain) and LAMP1 (a lysosomal marker). At baseline, ER-positive breast cancer cells showed elevated lysosomal cholesterol levels compared to TNBC cells. TNBC cells treated with U18666A rapidly accumulated lysosomal cholesterol, whereas the fraction of cholesterol-positive lysosomes in ER-positive breast cancer cells did not significantly increase from baseline (Fig. 6d-e). NPC1 inhibition decreases endoplasmic reticulum cholesterol levels in TNBC by trapping cholesterol in the lysosome, resulting in enhanced SREBP-2 activation and loss of sensitivity to pitavastatin. Together, these data support a model in which TNBC cells are uniquely sensitive to statins due to elevated endoplasmic reticulum cholesterol, which results in impaired SREBP-2 activation in response to statin. Combination AKT inhibitor and statin more potently suppresses SREBP-2 activation, resulting in cytotoxicity.

## Discussion

Given the prevalence of PI3K/AKT pathway hyperactivation in TNBC and the clinical potential of PI3K/AKT inhibitors, we leveraged an unbiased, genome-scale screen to identify collateral vulnerabilities using PI3K/AKT inhibitors as anchor drugs. We identified synergy between AKT inhibition and disruption of cholesterol homeostasis genes, including *SREBF2*, a master transcriptional regulator of cholesterol biosynthesis. Inhibition of cholesterol biosynthesis with statins synergized with AKT inhibitors in a panel of TNBC cell lines. The combination of AKT inhibitor and pitavastatin induced apoptotic cell death in TNBC cells, mouse xenografts of TNBC and PDOs of ER-negative breast cancer. ER-positive breast cancer cells and PDOs were resistant to pitavastatin and the combination of AKT inhibitor and pitavastatin. Previous studies have also shown that HR-negative breast and prostate cancer cells are hypersensitive to statins due to impaired SREBP-2 activation^23–25^. Here, we found that ER expression alone is not sufficient to mediate resistance to statins, nor is ER degradation sufficient to sensitize ER-positive breast cancer cells to statins. Rather, subcellular cholesterol localization determines statin sensitivity. Higher lysosomal and lower endoplasmic reticulum cholesterol levels in ER-positive breast cancer cells allow for rapid activation of SREBP-2 upon statin treatment. By contrast, TNBC cells have altered cholesterol trafficking with reduced lysosomal and enhanced endoplasmic reticulum cholesterol levels, resulting in statin sensitivity that was rescued by depletion of endoplasmic reticulum cholesterol.

While statins have been FDA-approved for the treatment of hypercholesterolemia since the 1980s, epidemiological and clinical data on the efficacy of statins in cancer is inconclusive^26–30^. Previous studies have reported that lipophilic statins exert anti-cancer effects in preclinical models, yet no conclusive mechanisms of sensitivity have been described^31–38^. Our findings explain the apparent failure of statins as anti-cancer agents in the clinic. Most clinical trials of statins in cancer have not evaluated pitavastatin, yet pitavastatin is the only statin that can reach cytotoxic anti-cancer concentrations in human plasma at clinically administered doses^30,39^. In addition to statin selection, patient diet has not been considered in clinical trials. Several studies in preclinical models have shown that diets rich in the GGPP precursor, geranylgeraniol, can restore cellular GGPP levels and rescue the antiproliferative effects of statins. Standard mouse chow is rich in oils that contain geranylgeraniol, which has likely confounded most preclinical studies of statins^14,15^. Lastly, our data highlight the importance of patient selection for statin clinical trials. Despite studies in cancer cell lines showing statin efficacy in TNBC, most trials have evaluated the efficacy of statins in HR-positive breast cancer^23,31^. Our studies suggest that lysosomal cholesterol levels and statin-induced HMGCR expression could serve as biomarkers of statin response.

Although millions of people worldwide are prescribed statins for cholesterol-lowering benefit, the epidemiological data are insufficient to definitively assess whether statin use affects incidence or outcomes in TNBC. TNBC accounts for approximately 12% of breast cancer cases and is more common in younger women who are less likely to be statin users^1^. Since pitavastatin is infrequently prescribed in the USA, there is not a large enough population of TNBC patients to perform any meaningful retrospective analysis on the role of pitavastatin in TNBC.

In summary, we have identified synergy between AKT inhibitors and pitavastatin in TNBC. By co-targeting TNBC cells with statins and AKT inhibitors, SREBP-2 activation and cholesterol biosynthesis are potently inhibited, further impairing upregulation of HMGCR expression, resulting in GGPP depletion and cytotoxicity. This combination is selective for TNBC cells because of dysregulated cholesterol homeostasis. Since both pitavastatin and AKT inhibitors (AZD5363, capivasertib) are FDA-approved drugs, this motivates the evaluation of the efficacy of this combination in clinical trials of TNBC.

## Methods

### Cell lines

The following commercially available cell lines were used: SUM159 (Asterand Bioscience/BioIVT, SUM159PT), MDA-MB-468 (ATCC, HTB-132), BT20 (ATCC, HTB-19), T47D (ATCC, HTB-133), MCF7 (ATCC, HTB-22), BT474 (ATCC, HTB-20), HCC70 (ATCC, CRL-2315), MCF10A (ATCC, CRL-10317), HepG2 (ATCC, HB-8065), BT549 (ATCC, HTB-122), HCC1937 (ATCC, CRL-2336), ZR-75-1 (ATCC, CRL-1500), MDA-MB-361 (ATCC, HTB-27), HEK 293T (ATCC, CRL-11268), parental T47D (Myles Brown lab), fulvestrant-resistant T47D clones 1-3 (Myles Brown lab). SUM159, MDA-MB-468, T47D, MCF7, BT474, HCC70, BT549, HCC1937, ZR-75-1 and parental T47D cells were cultured in RPMI-1640 medium (Gibco, 11875093) supplemented with 10% fetal bovine serum (FBS; GeminiBio, 100-106). Fulvestrant-resistant T47D cells were cultured in RPMI-1640 medium (Gibco, 11875093) supplemented with 10% fetal bovine serum (FBS; GeminiBio, 100-106) and 100 nM fulvestrant (Selleckchem, S1191). BT20 and HepG2 cells were cultured in Eagle’s Minimum Essential Medium (EMEM; Corning, 10009CV) supplemented with 10% FBS (GeminiBio, 100-106). MDA-MB-361 and HEK 293T cells were cultured in Dulbecco’s Modified Eagle’s Medium (DMEM) with L-Glutamine, 4.5 g/L glucose and sodium pyruvate (Fisher Scientific, MT10013CV) supplemented with 10% FBS (GeminiBio, 100-106). MCF10A cells were cultured in standard MCF10A growth medium without antibiotics (DMEM/F12 medium (Wisent Bioproducts, 319-075-CL), 5% horse serum (Gemini Bio, 100508), 10 µg/mL insulin (ThermoFisher Scientific/Gibco, A11382II), 0.5 mg/mL hydrocortisone (Sigma-Aldrich, H4001), 20 ng/mL EGF (R&D Systems, 236-EG-01M) and 100 ng/mL cholera toxin (List Biological Laboratories, 100B)). Cell lines were maintained at 37°C in a 5% CO_2_ cell culture incubator and passaged at 70-90% confluency. To passage, cells were washed once with 1X PBS and incubated for 5-10 minutes at 37°C with 0.25% Trypsin, 0.1% EDTA (Fisher Scientific, MT25053CI). Cells were passaged up to 5 times in the same dish and were maintained in culture for up to one month. Cells routinely tested negative for mycoplasma contamination.

### Genome-wide CRISPR/Cas9 screen

#### Viral transduction, cell seeding, drug treatments and harvest

SUM159 cells were transduced at a multiplicity of infection (MOI) of ∼0.3 with CRISPR/Cas9 lentivirus (Steve Elledge lab) containing 94,495 sgRNAs, with 3-4 sgRNAs per gene, and at least 500-fold representation. Cells were selected with 1 µg/mL puromycin (Corning, 61-385-RA) for 3 days. After selection, day 0 cell pellets were collected, and the remaining infected cells were expanded for 3 days and then plated into treatment arms at 100,000 cells/mL (2 x 10^6^ cells/plate) in tissue culture-treated 15-cm plates. Cells were treated for 72 hours with DMSO (vehicle) or cytostatic doses of the PI3Kα-selective inhibitor BYL719 (0.4 µM; Active Biochem, A-1214) or the catalytic AKT inhibitor, GDC-0068 (3 µM; Selleckchem, S2808). At the end point, cell pellets were harvested and processed as described below.

#### Genomic DNA isolation for sequencing

Cell pellets were resuspended in 10 mM Tris (Fisher Scientific, BP152-500) pH 8.0 and 10 mM EDTA (Fisher Scientific, E478-500) (TE buffer) to a final concentration of 2-10 million cells per 1 mL of TE buffer. Cell pellets were disrupted by pipetting. SDS (0.5% final concentration; AmericanBio, AB01920-00500) and proteinase K (0.5 mg/mL final concentration; Invitrogen, 25530049) were added, and cells were incubated in a 55°C water bath overnight, with a few inversions to promote cell lysis. When digestion was complete (homogenous, clear solution), NaCl (0.2 M final concentration; Fisher Scientific, BP358-10) was added. Phenol-chloroform/chloroform extraction was then performed using MaXtract High Density tubes (Qiagen, 129073). Tubes were pre-spun according to the manual. Samples were mixed with equal parts phenol:chloroform (Invitrogen, 15593031) in the MaXtract High Density tubes, shaken for 1 minute to extract and spun at 1500 x g for 5 minutes. The aqueous DNA phase separated on top. This extraction was repeated with chloroform (Fisher Scientific, C298-500), and the aqueous phase was transferred to a 50 mL conical tube. Conical tubes were incubated at 50°C for 1 hour to evaporate residual chloroform. RNase A (25 µg/mL final concentration; Qiagen, 19101) was added, and samples were incubated at 37°C overnight. The phenol:chloroform/chloroform extraction was repeated. DNA was precipitated by adding 1/10 v/v 3 M sodium acetate (Sigma-Aldrich, S2889) pH 5.2 and 2 volumes 100% ethanol (Pharmco, 111000200) and incubated at -20°C overnight. Samples were spun for 30-45 minutes at 4500 rpm at 4°C. DNA was washed 3 times with 1-1.5 mL 70% ethanol and allowed to dry open cap at 37°C for 10-20 minutes. DNA was resuspended in 1 mL of TE buffer by gently pipetting and incubating overnight at 55°C.

#### PCR amplification of genomic DNA

Genomic DNA was PCR amplified in 3 consecutive steps. All PCR was performed using Q5 Hot Start High-Fidelity DNA Polymerase (NEB, M0493L) and the Q5 Reaction Buffer Pack (NEB, B9027S). For PCR #1, 400 µg of DNA were added to the reaction for each condition with 6 µg of DNA per 50 µL reaction. PCR #1 primers amplified sgRNA sequences from the genomic DNA:

LC353F (forward): 5’ - AAT GGA CTA TCA TAT GCT TAC CGT AAC TTG AAA GTA TTT CG - 3’

LCR2L (reverse): 5’ - TCT ACT ATT CTT TCC CCT GCA CTG TTG TGG GCG ATG TGC GCT CTG - 3’

For PCR #1, the thermocycling parameters were 98°C for 2 minutes, 24 cycles of (98°C for 10 seconds, 65°C for 30 seconds, 72°C for 45 seconds), and 72°C for 10 minutes. Following PCR #1, reactions for each condition were pooled and 10 µL of each reaction was run on an agarose gel to confirm a 287 bp product. Isopropanol (Pharmco, 231000099) and 3M sodium acetate (Sigma-Aldrich, S2889) pH 5.2 were added to each sample, and samples were precipitated at -20°C for 1 hour. Samples were spun at 10,000 rpm for 30 minutes and the pellet was washed 2 times with fresh 70% ethanol. Samples were air dried for 10-15 minutes and then resuspended in 50-100 µL of TE buffer. Pellets were dissolved at 55°C for 1 hour and mixed by pipetting a few times that hour.

PCRs #2 and #3 were performed to attach Illumina adaptors and barcodes to samples. For PCR #2, 500 ng of DNA per sample of the PCR #1 product was added to each reaction. PCR #2 included primers of variable sequence length to increase library complexity:

Forward primers were an equimolar cocktail of the following staggered primers: KMN_stagger_PCR2_F01: 5’ -

ACACTCTTTCCCTACACGACGCTCTTCCGATCTTCTTGTGGAAAGGACGAACACCG - 3’

KMN_stagger_PCR2_F02: 5’ -

ACACTCTTTCCCTACACGACGCTCTTCCGATCTcTCTTGTGGAAAGGACGAACACCG - 3’

KMN_stagger_PCR2_F03: 5’ -

ACACTCTTTCCCTACACGACGCTCTTCCGATCTagTCTTGTGGAAAGGACGAACACCG - 3’

KMN_stagger_PCR2_F04: 5’ -

ACACTCTTTCCCTACACGACGCTCTTCCGATCTgagTCTTGTGGAAAGGACGAACACCG - 3’

KMN_stagger_PCR2_F05: 5’ -

ACACTCTTTCCCTACACGACGCTCTTCCGATCTcgagTCTTGTGGAAAGGACGAACACCG - 3’

KMN_stagger_PCR2_F06: 5’ -

ACACTCTTTCCCTACACGACGCTCTTCCGATCTtcgacTCTTGTGGAAAGGACGAACACCG - 3’

KMN_stagger_PCR2_F07: 5’ -

ACACTCTTTCCCTACACGACGCTCTTCCGATCTatcaacTCTTGTGGAAAGGACGAACACCG - 3’

KMN_stagger_PCR2_F08: 5’ -

ACACTCTTTCCCTACACGACGCTCTTCCGATCTgaacgaaTCTTGTGGAAAGGACGAACACCG - 3’

Reverse primer (reverse): 5’ - GTGACTGGAGTTCAGACGTGTGCTCTTCCGATCTTCTACTATTCTTTCCCCTGCACTGT - 3’

For PCR #2, the thermocycling parameters were 98°C for 30 seconds, 6 cycles of (98°C for 10 seconds, 55°C for 30 seconds, 72°C for 45 seconds) and 72°C for 10 minutes. Following PCR #2, 3-5 µL of each reaction was run on an agarose gel to confirm a 302 bp product. For PCR #3, 2 µL per sample of the PCR #2 product was added to each reaction. PCR #3 primer sequences were as follows:

KMN_LCV2_PCR3F (forward): 5’ - AATGATACGGCGACCACCGAGATCTACACTCTTTCCCTACACGACGCTCTTCCGATCT - 3’

P7-indexing primer (reverse, xxxxxxxx denotes 8 nucleotide barcode): 5’ – CAAGCAGAAGACGGCATACGAGATxxxxxxxxGTGACTGGAGTTCAGACGTGT - 3’

For PCR #3, the thermocycling parameters were 98°C for 30 seconds, 6 cycles of (98°C for 10 seconds, 55°C for 30 seconds, 72°C for 45 seconds) and 72°C for 10 min. Following PCR #3, 5 µL of each reaction was run on an agarose gel to confirm a 358 bp product. Samples were mixed proportionally, and the mixed sample was run on an agarose gel. DNA was isolated using the QIAquick Gel Extraction Kit (Qiagen, 28704). DNA was sequenced by next-generation sequencing at the Biopolymers Facility at Harvard Medical School (NextSeq 500) in two separate runs with 400 million plus reads per run and single index reads. Sequencing data was pooled for analysis.

#### MAGeCK analysis of screen data

Next-generation sequencing data was processed for MAGeCK analysis. Reads were trimmed, aligned to the reference genome and counted. Read count tables were analyzed by MAGeCK^40^.

### Custom CRISPR/Cas9 minipool screen

SUM159, MDA-MB-468 and BT20 cells were transduced at an MOI of ∼0.3 with a CRISPR/Cas9 lentivirus (Genetic Perturbation Platform, Broad Institute) containing 3,011 sgRNAs, with 10-13 sgRNAs per gene and at least 1000-fold representation. This library also contained 50 sgRNAs targeting 5 essential genes (13 sgRNAs/gene) and 400 negative controls, including 100 no site and 300 intergenic sgRNAs. Cells were selected with 2 µg/mL puromycin for 3 days. After selection, day 0 cell pellets were collected, and the remaining infected cells were expanded for 4 days and then plated into treatment arms in tissue culture-treated 15-cm plates (SUM159: 94,375 cells/mL, MDA-MB-468: 377,500 cells/mL, BT20: 566,250 cells/mL). Cells were treated for 72 hours with DMSO (vehicle) or GR50 doses of the PI3Kα-selective inhibitor BYL719 (SUM159: 2.28 µM, MDA-MB-468: 13.9 µM, BT20: 2 µM) or the catalytic AKT inhibitor, GDC-0068 (SUM159: 4.34 µM, MDA-MB-468: 8.49 µM, BT20: 0.9 µM). At the end point, cell pellets were harvested, and genomic DNA was extracted using the Nucleospin Blood L Midi genomic DNA extraction kit (Macherey-Nagel, 740954.20). Genomic DNA was PCR amplified by the Genetic Perturbation Platform at the Broad Institute using the following primers:

P5 ARGON (forward): 5’ - TTGTGGAAAGGACGAAACACCG - 3’

P7 BEAKER (reverse): 5’ - CCAATTCCCACTCCTTTCAAGACCT - 3’

The thermocycling parameters were 95°C for 5 minutes, 28 cycles of (95°C for 30 seconds, 53°C for 30 seconds, 72°C for 20 seconds) and 72°C for 10 minutes. Next-generation sequencing was also performed by the Genetic Perturbation Platform at the Broad Institute (Hiseq 2500/50 cycles) with approximately 150 million reads per sequencing lane. Data analysis was performed using the Broad Institute’s CRISPR Gene Scoring Tool.

### Cell density (Sulforhodamine B) assays

Cells were seeded in tissue culture-treated 96-well plates in 80-100 µL of appropriate growth media (SUM159: 1000-2000 cells/well, MDA-MB-468: 4000 cells/well, BT20: 6000 cells/well, T47D: 6000 cells/well, MCF7: 4000 cells/well, BT474: 6000 cells/well, HCC70: 6000 cells/well, MCF10A: 2000 cells/well, HepG2: 4000 cells/well, BT549: 4000 cells/well, HCC1937: 4000 cells/well, ZR-75-1: 4000 cells/well, MDA-MB-361: 7000 cells/well, parental T47D: 6000 cells/well, Fulvestrant-resistant T47D clones 1-3: 6000 cells/well). Cells were incubated for 24 hours, and cell density was assayed using sulforhodamine B (SRB; Sigma-Aldrich, 230162) staining, as previously described^41^, to determine the number of cells at the start of the experiment (Day 0). Cells were then treated with 5-20 µL of drug for indicated periods of time to bring the final volume in each well to 100 µL. At the endpoint, relative cell density was assayed using SRB staining. Cell density at each time point was normalized to the day 0 control. For dose curve and double dose curve experiments, these values were normalized from 0-100 using GraphPad PRISM, where an empty well (background) served as the 0% reference, and untreated cells served as the 100% reference. Normalized cell densities were plotted versus log10 drug concentration, and a nonlinear curve was fit using the log(inhibitor) vs. normalized response -- Variable slope function in GraphPad PRISM. IC50 values were calculated by GraphPad PRISM based on the nonlinear curve fit. Cell density data for experiments that were not dose curves or double dose curves were transformed to log2(Y) values using GraphPad PRISM and plotted such that values less than 0 indicate loss of cells or cell death.

### GR50 calculator

GR50 values were calculated for the CRISPR/Cas9 minipool screen using the online GR calculator (http://www.grcalculator.org/grcalculator/). Cells were seeded, and cell density was measured as described for the cell density assays. Cell densities at day 0 (24 hours after seeding) and 72 hours post-drug treatment were used to calculate GR50 values.

### Synergy calculations

For proliferation assays with two inhibitors, synergy scores were calculated using the synergyfinder R package^42^. HSA synergy scores were reported for each drug dose combination tested and displayed as a heatmap.

### Incucyte cell death assays

Cells were seeded in black-walled, tissue culture-treated 96-well plates as described for the cell density assays. 24 hours later, the media was changed to 90 µL of growth media containing 1:1000 Incucyte Nuclight Rapid Red Dye (Sartorius, 4717) and 1:1000 NucView 488 Caspase-3/7 substrate (Biotium, 30029). Cells were treated with 10 µL of drug-containing media. Plates were then placed in the Incucyte instrument, and images were taken every 2 hours for 72 hours. The Incucyte software was used to train a model to count cells as those expressing the nuclear dye, Incucyte Nuclight Rapid Red Dye. A model was also trained to count green cells, or those with cleaved caspase-3/7 signal. Four images were taken per well and averaged, and quadruplicate wells were assayed for each condition. Percentage of cell death was measured as the number of cells with an overlapping signal (red and green) divided by the number of red cells.

### Mouse studies

All animal experiments were performed at Beth Israel Deaconess Medical Center (BIDMC) in collaboration with the laboratory of J.G.C. and in accordance with the guidelines of the BIDMC Institutional Animal Care and Use Committee (IACUC). HCC70 cells were maintained in RPMI-1640 supplemented with 10% FBS, and cells tested negative for mycoplasma before injection. On the day of injection, HCC70 cells were washed twice with 1X PBS, trypsinized and counted. A total of 5 x 10^6^ cells per mouse were resuspended in 100 µL of serum-free RPMI-1640 and placed on wet ice. Cells were mixed with Matrigel (Corning, 356230) in a 1:1 (v/v) ratio and injected subcutaneously into the flanks of 50 NSG mice.

Tumors were allowed to grow for 18 days before switching mice from standard chow to a low geranylgeraniol diet for 3 days (Research Diets, D22092101l – modified open standard diet with 15 kcal % fat canola oil). 10-14 mice were then assigned to each treatment group (vehicle, AZD5363, pitavastatin, combination), and treatments were administered daily by oral gavage. AZD5363 (AstraZeneca) was prepared as a suspension in 0.5% carboxymethylcellulose and was dosed daily for 4 days of the week at 100 mg/kg followed by a 3-day AKT inhibitor holiday. Pitavastatin (Selleckchem, S1759) was prepared by diluting 100 mM pitavastatin in DMSO in 0.5% carboxymethylcellulose and was dosed daily at 100 mg/kg. Tumors were measured with calipers 2 times per week (length and width) for 35 days (24 days after the start of treatments). After 24 days of treatment, all mice were euthanized. Four mice in each group were treated with AZD5363 (2 hours) or pitavastatin (6 hours) prior to euthanasia and sections of tumor and liver were snap frozen and fixed in 10% formalin for immunoblotting and immunohistochemistry, respectively.

Tissues were homogenized in cold RIPA lysis buffer (150 mM Tris-HCl, 150 mM NaCl, 0.5% (w/v) sodium deoxycholate (Sigma-Aldrich, D6750), 1% (v/v) NP-40 (Sigma-Aldrich, I3021), pH 7.5) containing 0.2% SDS, 2 mM Na_4_P_2_O_7_, 10 mM NaF, 0.5% (v/v) protease inhibitor cocktail, 100 nM calyculin A, 4 mM sodium orthovanadate (Sigma-Aldrich, S6508) and 2X HALT protease and phosphatase inhibitor cocktail (Fisher Scientific, PI78443), added just before use. Tissue homogenization was facilitated by use of the gentleMACS dissociator (Miltenyi Biotec, 130-093-235). Samples were then centrifuged at 4,000 x g for 5 minutes at 4°C. The supernatants were transferred to new tubes, and samples were spun at maximum speed for 15 minutes at 4°C. The supernatants were again transferred to new tubes, and this process of centrifugation and supernatant collection was repeated. Once supernatants were no longer cloudy, protein concentrations were measured, and immunoblotting was performed as described in the immunoblotting methods section.

### Patient-derived organoid cultures

Propagation and culturing of patient-derived organoid cultures (PDOs) was previously described^43^. Briefly, PDOs were incubated in 1× Dispase-II solution with 2 mg/mL collagenase for 30-45 minutes at 37°C and mechanically disrupted by passing through a 26G needle. PDOs were washed once with Advanced DMEM/F12 supplemented with 5% FBS and pelleted by centrifugation at 400 x g for 5 minutes. PDO fragments were embedded in Cultrex growth factor-reduced basement membrane extract type II (Trevigen, 3533-001-02), 50 µL drops were plated into a 24-well plate, and 500 µL PDO media was added 30 minutes later^43^. To assess drug sensitivity, 200-600 PDO fragments were plated into 8-well chamber slides, and 1 μM AZD5363 (Cayman Chemicals, 15406) and/or 5 μM pitavastatin (Selleckchem, S1759) was added the following day. After 96 hours of drug treatment, PDOs were pulsed with 10 μM 5-ethynyl-2′-deoxyuridine (EdU) for 4 hours and fixed with 4% paraformaldehyde for 30 minutes. CellProfiler software was used to measure PDO area for 10-20 PDOs at the endpoint.

To assess cell proliferation and apoptosis, fixed PDOs were permeabilized with wash buffer (0.3% Triton X-100 in PBS) for 20 minutes. EdU labeling was performed for 40 minutes using the EdU Click-IT imaging kit (Invitrogen, C10337) according to manufacturer’s description. PDOs were washed 3 times with wash buffer, blocked for 1 hour with blocking buffer (5% goat serum, 0.2% BSA, 0.3% Triton X-100 in PBS) and incubated with anti-cleaved caspase-3 (Cell Signaling Technology, 9661) in blocking buffer overnight at 4°C. The following day the PDOs were washed extensively with wash buffer, incubated with secondary antibody (Alexa Fluor 488) for 2 hours at room temperature, washed with wash buffer and mounted using Vectashield mounting media containing DAPI (Vector Laboratories, H-1200-10). PDOs were imaged with Zeiss LSM 880 confocal microscope. To assess proliferation, 10-20 PDOs per treatment condition were imaged and the ratio of EdU positive cells per total number of cells was quantified. To assess apoptosis, 10-20 PDOs per treatment condition were scored based on the presence of cleaved caspase-3 staining.

To assess signaling changes in response to drug treatments, PDOs were incubated with1 μM AZD5363 (Cayman Chemicals, 15406) and/or 5 μM Pitavastatin (Selleckchem, S1759) for 24 hours. PDOs were pooled from 6 wells and incubated for 1.5 hours on ice in Cell Recovery Buffer supplemented with phosphatase inhibitors (Roche, 4906845001). PDOs were lysed using RIPA buffer (Boston BioProducts, BP-115) containing protease and phosphatase inhibitors (Thermo Scientific, A32959) and then centrifuged at 13,000 rpm for 10 minutes at 4°C. Protein concentrations were measured using Pierce BCA protein assay kit. Sample concentrations were normalized using 6X SDS sample buffer (62.5 mM Tris pH 6.8, 5% SDS, 18% glycerol (Fisher Scientific G33-500), bromophenol blue (Fisher Scientific, BP115-25), 302 mM dithiothreitol (DTT) (Fisher Scientific, BP172-25) and RIPA lysis buffer). Cell lysates were boiled at 95°C for 5 minutes and stored at -20°C. For detection of HMGCR, cell lysates were not boiled or frozen, but were processed and run on the same day to avoid temperature changes that cause endoplasmic reticulum-associated protein aggregation. Immunoblotting was performed as described in the immunoblotting methods section.

**Table 1.**
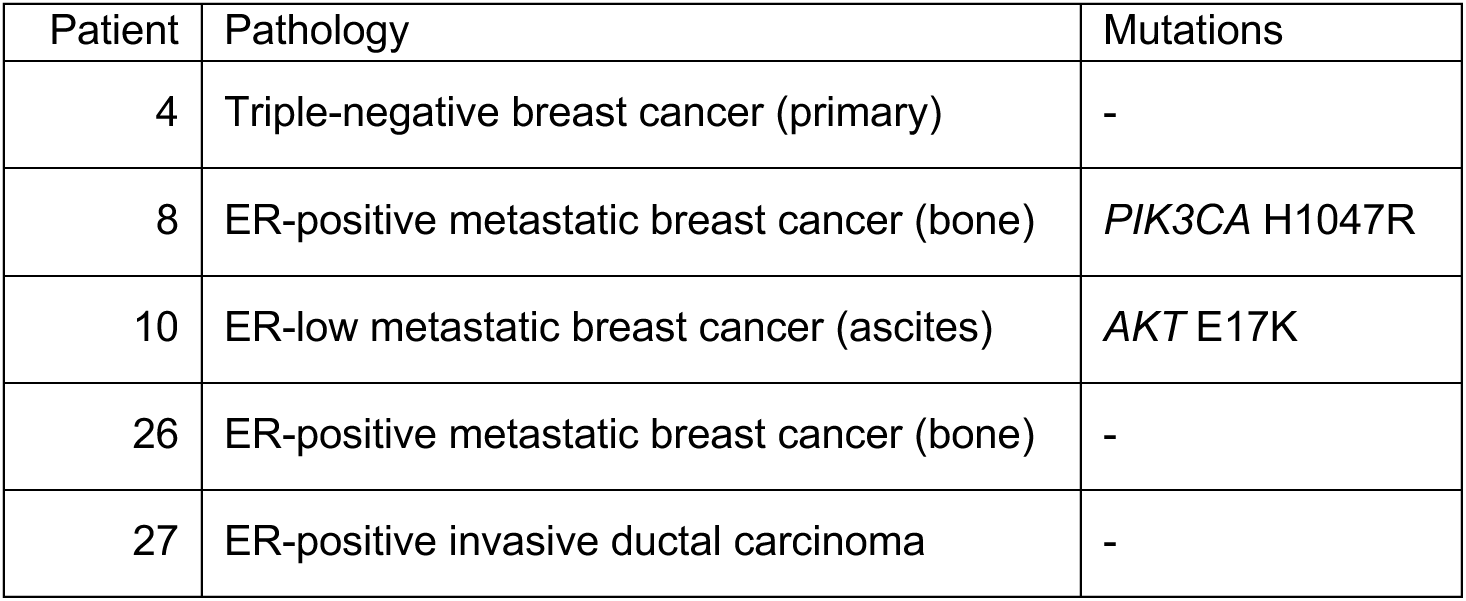
Patient-derived organoid characteristics.

### Immunoblotting

Cells were washed once in cold 1X PBS (Boston BioProducts, BM-220) and collected on wet ice in 4°C RIPA lysis buffer (150 mM Tris-HCl, 150 mM NaCl, 0.5% (w/v) sodium deoxycholate (Sigma-Aldrich, D6750), 1% (v/v) NP-40 (Sigma-Aldrich, I3021), pH 7.5) containing 0.1% sodium dodecyl sulfate (SDS; AmericanBio, AB01920-00500), 1 mM sodium pyrophosphate (Na_4_P_2_O_7_; Sigma-Aldrich, S390-500), 20 mM sodium fluoride (NaF; Fisher Scientific, S25547), 0.5% (v/v) protease inhibitor cocktail (104 mM AEBSF, 80 μM aprotinin, 4 mM bestatin, 1.4 mM E-64, 2 mM leupeptin and 1.5 mM pepstatin A; Sigma-Aldrich, P8340) and 50 nM calyculin A (LC Laboratories, C-3987), added just before use. Plates were scraped into 1.5 mL microcentrifuge tubes, vortexed, and incubated on wet ice for 15 minutes. Samples were then centrifuged at 14,000 rpm for 10 minutes at 4°C. The supernatants were transferred to new tubes, and protein concentrations were measured by the Bio-Rad DC protein assay (Bio-Rad, Reagent A: 5000113, Reagent B: 500-0114). Sample concentrations were normalized using 2X SDS sample buffer (62.5 mM Tris pH 6.8, 2% SDS, 10% glycerol (Fisher Scientific G33-500), bromophenol blue (Fisher Scientific, BP115-25), 5% (v/v) β-mercaptoethanol (Sigma-Aldrich, M3148)). Cell lysates were boiled at 95°C for 5 minutes and stored at -20°C. For detection of HMGCR, cell lysates were not boiled or frozen, but were processed and run on the same day to avoid temperature changes that cause endoplasmic reticulum-associated protein aggregation. Cell lysates were run by SDS-PAGE on 7.6-15% acrylamide gels in 1X running buffer (Boston BioProducts, BP-150). Proteins were transferred to nitrocellulose membranes at 100 volts for 90 minutes in cold 1X transfer buffer (Cell Signaling Technology, 12539S). Membranes were blocked with 5% (w/v) bovine serum albumin (BSA; Gold Biotechnology, A-420-100) in tris-buffered saline (TBS; Boston BioProducts, BM-301), rocking for at least 1 hour at room temperature. Membranes were washed briefly in TBS-tween (TBST; Boston Bioproducts, IBB-181X) and incubated in primary antibodies diluted in 5% (w/v) BSA in TBST with 0.01% (w/v) sodium azide (Fisher Scientific, S227I-25), rocking at 4°C overnight. Membranes were washed three times for 10 minutes in TBST, rocking at room temperature. Then, membranes were incubated for 1 hour rocking at room temperature with fluorophore-conjugated secondary antibodies (LI-COR Biosciences). Membranes were washed two times for 10 minutes each in TBST and one time for 10 minutes in TBS and imaged with the LI-COR Odyssey CLx or Odyssey M Imaging System (LI-COR Biosciences).

**Table 2.** Antibodies for immunoblotting.

|  | Antibody | Manufacturer | Identifier | Host | Dilution |
| --- | --- | --- | --- | --- | --- |
| Primary antibody | PARP | Cell Signaling Technology | 9542 | Rabbit | 1:1000 |
|  | pAKT S473 | Cell Signaling Technology | 4060 | Rabbit | 1:1000 |
|  | pAKT T308 | Cell Signaling Technology | 2965 | Rabbit | 1:1000 |
|  | AKT | Cell Signaling Technology | 4691 | Rabbit | 1:1000 |
|  | pPRAS40 T246 | Cell Signaling Technology | 2997 | Rabbit | 1:1000 |
|  | PRAS40 | Cell Signaling Technology | 2691 | Rabbit | 1:1000 |
| | pGSK3β S9 | Cell Signaling Technology | 9336 | Rabbit | 1:1000 |
| | GSK3β | Cell Signaling Technology | 9315 | Rabbit | 1:1000 |
|  | pS6 | Cell Signaling Technology | 5364 | Rabbit | 1:1000 |
|  | S6 | Cell Signaling Technology | 2217 | Rabbit | 1:1000 |
|  | pP70S6K | Cell Signaling Technology | 9234 | Rabbit | 1:1000 |
|  | P70S6K | Cell Signaling Technology | 2708 | Rabbit | 1:1000 |
|  | p4E-BP1 | Cell Signaling Technology | 2855 | Rabbit | 1:1000 |
|  | 4E-BP1 | Cell Signaling Technology | 9452 | Rabbit | 1:1000 |
|  | Unprenylated RAP1A | Santa Cruz Biotechnology | sc-373968 | Mouse | 1:50 |
|  | RHEB | Abnova | H00006009-M01 | Mouse | 1:50 |
|  | HDJ2 | ThermoFisher Scientifici | MA5-12748 | Mouse | 1:1000 |
|  | PARP | Cell Signaling Technology | 9542 | Rabbit | 1:1000 |
|  | Cleaved caspase-3 | Cell Signaling Technology | 9664 | Rabbit | 1:1000 |
|  | NPC1 | Cell Signaling Technology | 33422 | Rabbit | 1:1000 |
|  | HMGCR | Invitrogen | MA5-31335 | Mouse | 1:600 |
|  | SREBP-1 | Abcam | ab28481 | Mouse | 1:500 |
|  | SREBP-2 | BD Biosciences | 557037 | Mouse | 1:250 |
| | Estrogen receptor $\alpha$ (ERα) | Cell Signaling Technology | 8644 | Rabbit | 1:1000 |
|  | c-Myc | Cell Signaling Technology | 5605 | Rabbit | 1:1000 |
| | β-catenin | Cell Signaling Technology | 8480 | Rabbit | 1:1000 |
|  | LC3-I/II | Cell Signaling Technology | 4108 | Rabbit | 1:1000 |
|  | BiP | Cell Signaling Technology | 3183 | Rabbit | 1:1000 |
|  | ATF-4 | Cell Signaling Technology | 11815 | Rabbit | 1:1000 |
| | β-actin | Cell Signaling Technology | 4970 | Rabbit | 1:3000 |
|  | Vinculin | Cell Signaling Technology | 13901 | Rabbit | 1:1000 |
| Secondary antibody | IRDye® 800CW Goat anti-Rabbit IgG (H + L) | LI-COR | 926-32211 | Goat | 1:20,000 |
|  | IRDye® 800CW Goat anti-Mouse IgG (H + L) | LI-COR | 926-32210 | Goat | 1:20,000 |
|  | IRDye® 680CW Goat anti-Rabbit IgG (H + L) | LI-COR | 926-68021 | Goat | 1:20,000 |
|  | IRDye® 680CW Goat anti-Mouse IgG (H + L) | LI-COR | 926-68070 | Goat | 1:20,000 |

### Sterol and non-sterol rescue experiments

Cells were seeded as described for cell proliferation assays. Cells were incubated for 24 hours, and cell density was assayed using SRB staining to determine the number of cells at the start of the experiment (day 0). Cells were then treated with 5 µL of drug and 5 µL of vehicle (7:3 MeOH:NH_4_OH) or sterol and non-sterol intermediates for 72 hours to bring the final volume in each well to 100 µL: 1 mM mevalonate (Sigma-Aldrich, 90469), 5 µg/mL cholesterol (Sigma-Aldrich, C4951) or 5 µg/mL GGPP (Cayman Chemicals, 63330). At the endpoint, relative cell density was assayed using SRB staining. Cell density at 72 hours was normalized to the day 0 control. Normalized cell densities were log2-transformed and plotted in GraphPad PRISM such that values greater than 0 indicate increased cell density, while values less than 0 indicate decreased cell density at 72 hours compared to the day 0 control.

### RNA-sequencing

Cells were plated at 250,000-500,000 cells/mL in RPMI-1640 medium supplemented with 10% FBS at 2 mL per well in tissue culture-treated 6-well plates to achieve 75% density by the endpoint. The next day, stocks of AZD5363 (Cayman Chemicals, 15406) and pitavastatin (Selleckchem, S1759) were prepared in DMSO, and cells were treated for 24 or 48 hours. A master mix of each drug stock was prepared and used to treat all conditions across biological replicates, stored at -80°C between treatments. After 24 or 48 hours of treatment, each well was washed once with 1 mL of cold 1X PBS and aspirated completely. Plates were snap frozen on dry ice and stored at -80°C until all biological replicates were collected. Four biological replicates were seeded on sequential days at the same time of day. Samples were collected for protein harvest in parallel to confirm expected drug effects on cell signaling. Snap frozen plates were thawed on ice, and RNA was extracted using Takara’s Nucleospin RNA Plus kit (Takara, 740984.50). RNA quantity and purity were assessed by Nanodrop 1000. Samples were submitted to Novogene for integrity assessment (Agilent 2100 analysis), mRNA library preparation (unstranded) and paired-end (150 bp) sequencing on a NovaSeq6000, S4 flow cell. Data was analyzed as described previously^44^.

### DAVID Gene ontology analysis

Genes that were uniquely up- or downregulated with AZD5363, pitavastatin or combination treatment in MDA-MB-468 or T47D cells in the RNA-sequencing data were analyzed using the DAVID 2021 Bioinformatics Resource Functional Annotation tool. Differentially regulated genes were uploaded as lists with the background genome set to all genes detected in the RNA-sequencing analysis of that cell line, and biological processes were plotted by -log10(p-value) where the p-value is an EASE Score, a modified Fisher Exact p-value for gene enrichment analysis^45,46^.

### siRNA Transfections

Cells were seeded in 96-well plates in 90 µL of appropriate growth media (SUM159: 1000-2000 cells/well, MDA-MB-468: 4000 cells/well, BT20: 6000 cells/well, T47D: 6000 cells/well, MCF7: 4000 cells/well, BT474: 6000 cells/well). Cells were incubated for 24 hours before transfection. Cells were transfected with Lipofectamine RNAiMAX transfection reagent (ThermoFisher Scientific, 13778150) as follows. All reagents were brought to room temperature before proceeding with the transfection. Lipofectamine RNAiMAX was diluted in Opti-MEM reduced serum media (ThermoFisher Scientific, 31985070) with 1.5 µL of Lipofectamine RNAiMAX for every 25 µL of Opti-MEM media, vortexed and incubated at room temperature for 5 minutes. siRNAs were diluted to 500 nM in Opti-MEM media, vortexed and incubated at room temperature for 5 minutes. Equal volumes of Lipofectamine RNAiMAX in Opti-MEM media and siRNA in Opti-MEM media were combined, vortexed and incubated for 20 minutes at room temperature. For a lipofectamine only control, equal volumes of Lipofectamine RNAiMAX in Opti-MEM and Opti-MEM media were combined, vortexed and incubated at room temperature for 20 minutes. 10 µL of the Lipofectamine RNAiMAX, siRNA, Opti-MEM mixtures were then added to appropriate wells to bring the final volume in each well to 100 µL and the final concentration of siRNA to 25 nM. After 24 hours, media was changed to 90 µL of fresh media using gentle pipetting to avoid disrupting cells, and cell density of lipofectamine only control cells was assayed using SRB staining to determine the number of cells at the start of the drug treatment (day 0). Cell density at 72 hours was normalized to the cell density at the start of drug treatment (day 0). Normalized cell densities were log2-transformed and plotted in GraphPad PRISM such that values greater than 0 indicate increased cell density, while values less than 0 indicate decreased cell density at 72 hours compared to the day 0 control.

**Table 3.**
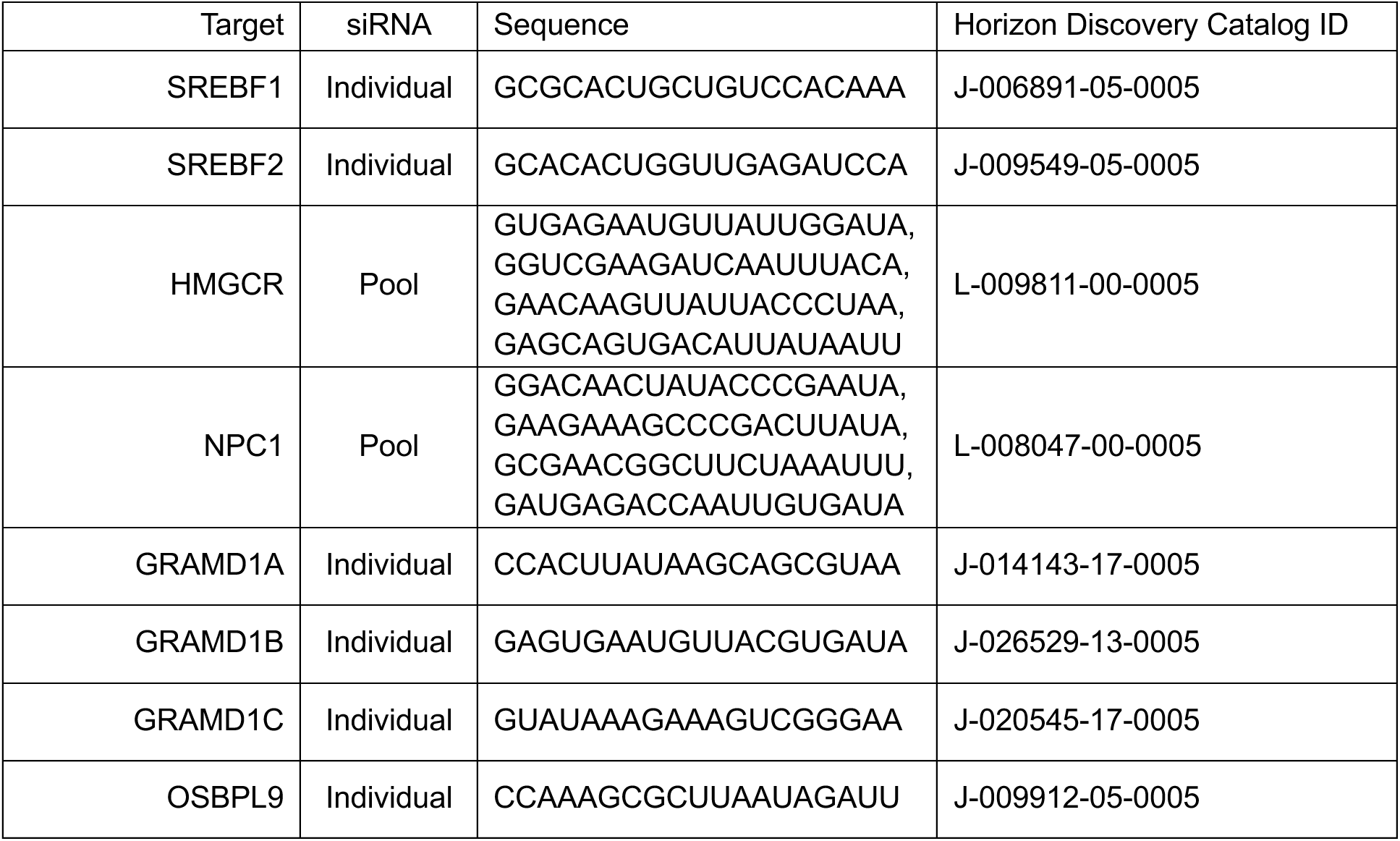
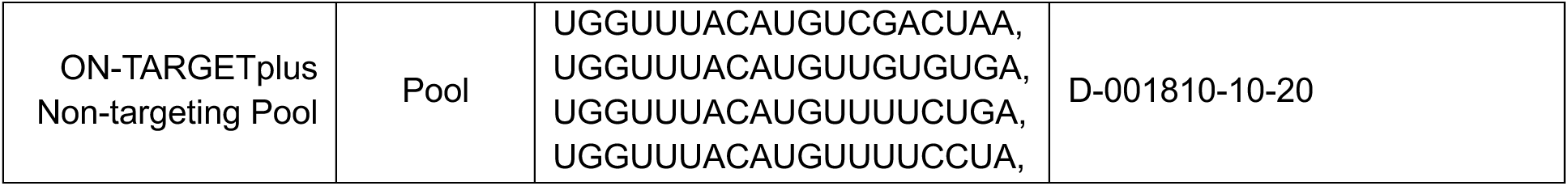
siRNA Product information.

### Real-time quantitative polymerase chain reaction (RT-qPCR)

RNA was isolated from cells with the NucleoSpin RNA Plus Kit (Macherey-Nagel, 740984) according to the manufacturer’s protocol. Reverse transcription was performed using the TaqMan Reverse Transcription kit (ThermoFisher Scientific, N8080234). cDNA was detected using PowerUp SYBR Green Master Mix for qPCR (ThermoFisher Scientific, A25776). A master mix of 250 µM forward primer, 250 µM reverse primer, 1X SYBR Green Master Mix, and nuclease-free water up to 10 µL per reaction was prepared. 10 µL of master mix was added to each well of a 384-well plate, and 2 µL of 2.5 ng/µL cDNA was added to each well for 5 ng of cDNA per reaction. The plate was spun briefly at 1000 rpm. RT-qPCR was performed using a CFX384 Touch Real-Time PCR Detection System (Bio-Rad). The thermocycling parameters were 50°C for 2 minutes, 95°C for 2 minutes, 40 cycles of (95°C for 15 seconds, 60°C for 1 minute), 65°C for 5 seconds and 95°C for 5 seconds. RT-qPCR was performed in technical triplicate, and quantification of mRNA expression was calculated by the 1′1′CT method with 18S ribosomal RNA as the reference gene. See Table 4 for RT-qPCR primers, which were designed using the NCBI Primer-BLAST tool. Primer efficiency was confirmed to be within 90-110% for each primer in each cell line.

**Table 4.**
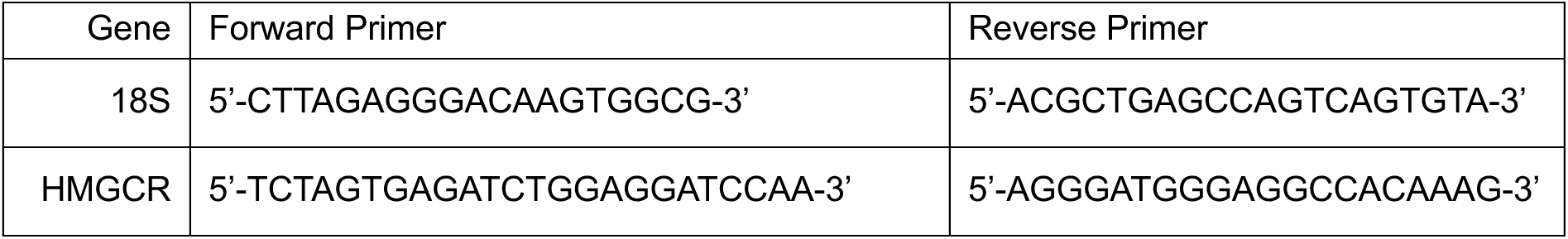
RT-qPCR primers.

### Plasmids

See Table 5 for detailed plasmid information. pHAGE-ESR1 was purchased from Addgene (116737). To generate pLenti6/V5-HMGCR, HMGCR was PCR-amplified out of pCMV-SPORT6-hHMGCR1 (Addgene, 86085) using the forward primer 5’- GTATACTGGATCCGCCGCCACCATGTTGTCAAGACTT-3’ and reverse primer 5’-TCGGAGCTCGAGGTGGCTGTCTTCTTGGT-3’. PCR-amplified HMGCR was cloned into pLenti6/V5-p53_wt p53 (gift from the Muranen lab) by double restriction enzyme digestion with BamHI (New England Biolabs, R0136) and XhoI (New England Biolabs, R0146) at 37°C for 1 hour, followed by ligation with T4 DNA ligase (New England Biolabs, M0202) following the manufacturer’s protocol. To generate mutations in the catalytic residues of HMGCR, four sequential site-directed mutagenesis reactions were performed following the QuikChange II XL Site-Directed Mutagenesis Kit protocol (Agilent, 200521). The following primers were generated using Agilent’s QuikChange Primer Design tool: H866A: 5’- CAGGACATCTTGTCAAAAGTGCCATGATTCACAACAGGTCGA-3’ (forward), 5’- TCGACCTGTTGTGAATCATGGCACTTTTGACAAGATGTCCTG-3’ (reverse), E559A: 5’- CCAATGGCAACAACAGCAGGTTGTCTTGTGGCC-3’ (forward), 5’- GGCCACAAGACAACCTGCTGTTGTTGCCATTGG-3’ (reverse), K691A: 5’- GTTAGTGGTAACTATTGTACTGACGCGAAACCTGCTGCTATAAATTGGAT-3’ (forward), 5’- ATCCAATTTATAGCAGCAGGTTTCGCGTCAGTACAATAGTTACCACTAAC-3’ (reverse), D767A: 5’- CATTGCCTGTGGACAGGCTGCAGCACAGAATGTTG-3’ (forward), 5’- CAACATTCTGTGCTGCAGCCTGTCCACAGGCAATG-3’ (reverse). To generate pLenti6-ER-mRFP, mRFP with a C-terminal KDEL sequence (ER-mRFP) was PCR amplified out of ER-mRFP (Addgene, 62236) using the forward primer 5’-GGTCGGATCCGCCGCCACCATGGACAGCAAAGG-3’ and the reverse primer 5’-GAGGCACCGGTGTTTAGAGCTCATCTTT-3’. PCR-amplified ER-mRFP was cloned into pLenti6/V5-p53_wt p53 (gift from the Muranen lab) by double restriction enzyme digestion with BamHI-HF (New England Biolabs, R0136) and AgeI-HF (New England Biolabs, R3552) at 37°C for 1 hour, followed by ligation with T4 DNA ligase (New England Biolabs, M0202) following the manufacturer’s protocol.

**Table 5.**
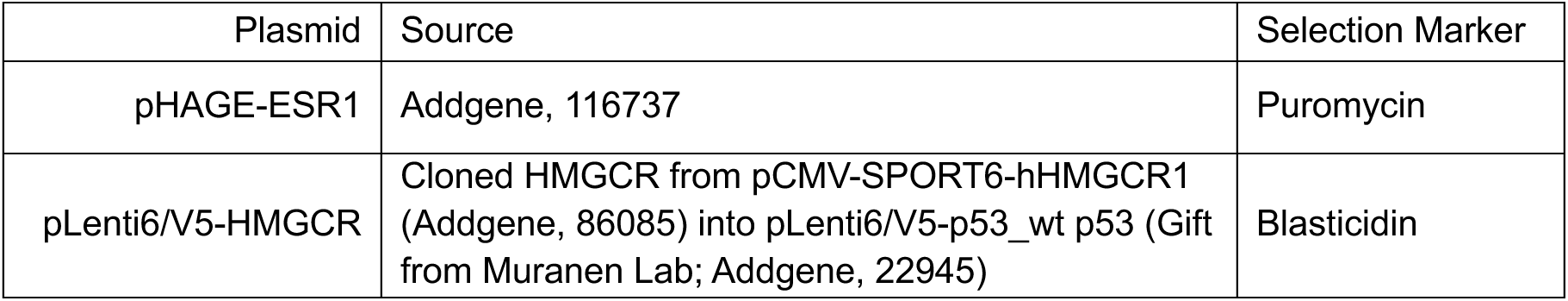
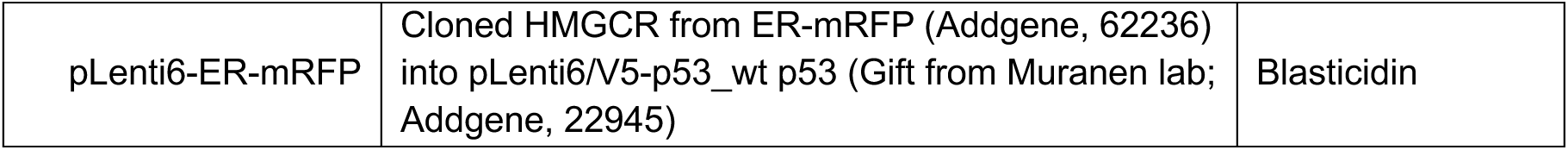
Plasmids.

| Plasmid | Source | Selection Marker |
| --- | --- | --- |
| pHAGE-ESR1 | Addgene, 116737 | Puromycin |
| pLenti6/V5-HMGCR | Cloned HMGCR from pCMV-SPORT6-hHMGCR1 (Addgene, 86085) into pLenti6/V5-p53_wt p53 (Gift from Muranen Lab; Addgene, 22945) | Blasticidin |
| pLenti6-ER-mRFP | Cloned HMGCR from ER-mRFP (Addgene, 62236) into pLenti6/V5-p53_wt p53 (Gift from Muranen lab; Addgene, 22945) | Blasticidin |

### Transfection and lentiviral transduction of plasmid DNA

HEK 293T cells were transfected with pHAGE-ESR1, pLenti6/V5-HMGCR or pLenti6-ER-mRFP plasmid DNA as follows. A master mix of 11.1 µg psPAX2 (Gal/Pol), 0.6 µg VSVG, 6.3 µg of plasmid DNA and 1200 µL of serum-free, antibiotic-free DMEM was prepared. 54 µL of 1 mg/mL polyethylenamine (PEI; Sigma-Aldrich, 408727) was added, and the master mix was vortexed for 10 seconds to mix and then incubated at room temperature for 15 minutes. Tissue culture-treated 10-cm dishes were coated with poly-L-lysine (PLL; Sigma-Aldrich, P1274) for 5 minutes and then washed three times with 1X PBS. HEK 293T cells were trypsinized and counted, and 9 x 10^6^ cells were resuspended in 7.8 mL of DMEM supplemented with 10% FBS. The approximately 1200 µL reaction mixture was added to the HEK 293T cell suspension, mixed and plated on the PLL-coated 10-cm plates. GlutaMAX supplement (ThermoFisher, 35050061) was added to a final concentration of 1X. The cells were incubated for 48 hours. 24 hours after transfection of HEK 293T cells, target cell lines were seeded in 6-well plates at 62,500-250,000 cells/mL in complete media (RPMI-1640 supplemented with 10% FBS). 48 hours after transfection of HEK 293T cells, virus was harvested. Media from the HEK 293T cells was passed through a 10 mL syringe with a 0.45 µm filter attached. Media was aspirated from the target cells, and virus was added to each well (pHAGE-ESR1: 125 µL). For pLenti6/V5-HMGCR and pLenti-ER-mRFP, virus was concentrated following the PEG-it virus precipitation protocol (LV810A-1). Viral particles were resuspended in 80 µL of PBS, and 30 µL (pLenti6/V5-HMGCR) or 10 µL (pLenti-ER-mRFP) was added to each well. The volume in each well was brought up to 1 mL with RPMI-1640 supplemented with 10% FBS. Polybrene (Sigma-Aldrich, 107689) was added at a final concentration of 5 µg/mL, and cells were incubated at 37°C for 4 hours. After 4 hours, 3 mL of media was added to each well, and cells were incubated at 37°C overnight. The next day, media was changed on the transduced cells to antibiotic containing media (1 µg/mL puromycin, 6 µg/mL blasticidin). Cells were maintained and expanded in antibiotic-containing media.

### Endoplasmic reticulum cholesterol depletion

Cells cultured in RPMI supplemented with 10% FBS were trypsinized and seeded into 90 µL of RPMI supplemented with 10% lipid-depleted FBS (Biowest, S148L) in 96-well plates. Cells were incubated for 24 hours and then treated with 10 µL of DMSO or high dose (10 µM) pitavastatin for 1 hour. After 1 hour, cell density was assayed using SRB staining to determine the number of cells at the start of the experiment (day 0). The media was changed to 100 µL of fresh RPMI supplemented with either 10% lipid-depleted FBS or 10% FBS (complete serum) containing DMSO or low dose (2 µM) pitavastatin for 72 hours. At the endpoint, relative cell density was assayed using SRB staining. Cell density at 72 hours was normalized to the day 0 control. These values were normalized from 0-100 using GraphPad PRISM, where an empty well (background) served as the 0% reference, and untreated cells within the +/- 10 µM pitavastatin pre-treatment groups served as the 100% reference.

### Immunofluorescence

Black-walled, glass bottom, tissue culture-treated plates were coated with poly-L-lysine (PLL; Sigma-Aldrich, P1274) for 5 minutes and then washed three times with 1X PBS. Cells were seeded at 100,000 cells/well in in 80 µL of RPMI supplemented with 10% FBS. Cells were incubated for 24 hours and then treated with 10 µL of DMSO or 1 µM U18666A (Cayman Chemicals, Cholesterol Cell-Based Detection Assay Kit, 10009779) and 10 µL of DMSO or 2 µM pitavastatin (Selleckchem, S1759) for 24 hours. Cells were then fixed with 4% formaldehyde solution (Millipore-Sigma, 1004965000) for 1 hour at room temperature and then washed three times with 1X PBS. Cells were then stained with Filipin III for 1 hour rocking at room temperature in the dark, as indicated by the Cholesterol Cell-Based Detection Assay Kit protocol (Cayman Chemicals, 10009779). After Filipin III staining, the cells were maintained in the dark for the remaining steps. Cells were washed three times with 1X PBS and then blocked with 0.5% bovine serum albumin (BSA; Gold Biotechnology, A-420-100) in 1X PBS for 1 hour rocking at room temperature. After blocking, cells were incubated with primary antibody for 1 hour rocking at room temperature (LAMP1; Cell Signaling Technology, 9091, 1:100). Cells were washed three times with 1X PBS and then incubated with secondary antibody for 30 minutes rocking at room temperature (Goat anti-Rabbit IgG Alexa Fluor 488; ThermoFisher Scientific, A-11008). Cells were then washed two times with 1X PBS and imaged on the Keyence BZ-X800 microscope.

**Table 6.** Key reagents.

| Reagent | Supplier | Product number |
| --- | --- | --- |
| BYL719 | Active Biochem | A-1214 |
| GDC-0068 | MedChemExpress | RG7440 |
| AZD5363<br>(capivasertib) | Cayman Chemicals (*in vitro* studies),<br>AstraZeneca (*in vivo* studies) | 15406 |
| ARQ-092 | Cayman Chemicals | 1313881-70-7 |
| MK-2206 | Cayman Chemicals | 11593 |
| Torin 1 | Tocris Bioscience | 4247 |
| Lovastatin | Selleckchem | S2061 |
| Rosuvastatin | Selleckchem | S2169 |
| Pitavastatin | Selleckchem | S1759 |
| Bortezomib | Cayman Chemicals | 10008822 |
| Fulvestrant | Selleckchem | S1191 |
| GGTI-298 | Selleckchem | S7466 |
| FTI-277 | Selleckchem | S7465 |
| Fatostatin | Selleckchem | S9785 |
| Cycloheximide | Cayman Chemicals | 14126 |
| MG132 | Cayman Chemicals | 10012628 |
| Chloroquine | Sigma-Aldrich | C6628 |
| Tunicamycin | Sigma-Aldrich | T7765 |
| U18666A | Cayman Chemicals | Cholesterol Cell-Based Detection Assay Kit,<br>10009779 |
| OSW-1 | MedChemExpress | HY-101213 |
| Mevalonate | Sigma-Aldrich | 90469 |
| GGPP | Cayman Chemicals | 63330 |
| Cholesterol | Sigma-Aldrich | C4951 |
| Filipin III | Cayman Chemicals | Cholesterol Cell-Based Detection Assay Kit,<br>10009779 |
| Puromycin | Corning | 61-385-RA |
| Blasticidin | InvivoGen | ant-bl-1 |

