## Supplementary Data for "Cholesterol biosynthesis inhibition synergizes with AKT inhibitors in triple-negative breast cancer"

**Acknowledgements** This work was supported by the following grants: Ludwig Center at Harvard (A.T., M.B., K.C., S.J.E., T.M.), CA253097 (A.T.), NSF Graduate Research Fellowship (A.L.H.), Harvard University Landry Cancer Biology Fellowship (A.L.H., H.E.M.), NIH P01CA250959 (N.Z., M.B.), Susan G. Komen Foundation grant # CCR18547665 (T.M.), Harvard Stem Cell Institute grant # DP-0194-21-00 (T.M.), Sigrid Juselius Foundation (J.M.H.), Orion Research Foundation (J.M.H.), Maud Kuistila Memorial Foundation (J.M.H.), AACR fellowship (J.M.H.), Cancer Research UK Grand Challenge and the Mark Foundation for Cancer Research to the SPECIFICANCER team (K.C.), R01CA234600 (S.J.E.), Howard Hughes Medical Institute (S.J.E.), Department of Defense Impact Award W81XWH-20-1-0867 (D.A.F), Sir Henry Wellcome Fellowship 220464/A/20/Z (R.R.M.).

We thank Jonah Lee for assisting with mouse experiments and members of the Ludwig Center at Harvard University for constructive comments.

**Author contributions** Conceptualization: A.L.H., A.T.; methodology: A.L.H., T.D.M., H.E.M., N.K., N.S.P., D.E.R., K.C., S.J.E., D.A.F., S.T.B., R.R.M., A.T.; investigation: A.L.H., J.M.H., E.L, J.G.C.; resources: T.D.M., N.Z., N.K., N.S.P, D.E.R., M.B., S.J.E, S.T.B.; writing-original draft: A.L.H., A.T.; writing-review: A.L.H., T.D.M., H.E.M., J.M.H., N.Z., E.L., N.K., D.E.R., M.B., K.C., S.J.E., T.M., D.A.F., S.T.B., J.G.C., R.R.M., A.T.; writing-editing: A.L.H., A.T.; supervision: A.T.

**Competing interests** A.T. is a consultant for NovoNordisk Holdings, Inc. and receives funding support from BioHybrid Solutions; M.B. is a consultant and serves on the SAB for Novartis. M.B. receives sponsored research support from Novartis. M.B. serves on the SAB of FibroGen. M.B. serves on the SAB and holds equity in GV20 Therapeutics. M.B. serves on the SAB of and holds equity in Kronos Bio; K.C. is an advisor at Genentech and serves on the scientific advisory board of Erasca, Inc; S.J.E. is a founder of and holds equity in TScan Therapeutics, MAZE Therapeutics, ImmuneID and Mirimus, serves on the scientific advisory boards of Homology Medicines, ImmuneID, MAZE Therapeutics and T-Scan Therapeutics; S.T.B. is employed by Astra Zeneca, Inc; R.R.M. receives consulting fees from Nested

Therapeutics (Cambridge, U.S.) and serves on the Scientific Advisory Board of CLOVES Syndrome Community.

**Data Availability** Source data and annotated analysis workflows are available on the following OSF project website: https://osf.io/6pw9d/. RNA-sequencing data have been deposited with GEO under series accession number GSE252944. A complete list of reagents is provided in Tables 2-6.

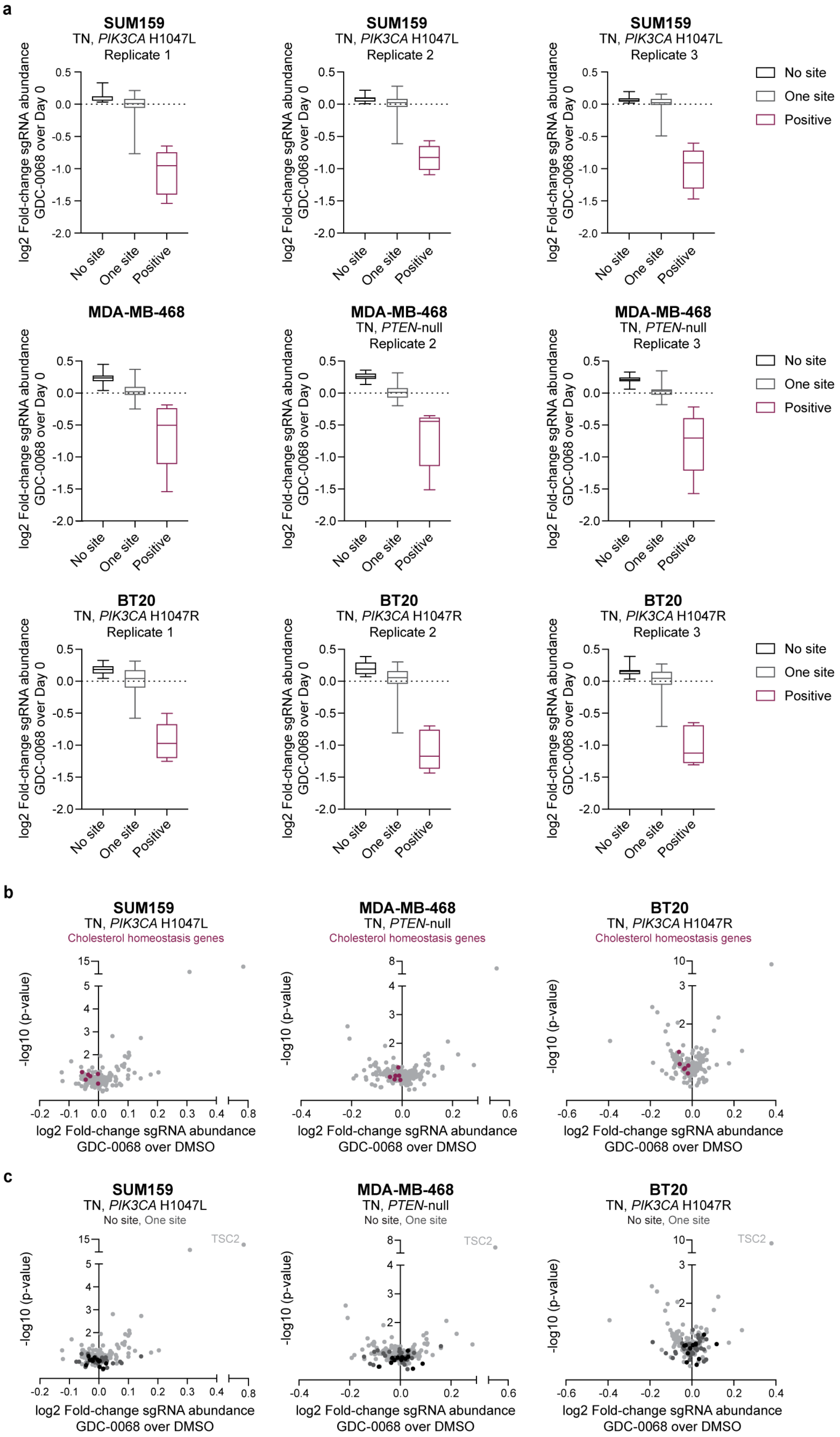


**Extended Data Fig.1 | Data supporting Fig.1 Custom CRISPR/Cas9 minipool library screen validates cholesterol homeostasis hits in a panel of TNBC cells. a,** Comparison of the log2 fold-change in sgRNA abundance for no site, one site and positive control sgRNAs in the GDC-0068 arm compared to the Day 0 arm of the custom CRISPR/Cas9 minipool library screen. No site sgRNAs do not cut anywhere in the genome. One site sgRNAs introduce single cuts in the genome at intergenic sites that are predicted to have little to no deleterious effects. Positive control sgRNAs target genes that are essential for cell viability. Data are plotted as box and whisker plots by replicate for SUM159, MDA-MB-468 and BT20 cells. The box extends from the $25^{th}$ to $75^{th}$ percentiles, and whiskers extend from minimum to maximum values. **b,** Volcano plots showing log2 fold-change in sgRNA abundance in the GDC-0068 arm (SUM159: 4.34 μM, MDA-MB-468: 8.49 μM, BT20: 0.9 μM) of the custom CRISPR/Cas9 minipool screen compared to the DMSO arm versus -log10 (p-value). Cholesterol homeostasis genes with negative log2 fold-changes in all three TNBC cell lines are highlighted (*ABCA12*, *ABCB4*, *CYP39A1*, *SREBF2*, *TMEM97*, *VPS4B*). Data are represented as the mean of 3 technical replicates for each cell line. **c,** Volcano plots showing log2 fold-change in sgRNA abundance in the GDC-0068 arm of the custom CRISPR/Cas9 minipool screen compared to the DMSO arm versus -log10 (p-value). Controls are highlighted, including no site and one intergenic site controls and the positive control *TSC2*. Data are represented as the mean of 3 technical replicates for each cell line.

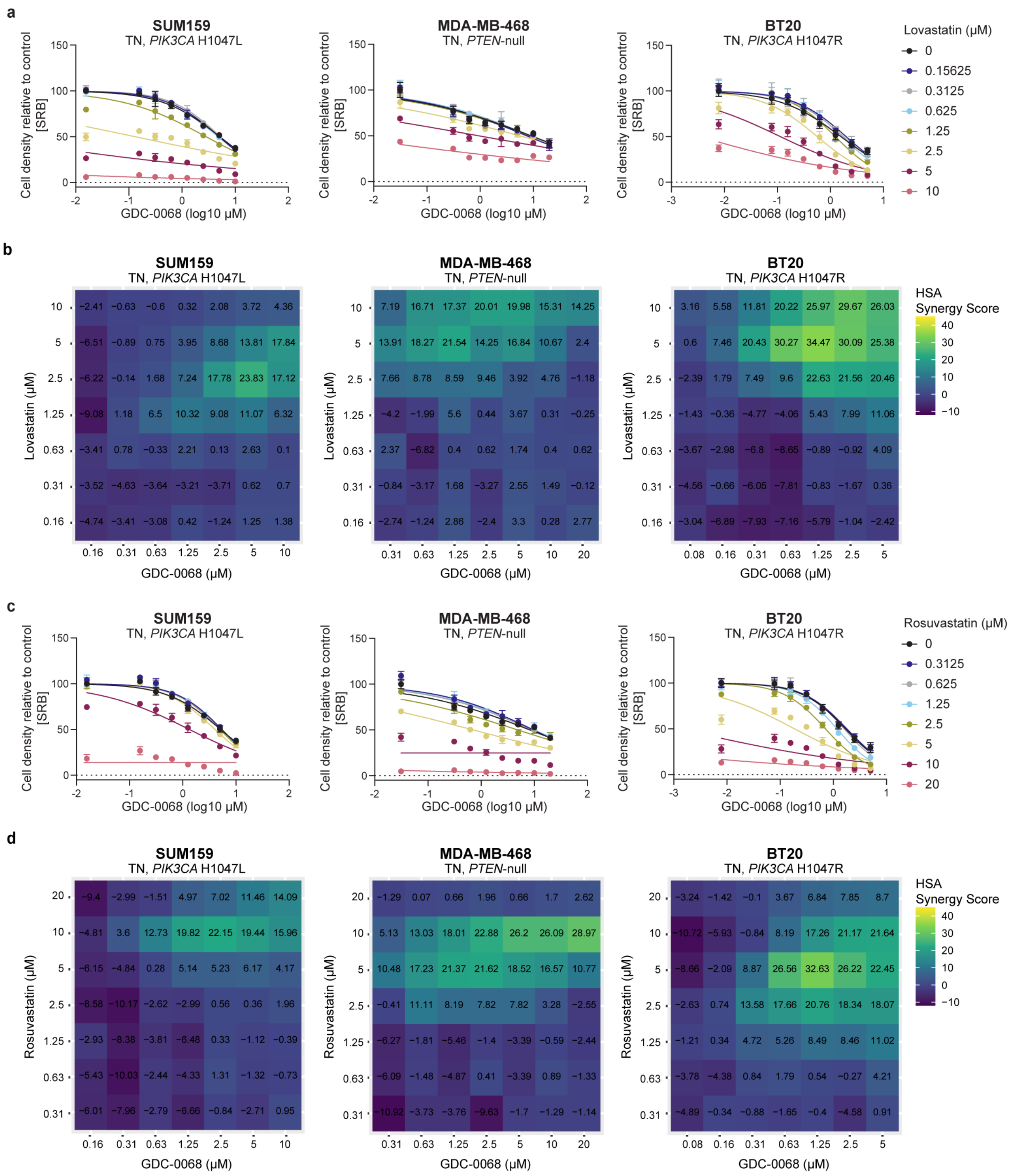


**Extended Data Fig.2 | Data supporting Fig.2 Lovastatin and rosuvastatin synergize with AKT inhibitors in TNBC cells. a,** TNBC cell lines (SUM159, MDA-MB-468, BT20) were treated with increasing doses of GDC-0068 (SUM159: 0-10 μM, MDA-MB-468: 0-20 μM, BT20: 0-5 μM) and lovastatin (0-10 μM) or **c,** rosuvastatin (0-20 μM) for 72 hours, and cell density was measured by SRB assay. Data are represented as mean ± SD (N=3 technical replicates). **b,** TNBC cell lines (SUM159, MDA-MB-468,

BT20) were treated with increasing doses of GDC-0068 (SUM159: 0-10 μM, MDA-MB-468: 0-20 μM, BT20: 0-5 μM) and lovastatin (0-10 μM) or **d,** rosuvastatin (0-20 μM) for 72 hours, and cell density was measured by SRB assay. HSA synergy scores were calculated using SynergyFinder and are reported in the heatmaps (N=3 technical replicates).

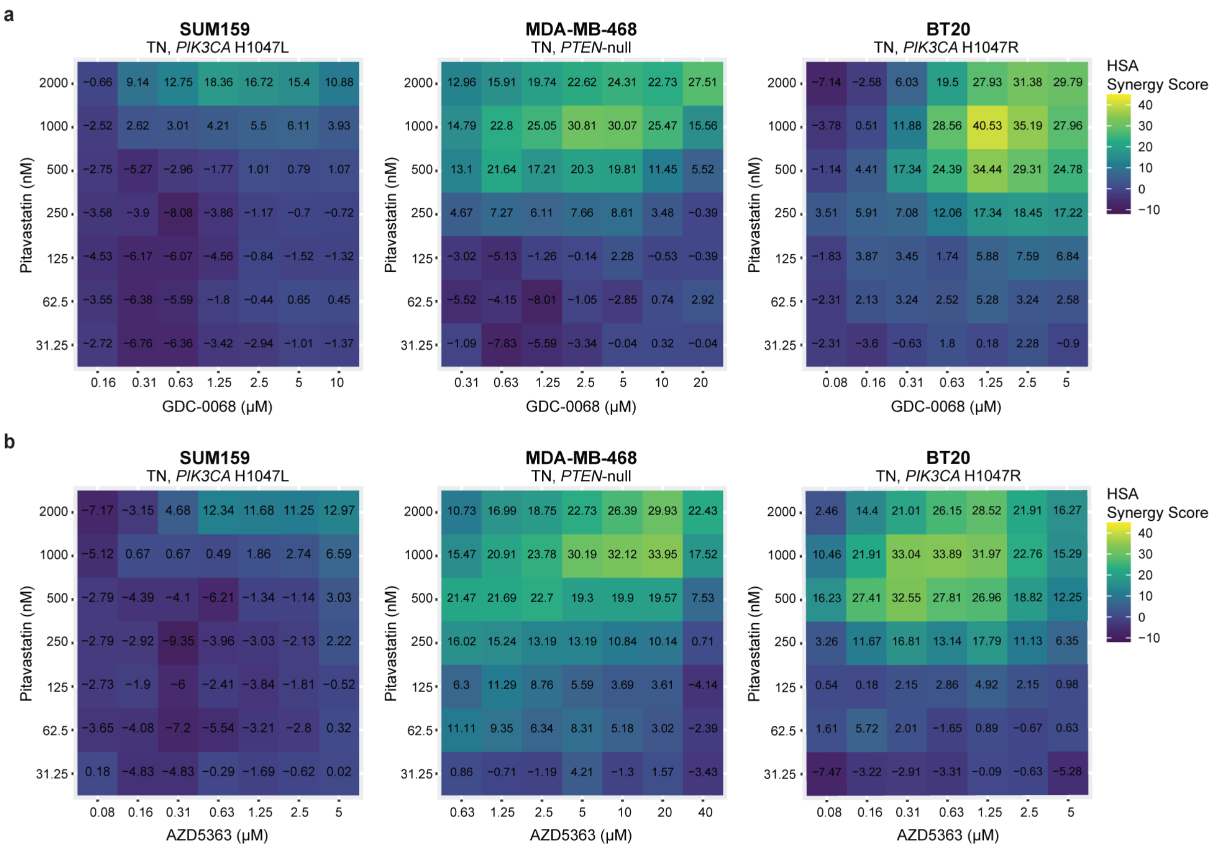


**Extended Data Fig.3 | Data supporting Fig.2 Pitavastatin synergizes with AKT inhibitors in TNBC cells. a-b,** TNBC cell lines (SUM159, MDA-MB-468, BT20) were treated with increasing doses of GDC-0068 (SUM159: 0-10 μM, MDA-MB-468: 0-20 μM, BT20: 0-5 μM) (**a**) or AZD5363 (SUM159: 0-5 μM, MDA-MB-468: 0-40 μM, BT20: 0-5 μM) (**b**) and pitavastatin (0-2000 nM) for 72 hours, and cell density was measured by SRB assay. HSA synergy scores were calculated using SynergyFinder and are reported in the heatmaps (N=3 technical replicates).

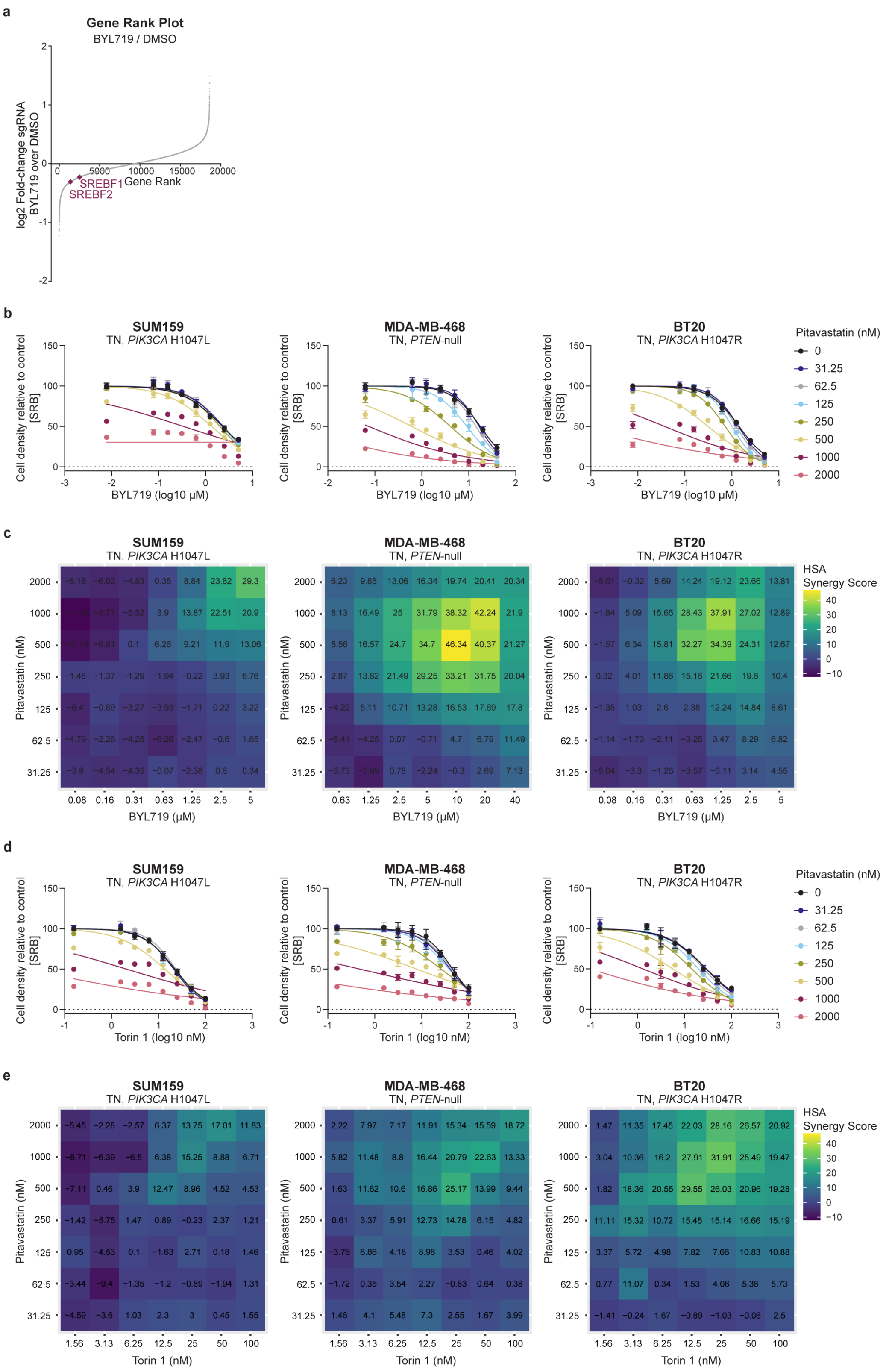


**Extended Data Fig.4 | Data supporting Fig.2 Pitavastatin synergizes with the PI3Kα inhibitor BYL719 and the mTORC1/2 inhibitor Torin 1 in TNBC cells. a,** Rank plots showing the log2 fold-change of each gene plotted against the rank dropout for the BYL719 treatment arm of the CRISPR/Cas9 screen compared to the DMSO arm. The transcription factors *SREBF1* and *SREBF2* are highlighted. The plot was generated using MAGeCK with a read count cutoff of 50 (N=3 technical replicates). **b,** TNBC cell lines (SUM159, MDA-MB-468, BT20) were treated with increasing doses of BYL719 (SUM159: 0-5 μM, MDA-MB-468: 0-40 μM, BT20: 0-5 μM) and pitavastatin (0-2000 nM) for 72 hours, and cell density was measured by SRB assay. Data are represented as mean ± SD (N=3 technical replicates). **c,** HSA synergy scores were calculated for the dose curves shown in **b** using SynergyFinder and are reported in the heatmaps (N=3 technical replicates). **d,** TNBC cell lines (SUM159, MDA-MB-468, BT20) were treated with increasing doses of Torin 1 (0-100 nM) and pitavastatin (0-2000 nM) for 72 hours, and cell density was measured by SRB assay. Data are represented as mean ± SD (N=2 technical replicates). **e,** HSA synergy scores were calculated for the dose curves shown in **d** using SynergyFinder and are reported in the heatmaps (N=2 technical replicates).

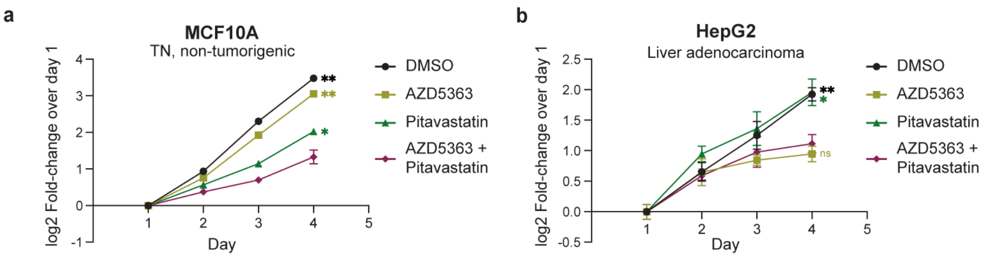


**Extended Data Fig.5 | Data supporting Fig.2 Pitavastatin and AZD5363 do not synergize in MCF10A and HepG2 cells. a,** The triple-negative, non-tumorigenic mammary epithelial cell line, MCF10A, was treated with DMSO, 3 μM AZD5363, 2 μM pitavastatin or a combination of AZD5363 and pitavastatin for 72 hours, and cell density was measured daily by SRB assay. Data are represented as mean ± SD (N=3 technical replicates). Statistical analysis was performed using two-way analysis of variance (ANOVA) with Dunnett's multiple comparison test; asterisks (*) indicate significant differences compared to the AZD5363 and pitavastatin combination treatment on day 4 (*, $p = 0.0332$, **, $p = 0.0021$). **b,** The liver adenocarcinoma cell line, HepG2, was treated with DMSO, 20 μM AZD5363, 0.5 μM pitavastatin or a combination of AZD5363 and pitavastatin for 72 hours, and cell density was measured daily by SRB assay. Data are represented as mean ± SD (N=3 technical replicates). Statistical analysis was performed using two-way analysis of variance (ANOVA) with Dunnett's multiple comparison test; asterisks (*) indicate significant differences compared to the AZD5363 and pitavastatin combination treatment on day 4 (*, $p = 0.0332$, **, $p = 0.0021$).

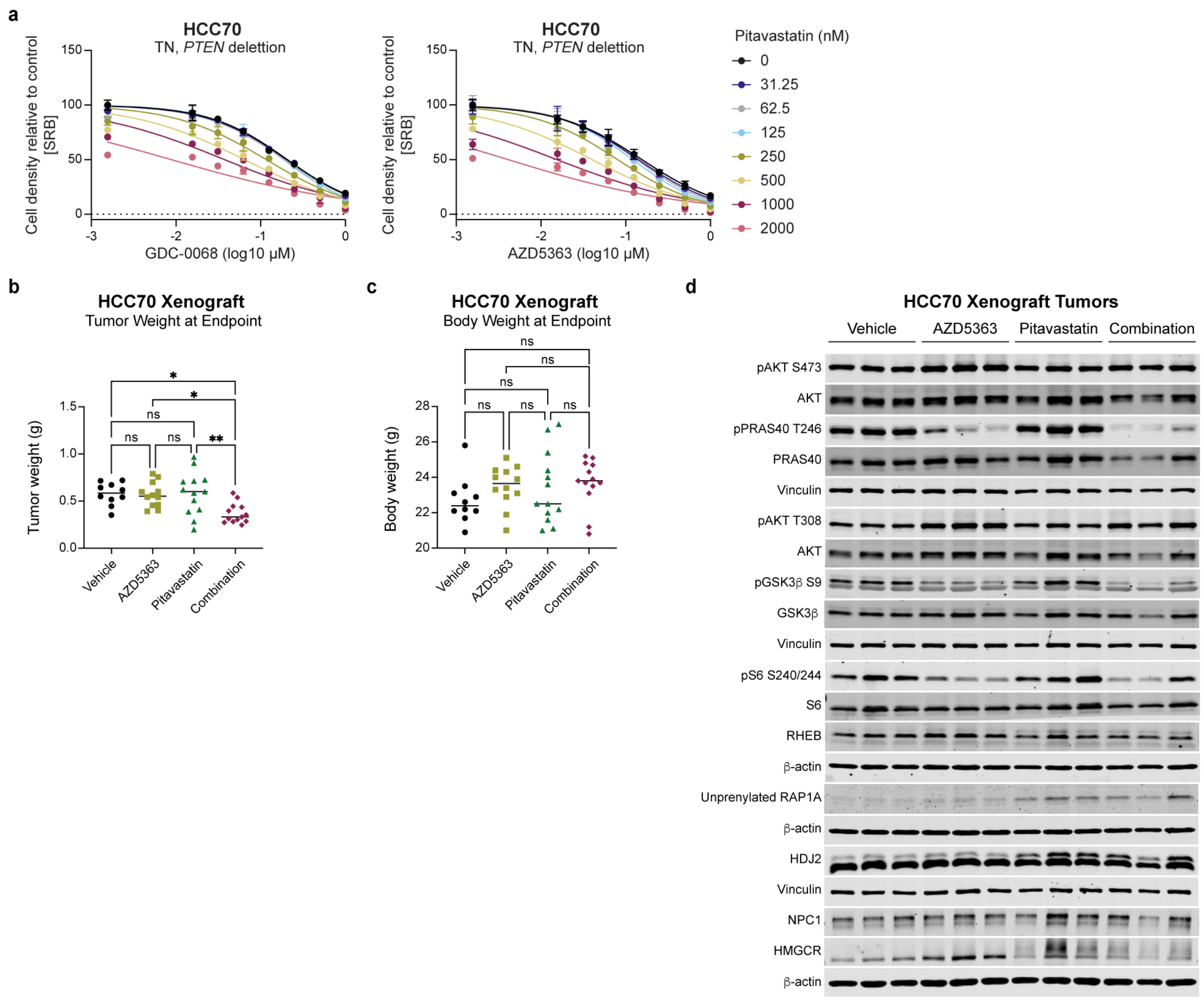


**Extended Data Fig.6 | Data supporting Fig.3 AZD5363 and pitavastatin synergize in HCC70 cells *in vitro* and in mouse xenografts with on-target efficacy. a,** HCC70 cells were treated with increasing doses of GDC-0068 (0-1 μM) or AZD5363 (0-1 μM) and pitavastatin (0-2000 nM) for 72 hours, and cell density was measured by SRB assay. Data are represented as mean ± SD (N=3 technical replicates). **b,** Tumor weight was measured at the endpoint. **c,** Mouse body weight was measured at the endpoint. For **b-c**, statistical analysis was performed using two-way analysis of variance (ANOVA) with Tukey's multiple comparison test; asterisks (*) indicate significant differences compared to the AZD5363 and pitavastatin combination treatment at the endpoint (*, $p = 0.0332$, **, $p = 0.0021$). **d,** Select tumors (3 per treatment group) were harvested 2 hours after the last AZD5363 treatment and 6 hours after the last pitavastatin treatment and immunoblotted for $pAKT^{Ser473}$, $pPRAS40^{Thr246}$, $pAKT^{Thr308}$, $pGSK3\beta^{Ser9}$, $pS6^{Ser240/244}$, RHEB, unprenylated RAP1A, HDJ2, NPC1, HMGCR, vinculin and β-actin.

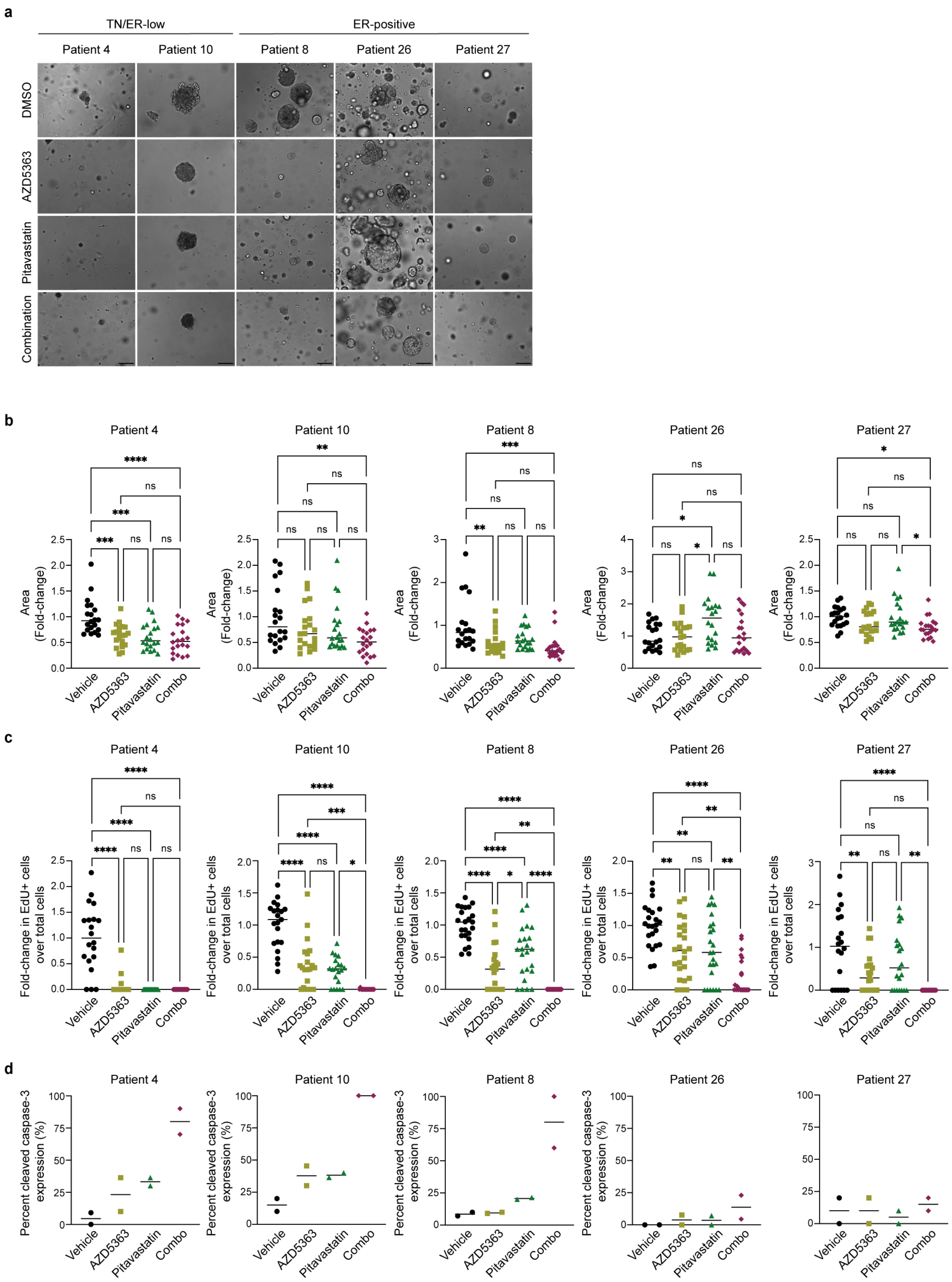


**Extended Data Fig.7 | Data supporting Fig.3 TN/ER-low PDOs are sensitive to combination AZD5363 and pitavastatin. a,** A panel of breast cancer PDOs were treated with DMSO, 1 μM AZD5363,

5 μM pitavastatin or the combination of AZD5363 and pitavastatin for 96 hours and organoid size and morphology were assessed. A representative image for each PDO in each treatment condition is shown. Scale bars are 200 μm. **b,** The area of 20 PDOs per treatment condition was quantified and normalized to the vehicle-treated condition. **c,** EdU+ cells were quantified for 20 PDOs per treatment condition and normalized to total cell number and to the vehicle-treated condition. For **b-c**, statistical analysis was performed using two-way analysis of variance (ANOVA) with Tukey's multiple comparison test; asterisks (*) indicate significant differences (*, $p = 0.0332$, **, $p = 0.0021$, ***, $p = 0.0002$, ****, $p < 0.0001$). **d,** The percentage of cleaved caspase-3-expressing PDOs was quantified for 2 images per treatment condition.

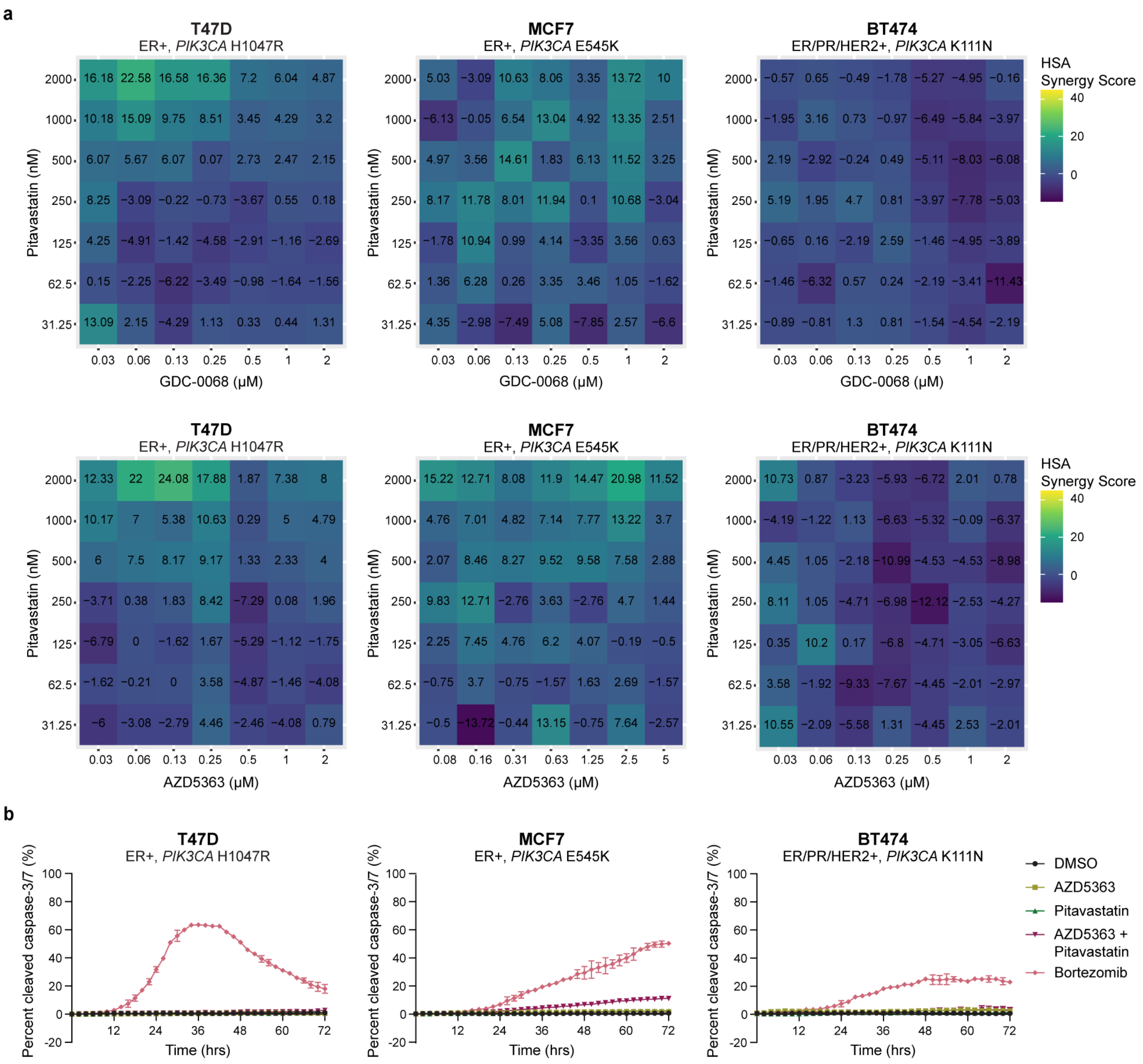


**Extended Data Fig.8 | Data supporting Fig.4 Pitavastatin does not synergize with AKT inhibition to induce cell death in ER-positive breast cancer cells. a,** ER-positive breast cancer cell lines (T47D, MCF7, BT474) were treated with increasing doses of GDC-0068 (0-2 μM) or AZD5363 (T47D: 0-2 μM, MCF7: 0-5 μM, BT474: 0-2 μM) and pitavastatin (0-2000 nM) for 72 hours, and cell density was measured by SRB assay. HSA synergy scores were calculated using SynergyFinder and are reported in the heatmaps (N=3 technical replicates). **b,** ER-positive breast cancer cell lines were treated with DMSO, AZD5363 (T47D: 0.25 μM, MCF7: 1.25 μM, BT474: 0.25 μM), pitavastatin (2 μM), a combination of AZD5363 and pitavastatin or bortezomib (10 μM) for 72 hours, and total cell number (rapid red nuclear dye) and number of dead cells (cleaved caspase-3/7 dye) were measured every 2 hours for 72 hours by

Incucyte live-cell analysis. Data are represented as mean ± SD of percent cleaved caspase-3/7 signal (N=4 technical replicates).

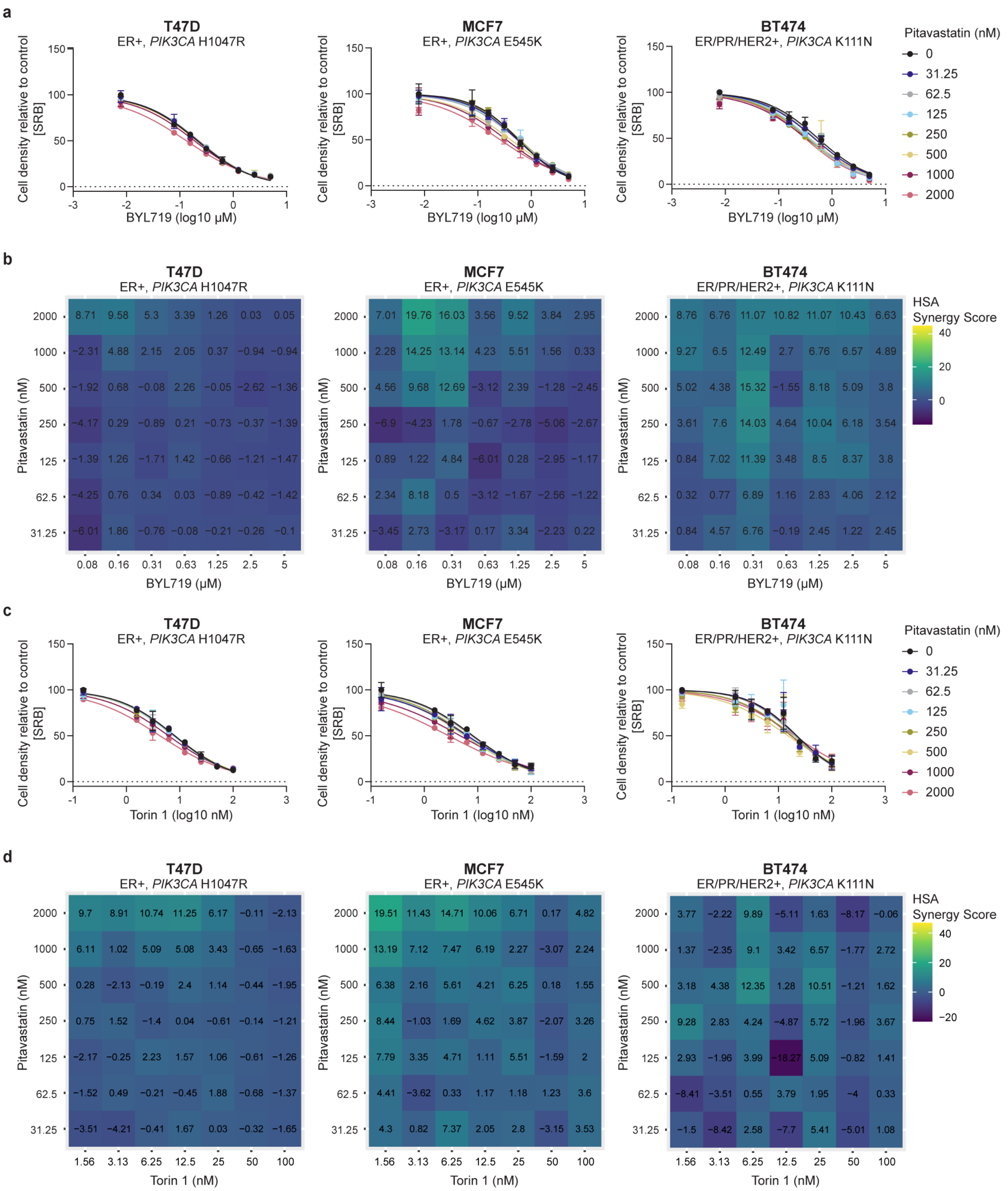


**Extended Data Fig.9 | Data supporting Fig.4 Pitavastatin does not synergize with the PI3Kα inhibitor BYL719 or the mTORC1/2 inhibitor Torin 1 in ER-positive breast cancer cells. a,** ER-positive breast cancer cell lines (T47D, MCF7, BT474) were treated with increasing doses of BYL719 (0-5 μM) and pitavastatin (0-2000 nM) for 72 hours, and cell density was measured by SRB assay. Data are represented as mean ± SD (N=3 technical replicates). **b,** HSA synergy scores were calculated for the dose curves shown in **a** using SynergyFinder and are reported in the heatmaps (N=3 technical replicates). **c,** ER-positive breast cancer cell lines (T47D, MCF7, BT474) were treated with increasing doses of Torin 1 (0-100 nM) and pitavastatin (0-2000 nM) for 72 hours, and cell density was measured by SRB assay. Data are represented as mean ± SD (N=2 technical replicates). **d,** HSA synergy scores were calculated for the dose curves shown in **c** using SynergyFinder and are reported in the heatmaps (N=2 technical replicates).

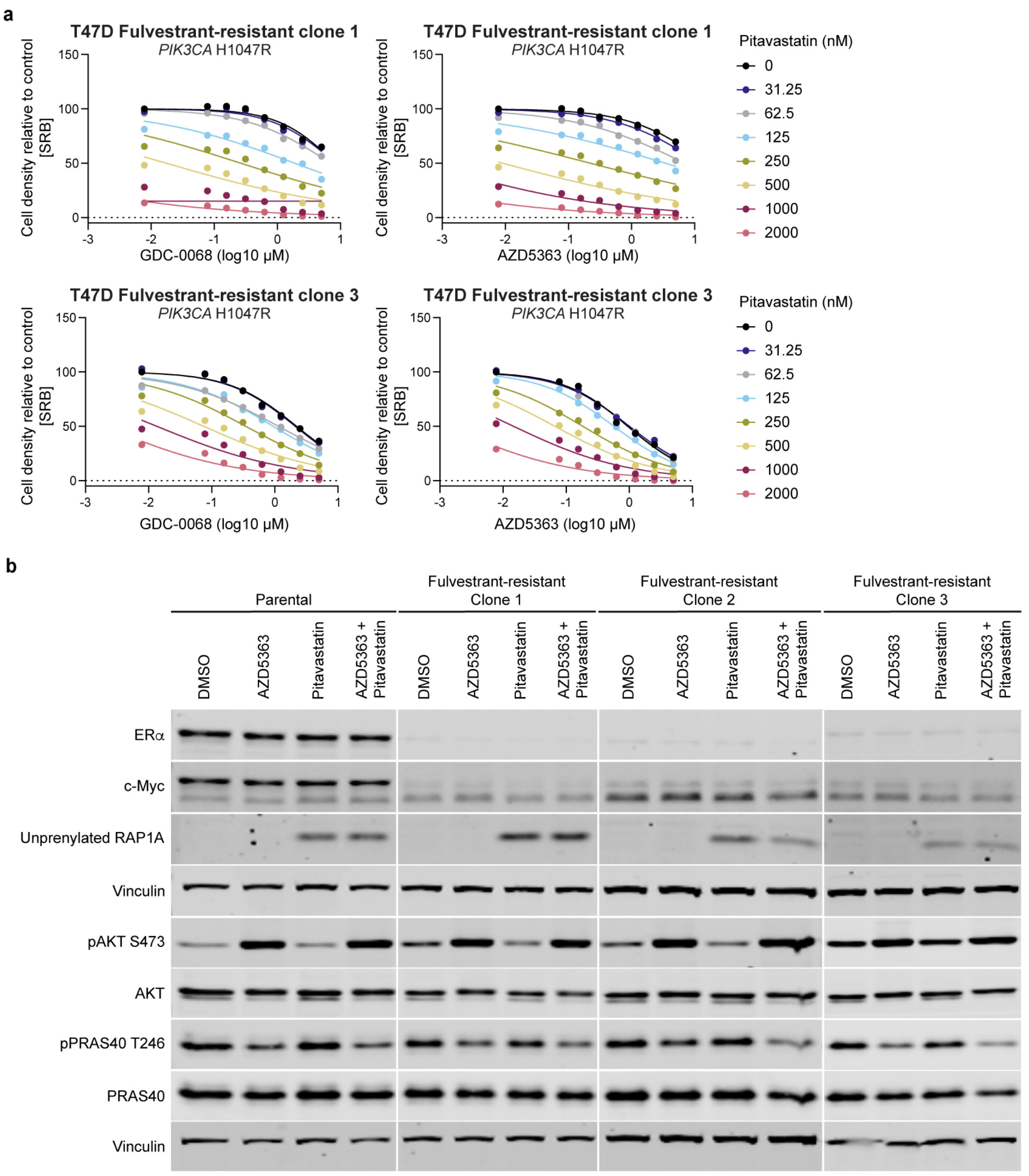


**Extended Data Fig.10 | Data supporting Fig.4 Pitavastatin synergizes with AKT inhibition in fulvestrant-resistant T47D cells. a,** Parental and fulvestrant-resistant T47D cells were treated with increasing doses of GDC-0068 (0-5 μM) or AZD5363 (0-5 μM) and pitavastatin (0-2000 nM) for 72 hours, and cell density was measured by SRB assay (N=1 technical replicate). **b,** Immunoblots of ERα, c-Myc, unprenylated RAP1A, vinculin, pAKT$^{Ser473}$ and pPRAS40$^{Thr246}$ in parental and fulvestrant-resistant T47D cells treated with DMSO, AZD5363 (0.25 μM), pitavastatin (0.5 μM) or the combination of AZD5363 and pitavastatin for 72 hours.

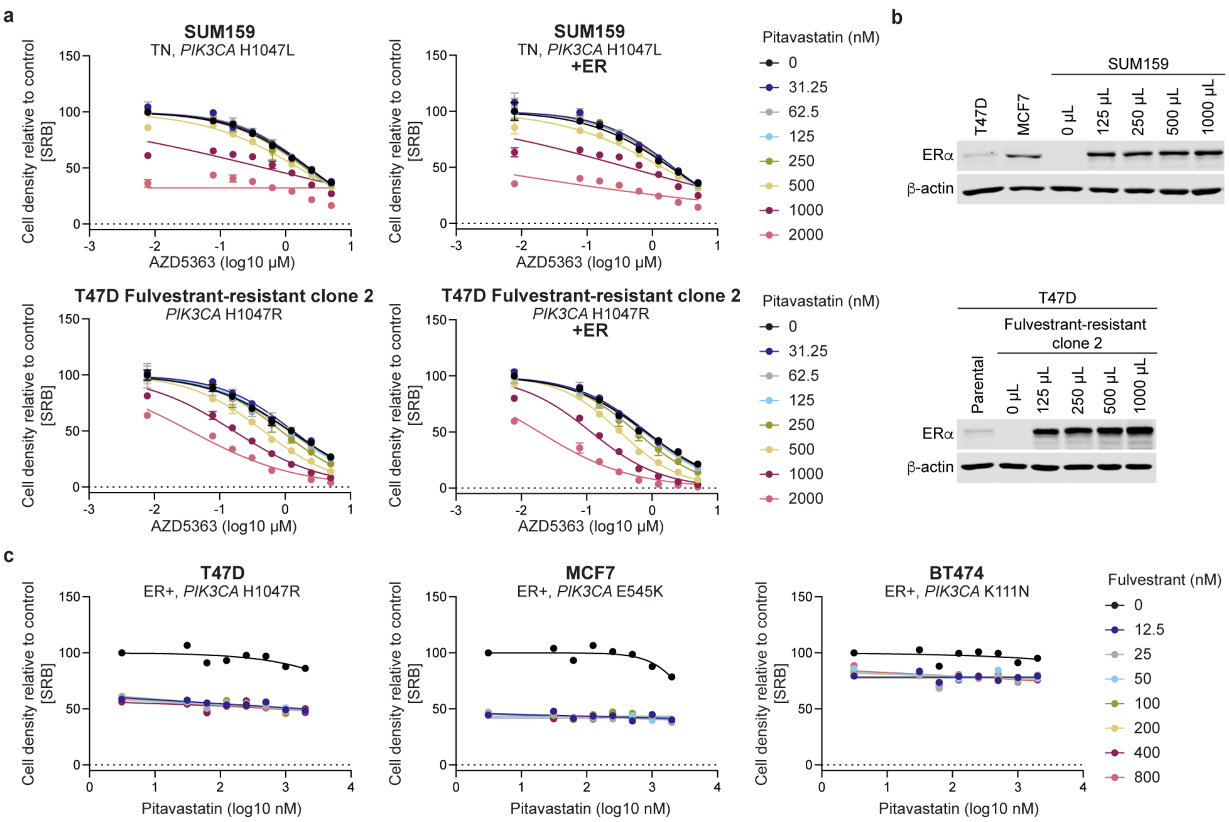


**Extended Data Fig.11 | Data supporting Fig.4 Modulation of ER expression does not affect pitavastatin sensitivity. a,** ER-negative breast cancer cell lines (SUM159, fulvestrant-resistant T47D clone 2) were transduced with lentiviral vectors to overexpress estrogen receptor (ER) and were maintained in culture for approximately 1 week. Parental and ER-overexpressing cells were then treated with increasing doses of AZD5363 (0-5 µM) and pitavastatin (0-2000 nM) for 72 hours, and cell density was measured by SRB assay. Data are represented as mean ± SD (N=2 technical replicates). **b,** Immunoblots of ERα and β-actin in SUM159 and fulvestrant-resistant T47D cells overexpressing ER with pHAGE-ESR1 compared to ER-positive breast cancer cell lines (T47D, MCF7) and parental T47D cells. **c,** ER-positive breast cancer cell lines (T47D, MCF7, BT474) were treated with increasing doses of fulvestrant (0-800 nM) and pitavastatin (0-2000 nM) for 72 hours, and cell density was measured by SRB assay. Data represent a single technical replicate (N=1 technical replicate).

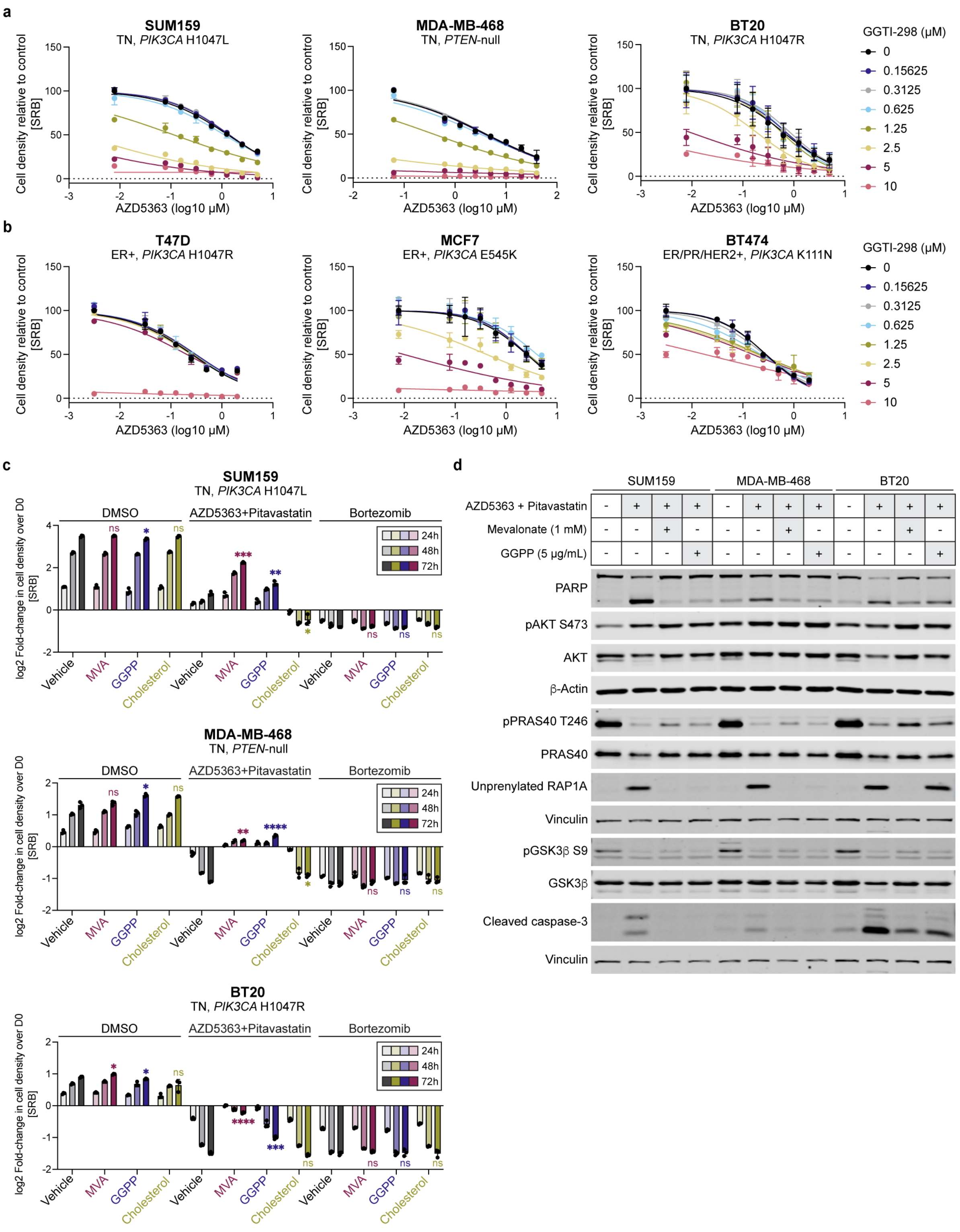


**Extended Data Fig.12 | Data supporting Fig.4 AKT inhibitors synergize with cholesterol biosynthesis inhibition to deplete GGPP. a,** TNBC cell lines (SUM159, MDA-MB-468, BT20) were treated with increasing doses of AZD5363 (SUM159: 0-5 µM, MDA-MB-468: 0-40 µM, BT20: 0-5 µM) and

GGTI-298 (0-10 μM) for 72 hours, and cell density was measured by SRB assay. Data are represented as mean ± SD (N=2 technical replicates). **b,** ER-positive breast cancer cell lines (T47D, MCF7, BT474) were treated with increasing doses of AZD5363 (T47D: 0-2 μM, MCF7: 0-5 μM, BT474: 0-2 μM) and GGTI-298 (0-10 μM) for 72 hours, and cell density was measured by SRB assay. Data are represented as mean ± SD (N=2 technical replicates). **c,** TNBC cell lines (SUM159, MDA-MB-468, BT20) were treated with DMSO, a combination of AZD5363 (SUM159: 5 μM, MDA-MB-468: 15 μM, BT20: 1.25 μM) and pitavastatin (SUM159: 4 μM, MDA-MB-468: 2 μM, BT20: 2 μM) or bortezomib (10 μM) and supplemented with vehicle (7:3 MeOH:$NH_4OH$), mevalonate (1 mM), GGPP (5 μg/mL) or cholesterol (5 μg/mL) for 24, 48 or 72 hours. Cell density was measured by SRB assay at each time point. Data are represented as mean ± SD (N=3 technical replicates). Statistical analysis was performed for the 72-hour time point using two-way analysis of variance (ANOVA) with Dunnett's multiple comparison test; asterisks (*) indicate significant differences compared to the 72-hour vehicle supplemented condition within the DMSO, AZD5363 and pitavastatin or bortezomib treatment group (*, p = 0.0332, **, p = 0.0021, ***, p = 0.0002, ****, p < 0.0001). **d,** Immunoblots of PARP, $pAKT^{Ser473}$, $pPRAS40^{Thr246}$, unprenylated RAP1A, $pGSK3\beta^{Ser9}$, cleaved caspase-3, β-actin and vinculin in TNBC cell lines (SUM159, MDA-MB-468, BT20) treated with DMSO or a combination of AZD5363 (SUM159: 5 μM, MDA-MB-468: 15 μM, BT20: 1.25 μM) and pitavastatin (SUM159: 4 μM, MDA-MB-468: 2 μM, BT20: 2 μM) and supplemented with vehicle (7:3 MeOH:$NH_4OH$), mevalonate (1 mM), GGPP (5 μg/mL) or cholesterol (5 μg/mL) for 22 hours.

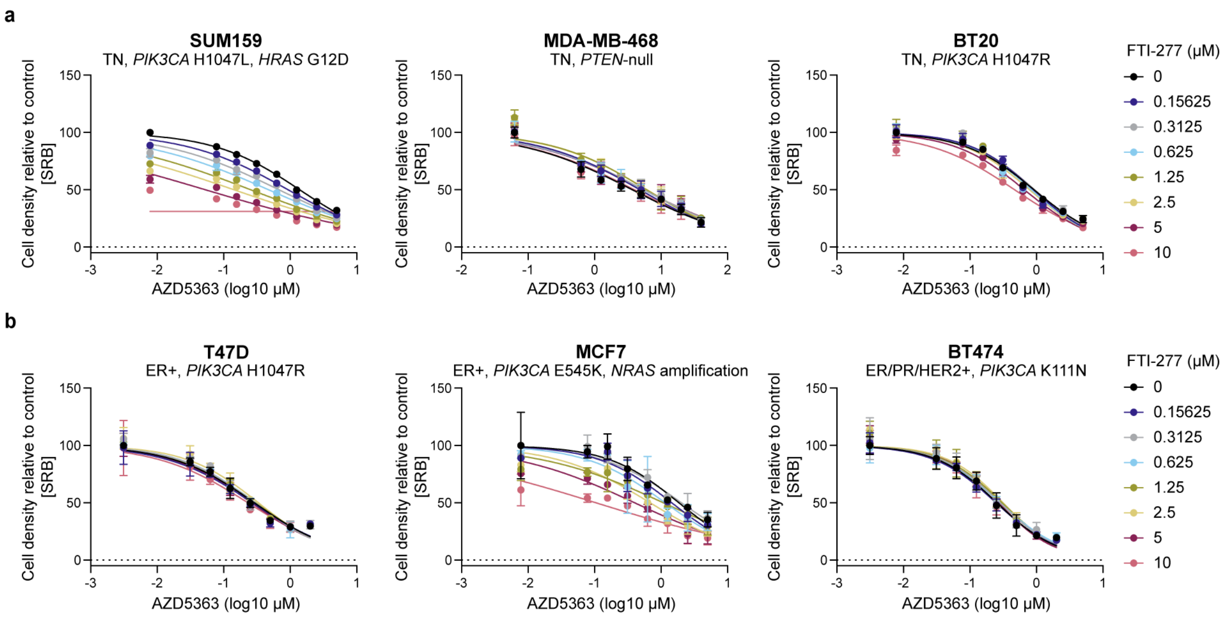


**Extended Data Fig.13 | Data supporting Fig.4 Farnesyltransferase inhibition with FTI-277 synergizes with AZD5363 in *RAS*-altered breast cancer cells. a,** TNBC cell lines (SUM159, MDA-MB-468, BT20) were treated with increasing doses of AZD5363 (SUM159: 0-5 µM, MDA-MB-468: 0-40 µM, BT20: 0-5 µM) and FTI-277 (0-10 µM) for 72 hours, and cell density was measured by SRB assay. Data are represented as mean ± SD (N=2 technical replicates). **b,** ER-positive breast cancer cell lines (T47D, MCF7, BT474) were treated with increasing doses of AZD5363 (T47D: 0-2 µM, MCF7: 0-5 µM, BT474: 0-2 µM) and FTI-277 (0-10 µM) for 72 hours, and cell density was measured by SRB assay. Data are represented as mean ± SD (N=2 technical replicates).

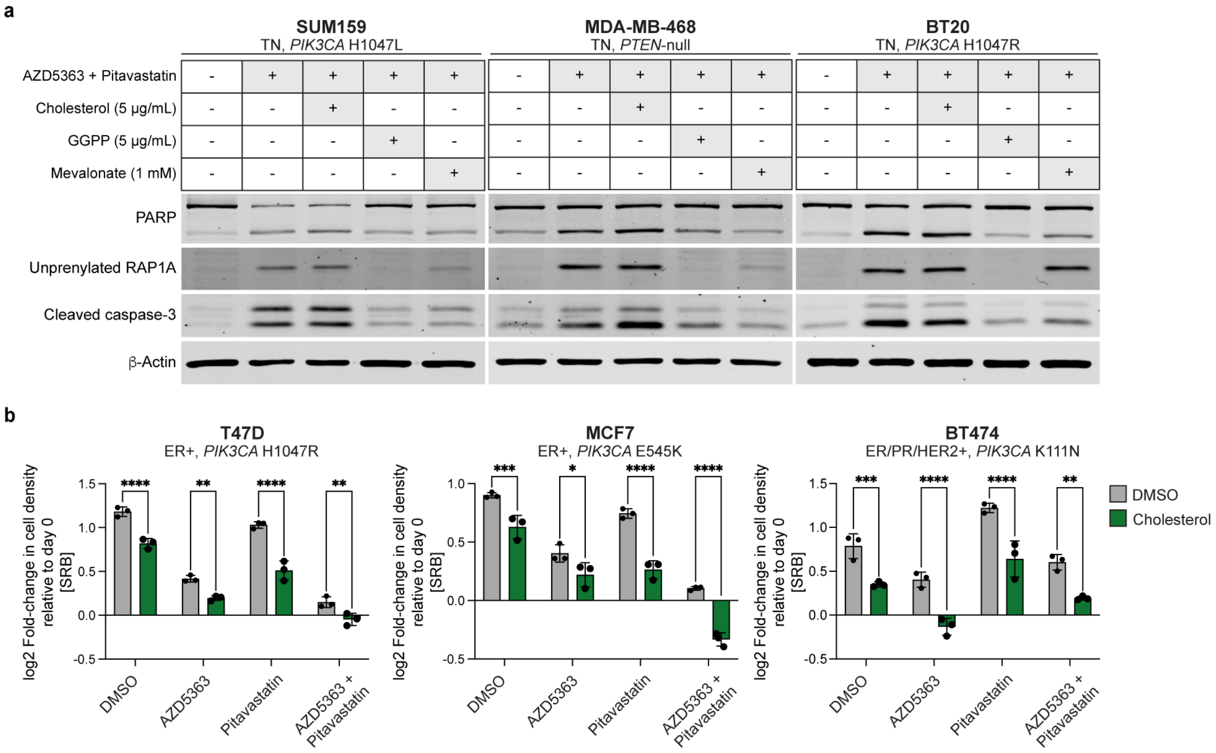


**Extended Data Fig.14 | Data supporting Fig.4 Cholesterol supplementation potentiates the cytotoxicity of AZD5363 and pitavastatin in TN and ER+ breast cancer cells. a,** Immunoblots of PARP, unprenylated RAP1A, cleaved caspase-3 and β-actin in TNBC cell lines (SUM159, MDA-MB-468, BT20) treated with DMSO or a combination of AZD5363 (SUM159: 5 µM, MDA-MB-468: 15 µM, BT20: 1.25 µM) and pitavastatin (SUM159: 4 µM, MDA-MB-468: 2 µM, BT20: 2 µM) and supplemented with vehicle (7:3 MeOH:$NH_4OH$), cholesterol (5 µg/mL), GGPP (5 µg/mL) or mevalonate (1 mM) for 22 hours. **b,** ER-positive breast cancer cell lines (T47D, MCF7, BT474) were treated with DMSO, AZD5363 (T47D: 0.25 µM, MCF7: 1.25 µM, BT474: 0.25 µM), pitavastatin (2 µM) or a combination of AZD5363 and pitavastatin for 72 hours and supplemented with DMSO or 5 µg/mL cholesterol. Cell density was measured by SRB assay. Data are represented as mean ± SD (N=3 technical replicates). Statistical analysis was performed using two-way analysis of variance (ANOVA) with Šidák's multiple comparison test; asterisks (*) indicate significant differences between the mean of DMSO-supplemented cells and the mean of cholesterol-supplemented cells for each treatment (*, p = 0.0332, **, p = 0.0021, ***, p = 0.0002, ****, p < 0.0001).

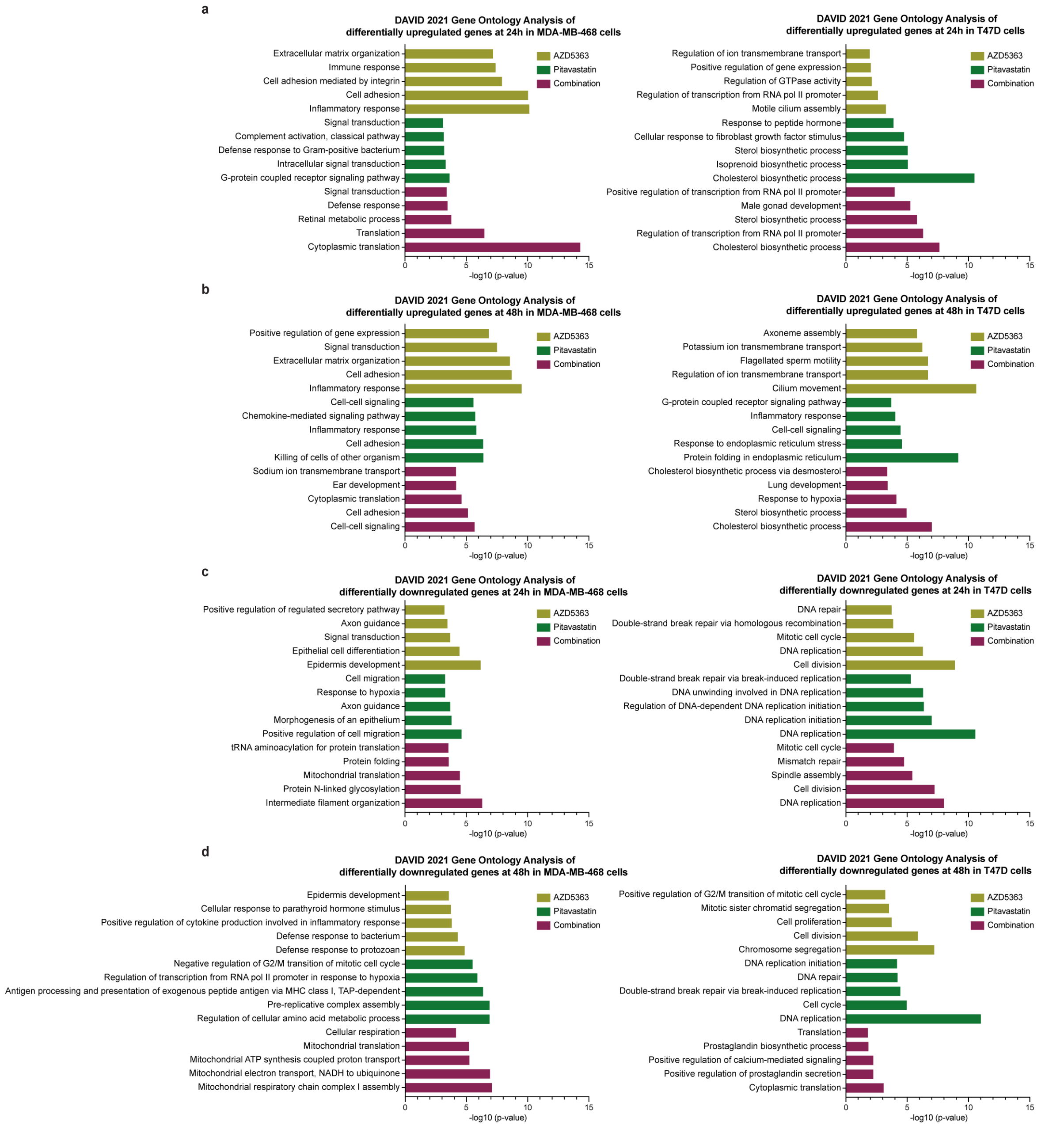


**Extended Data Fig.15 | Data supporting Fig.5 Cholesterol biosynthesis is uniquely upregulated after pitavastatin treatment in ER+ breast cancer cells. a-d,** DAVID 2021 gene ontology analysis of differentially regulated genes after AZD5363, pitavastatin or combination treatment for 24 or 48 hours in MDA-MB-468 and T47D cells from RNA-sequencing data. Pathways are plotted against the -log10 (p-value). **a,** Pathways enriched in uniquely upregulated genes in MDA-MB-468 and T47D cells at 24 hours. **b,** Pathways enriched in uniquely upregulated genes in MDA-MB-468 and T47D cells at 48 hours. **c,**

Pathways enriched in uniquely downregulated genes in MDA-MB-468 and T47D cells at 24 hours. **d,** Pathways enriched in uniquely downregulated genes in MDA-MB-468 and T47D cells at 48 hours.

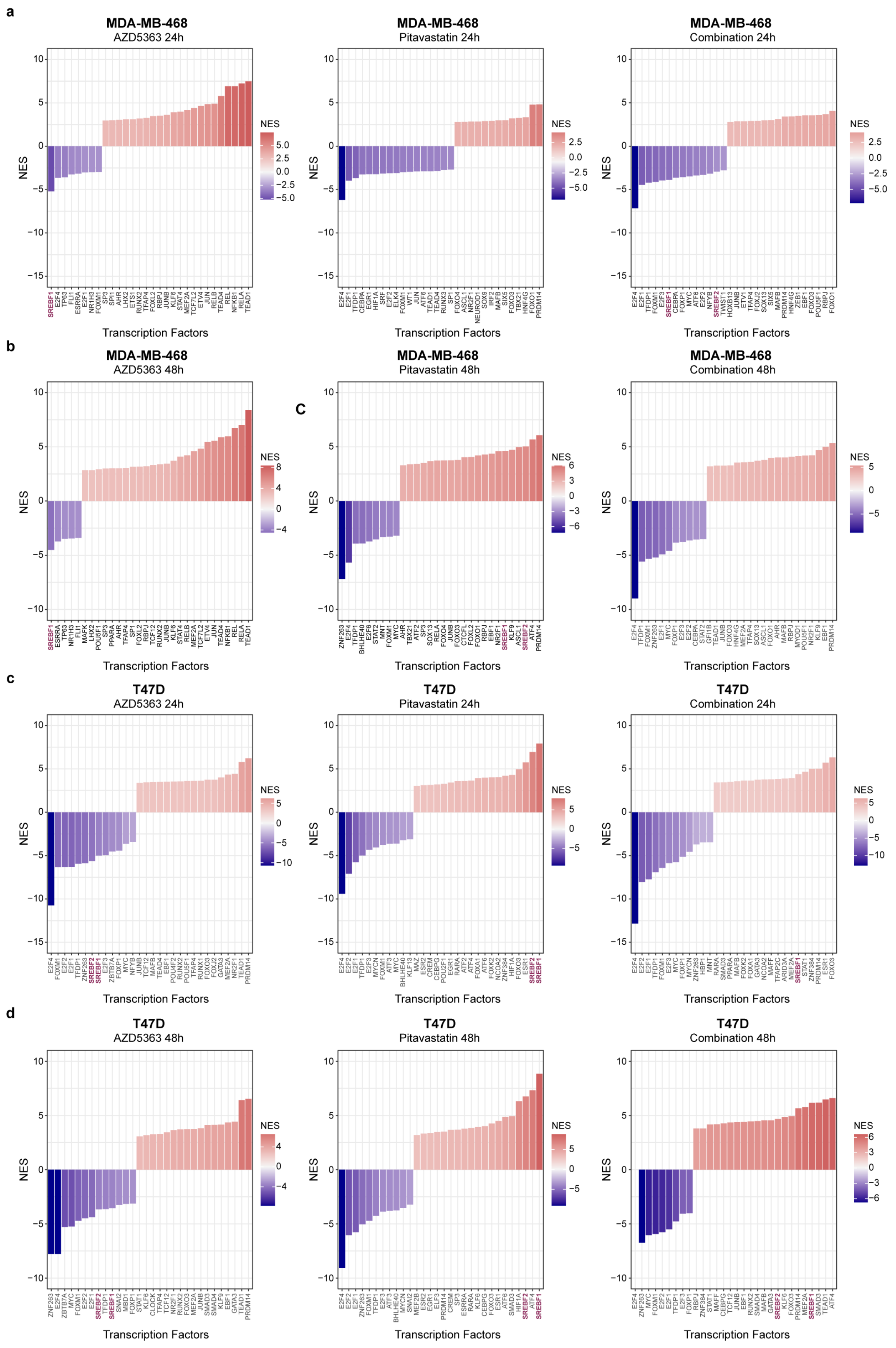


**Extended Data Fig.16 | Data supporting Fig.5 The predicted transcription factor activities of *SREBF1* and *SREBF2* are uniquely upregulated after AZD5363 and pitavastatin treatment in ER+**

**breast cancer cells. a-d,** DoRothEA transcription factor activity prediction of RNA sequencing data in MDA-MB-468 cells after 24 (**a**) or 48 hours (**b**) of treatment and in T47D cells after 24 (**c**) or 48 hours (**d**) of treatment. Normalized enrichment scores (NES) are plotted for transcription factors with activity that is predicted to be significantly altered by drug treatment. *SREBF1* and *SREBF2* are highlighted.

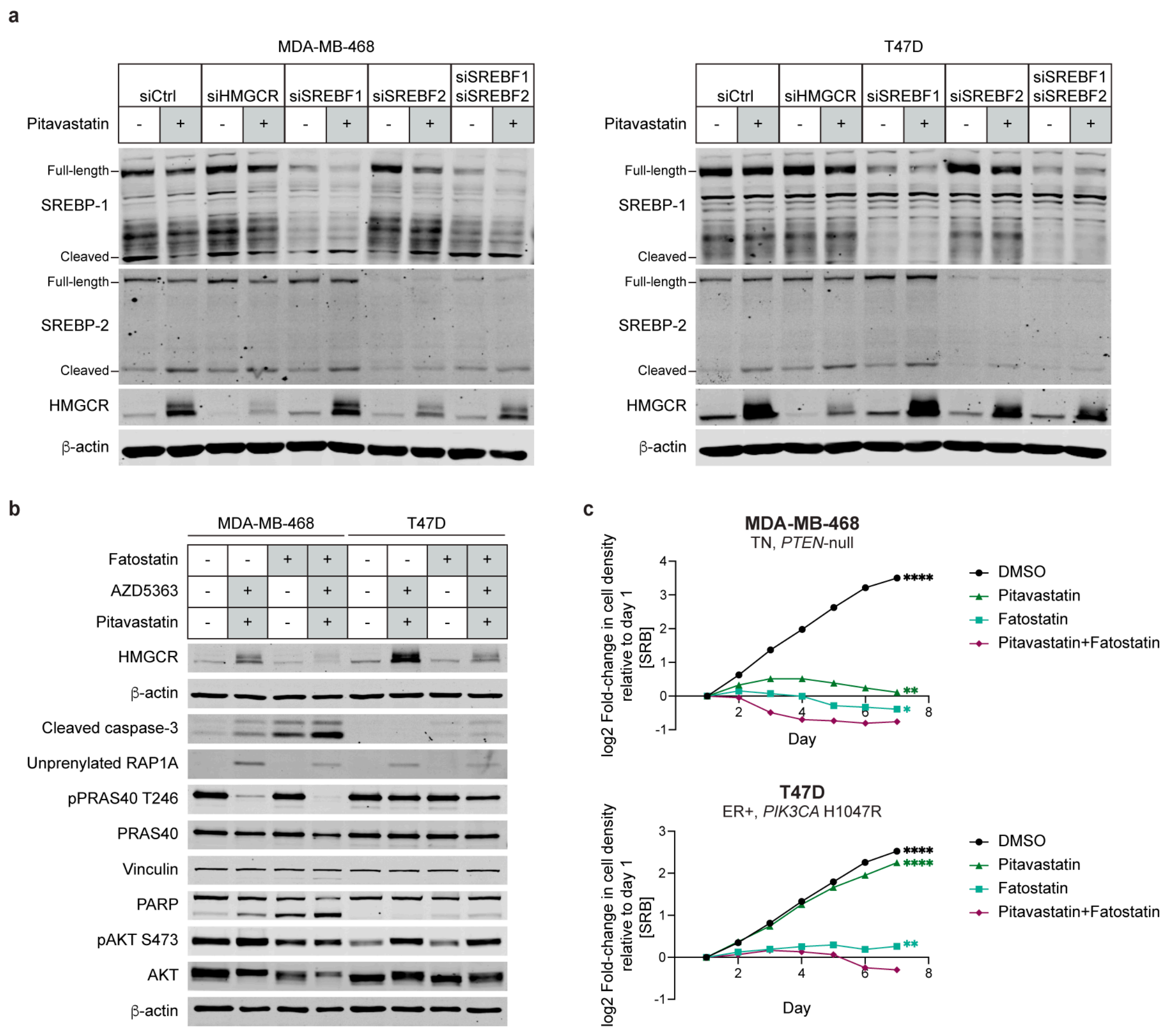


**Extended Data Fig.17 | Data supporting Fig.5 *SREBF2* depletion or inhibition sensitizes breast cancer cells to pitavastatin by limiting HMGCR upregulation. a,** Immunoblots of SREBP-1, SREBP-2, HMGCR and β-actin in MDA-MB-468 and T47D cells with siRNA knockdown of control (Ctrl), *HMGCR*, *SREBF1*, *SREBF2* or both *SREBF1* and *SREBF2* and treatment with DMSO or 2 μM pitavastatin for 24 hours. **b,** Immunoblots of HMGCR, cleaved caspase-3, unprenylated RAP1A, $pPRAS40^{Thr246}$, PARP, $pAKT^{Ser473}$ and β-actin in MDA-MB-468 and T47D cells treated with DMSO or 10 μM fatostatin for 24 hours, followed by treatment with AZD5363 (MDA-MB-468: 15 μM, T47D: 0.25 μM) and 2 μM pitavastatin for 24 hours. **c,** MDA-MB-468 and T47D cells were pre-treated with DMSO or 10 μM fatostatin for 2 hours, followed by treatment with DMSO or 2 μM pitavastatin for 7 days, and cell density was measured daily by SRB assay. Data are represented as mean ± SD (N=3 technical replicates). Statistical analysis was performed using two-way analysis of variance (ANOVA) with Dunnett's multiple comparison test;

asterisks (*) indicate significant differences compared to the pitavastatin and fatostatin combination treatment on day 7 (*, $p = 0.0332$, **, $p = 0.0021$, ***, $p = 0.0002$, ****, $p < 0.0001$).

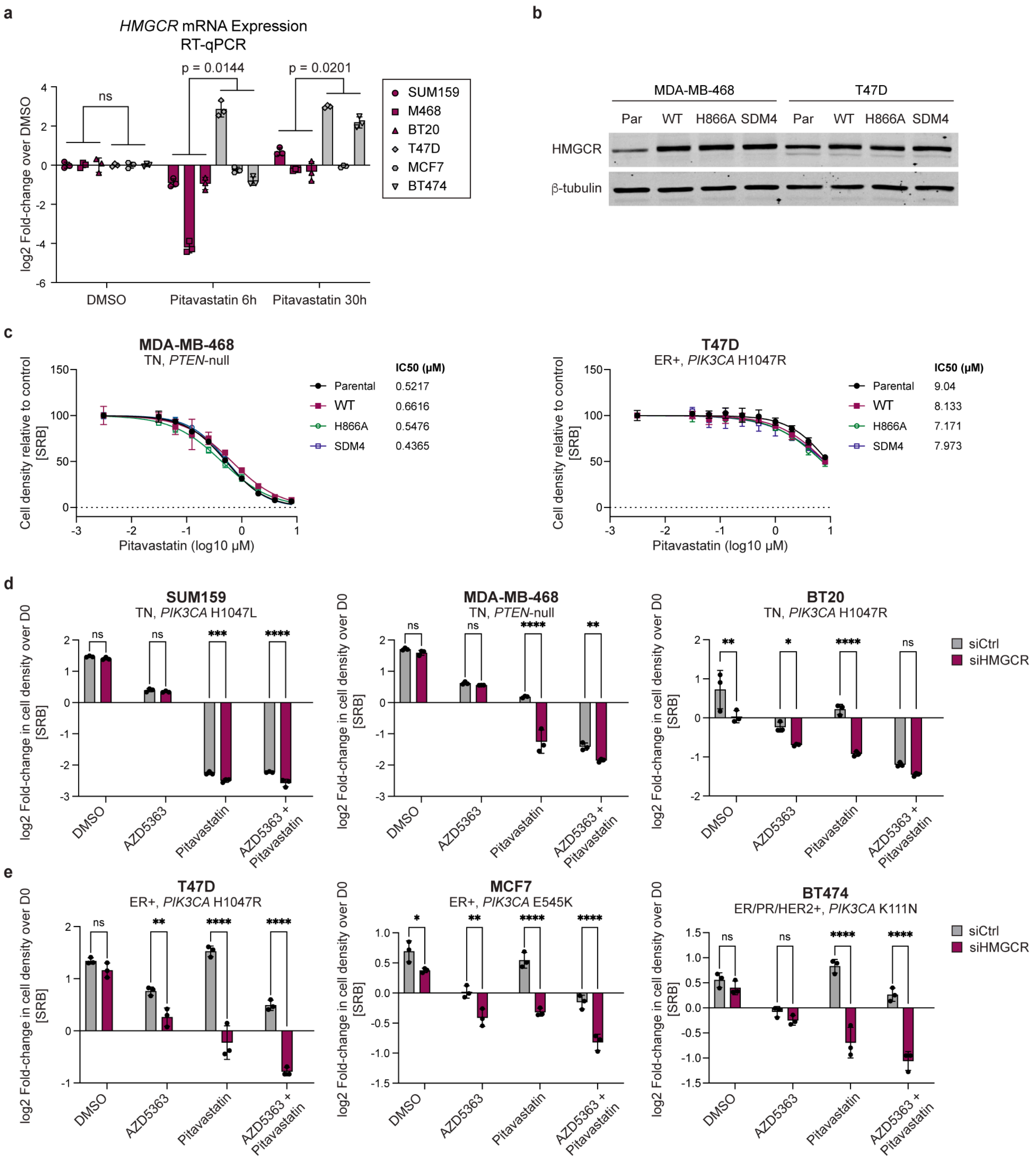


**Extended Data Fig.18 | Data supporting Fig.5 Depletion of HMGCR sensitizes ER+ breast cancer cells to pitavastatin and combination AZD5363 and pitavastatin. a,** TN (SUM159, MDA-MB-468, BT20) and ER-positive (T47D, MCF7, BT474) breast cancer cell lines were treated with DMSO or 2 µM pitavastatin for 6 or 30 hours, and *HMGCR* mRNA expression was measured by RT-qPCR. Data are represented as mean ± SD (N=3 technical replicates). Statistical analysis was performed using two-way analysis of variance (ANOVA) with Šidák's multiple comparison test. TNBC cell lines were compared to ER-positive breast cancer cell lines in each condition. **b,** Immunoblots of HMGCR and β-tubulin in parental and HMGCR-expressing MDA-MB-468 and T47D cells. Cells are expressing wild-type (WT), H866A or E559A, K691A, D767A and H866A pLenti6/V5-HMGCR. **c,** MDA-MB-468 and T47D cells expressing the pLenti6/V5-HMGCR constructs described in **b** were treated with a range of concentrations of pitavastatin (0-8 μM) for 72 hours, and cell density was measured by SRB assay. Data are represented as mean ± SD (N=3 technical replicates). **d,** TNBC cell lines (SUM159, MDA-MB-468, BT20) were transfected with siControl (siCtrl) or siHMGCR for 24 hours and then treated with DMSO, AZD5363 (SUM159: 5 μM, MDA-MB-468: 15 μM, BT20: 1.25 μM), pitavastatin (SUM159: 4 μM, MDA-MB-468: 2 μM, BT20: 2 μM) or a combination of AZD5363 and pitavastatin for 72 hours, and cell density was measured by SRB assay. Data are represented as mean ± SD (N=3 technical replicates). **e,** ER-positive breast cancer cell lines (T47D, MCF7, BT474) were transfected with siControl (siCtrl) or siHMGCR for 24 hours and then treated with DMSO, AZD5363 (T47D: 0.25 μM, MCF7: 1.25 μM, BT474: 0.25 μM), pitavastatin (2 μM) or a combination of AZD5363 and pitavastatin for 72 hours, and cell density was measured by SRB assay. Data are represented as mean ± SD (N=3 technical replicates). For **d-e,** statistical analysis was performed using two-way analysis of variance (ANOVA) with Šidák's multiple comparison test to compare the mean of siCtrl cells to the mean of siHMGCR cells for each treatment (*, $p = 0.0332$, **, $p = 0.0021$, ***, $p = 0.0002$, ****, $p < 0.0001$).

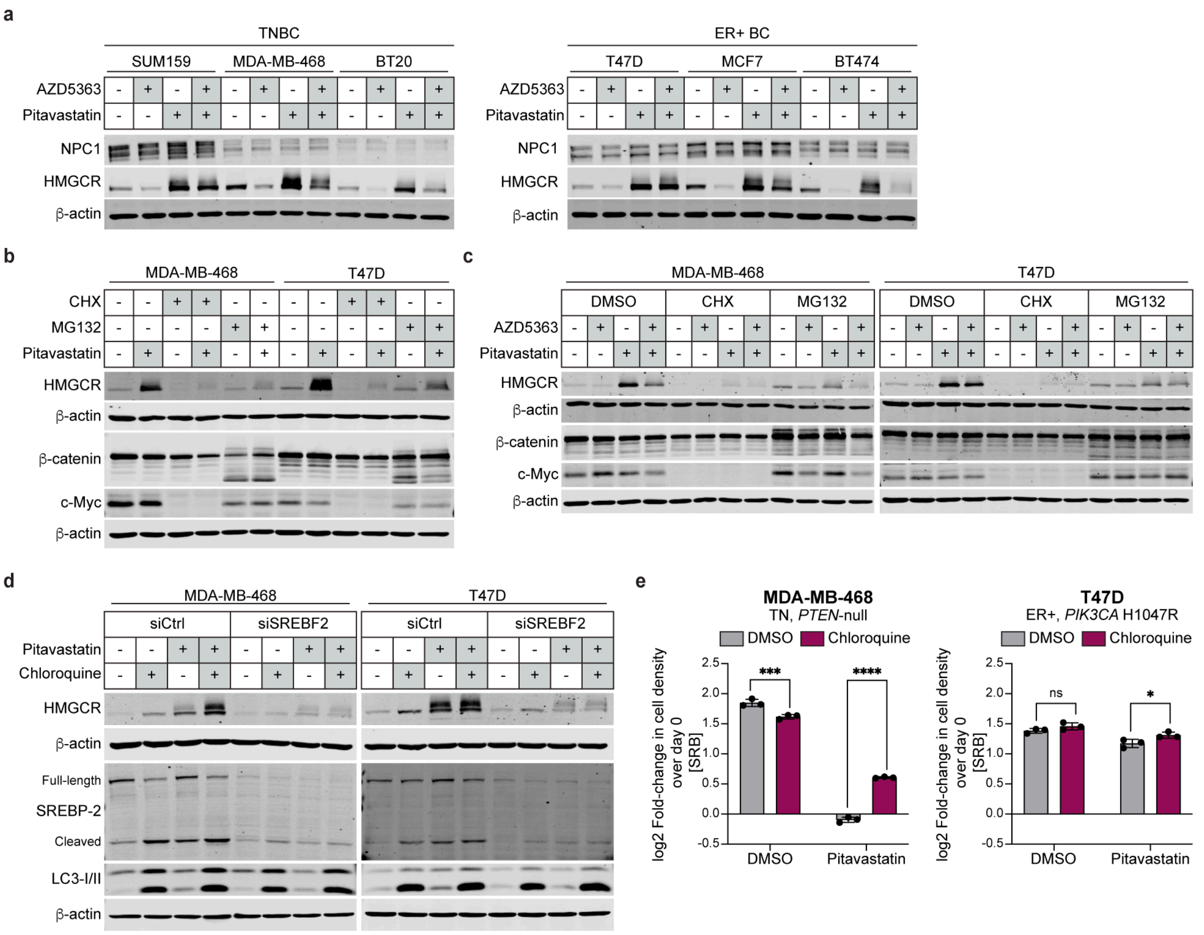


**Extended Data Fig.19 | Data supporting Fig.5 Pitavastatin-induced HMGCR upregulation is mediated by SREBP-2-dependent new synthesis of HMGCR. a,** Immunoblots of NPC1, HMGCR and β-actin in a panel of TN (SUM159, MDA-MB-468, BT20) and ER-positive (T47D, MCF7, BT474) breast cancer cells treated with DMSO or a combination of AZD5363 (SUM159: 2.5 μM, MDA-MB-468: 10 μM, BT20: 1.25 μM, T47D: 0.25 μM, MCF7: 1.25 μM, BT474: 0.25 μM) and 2 μM pitavastatin for 24h. **b,** Immunoblots of HMGCR, β-catenin, c-Myc and β-actin in MDA-MB-468 and T47D cells pre-treated with DMSO, 10 μg/mL cycloheximide (CHX) or 10 μM MG132 for 2 hours, followed by treatment with DMSO or 2 μM pitavastatin for 24 hours. **c,** Immunoblots of HMGCR, β-catenin, c-Myc and β-actin in MDA-MB-468 and T47D cells pre-treated with DMSO, 10 μg/mL cycloheximide (CHX) or 10 μM MG132 for 2 hours, followed by treatment with DMSO, AZD5363 (MDA-MB-468: 10 μM, T47D: 0.25 μM), 1 μM pitavastatin or the combination of AZD5363 and pitavastatin for 24 hours. **d,** Immunoblots of HMGCR, SREBP-2, LC3-I/II and β-actin in MDA-MB-468 and T47D cells transfected with siControl (siCtrl) or siSREBF2 for 24 hours followed by treatment with DMSO, 50 µM chloroquine, 2 µM pitavastatin or chloroquine and pitavastatin for 24 hours. **e,** MDA-MB-468 and T47D cells were co-treated with DMSO or 2 µM pitavastatin and DMSO or 5 µM chloroquine for 72 hours, and cell density was measured by SRB assay. Data are represented as mean ± SD (N=3 technical replicates). Statistical analysis was performed using two-way analysis of variance (ANOVA) with Šidák's multiple comparison test to compare the mean of DMSO-treated cells to the mean of chloroquine-treated cells for each treatment (*, $p = 0.0332$, ***, $p = 0.0002$, ****, $p < 0.0001$).

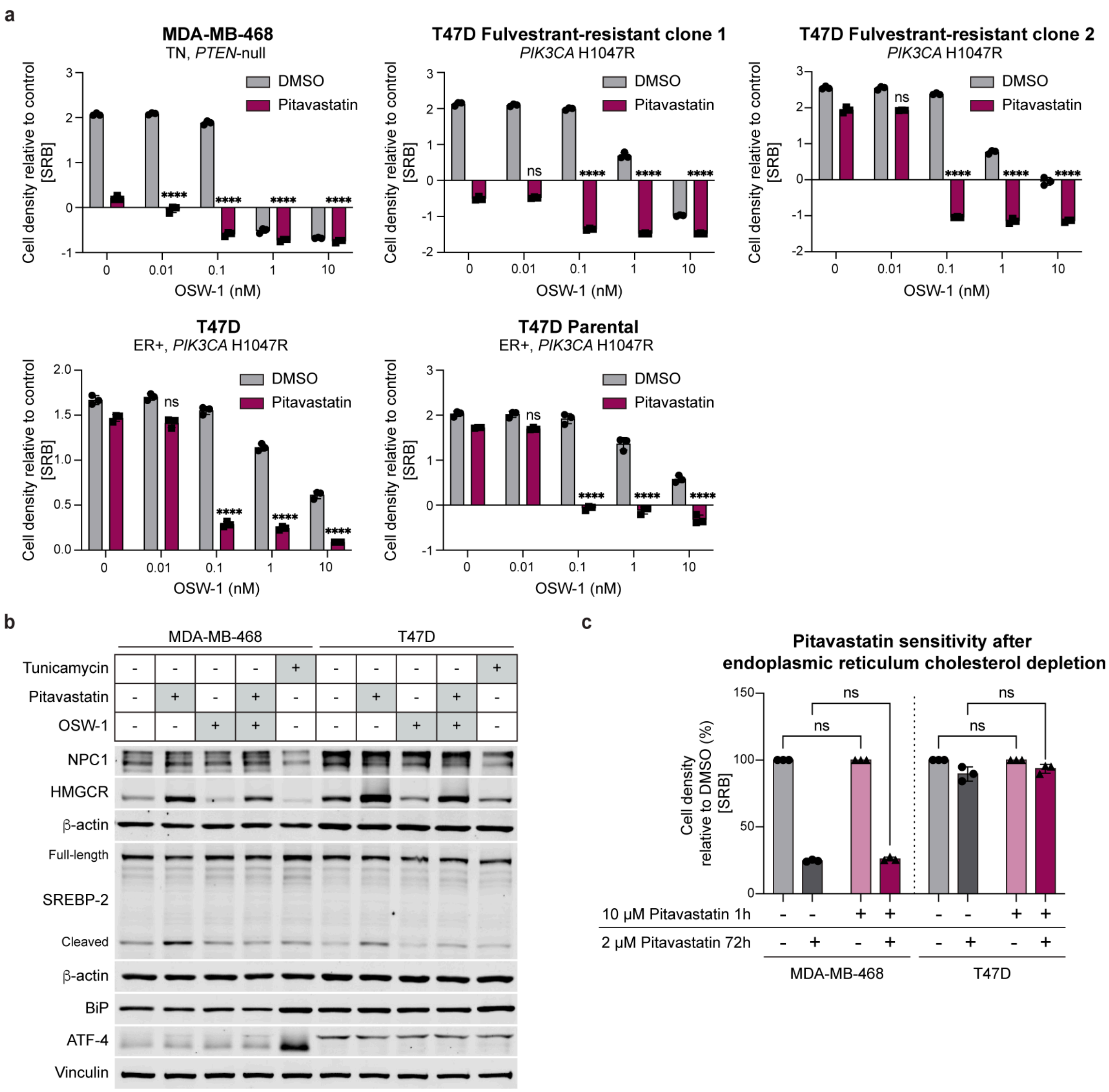


**Extended Data Fig.20 | Data supporting Fig.6 Accumulation of endoplasmic reticulum cholesterol sensitizes breast cancer cells to pitavastatin. a,** A panel of TN (MDA-MB-468, T47D fulvestrant-resistant clones 1 and 2) and ER-positive (T47D, parental T47D) breast cancer cells were co-treated with DMSO or 2 μM pitavastatin and increasing concentrations of OSW-1 (0-10 nM) for 72 hours, and cell density was measured by SRB assay. Data are represented as mean ± SD (N=3 technical replicates). Statistical analysis was performed using two-way analysis of variance (ANOVA) with Dunnett's multiple comparison test; asterisks (*) indicate significant differences compared to pitavastatin treatment without OSW-1 (****, $p < 0.0001$). **b,** Immunoblots of NPC1, HMGCR, SREBP-2, BiP, ATF-4, β-actin and vinculin in MDA-MB-468 and T47D cells treated with 2 μM tunicamycin, 2 μM pitavastatin, 0.1 nM OSW-1 or the combination of pitavastatin and OSW-1 for 24 hours. **c,** MDA-MB-468 and T47D cells were seeded into RPMI supplemented with 10% lipid-depleted serum and treated with 10 μM pitavastatin for 1 hour. The media was then changed to fresh RPMI supplemented with 10% fetal bovine serum (complete serum), and cells were treated with DMSO or 2 μM pitavastatin for 72 hours. Cell density was measured by SRB assay. Data are represented as mean ± SD (N=3 technical replicates). Statistical analysis was performed using two-way analysis of variance (ANOVA) with Tukey's multiple comparison test.

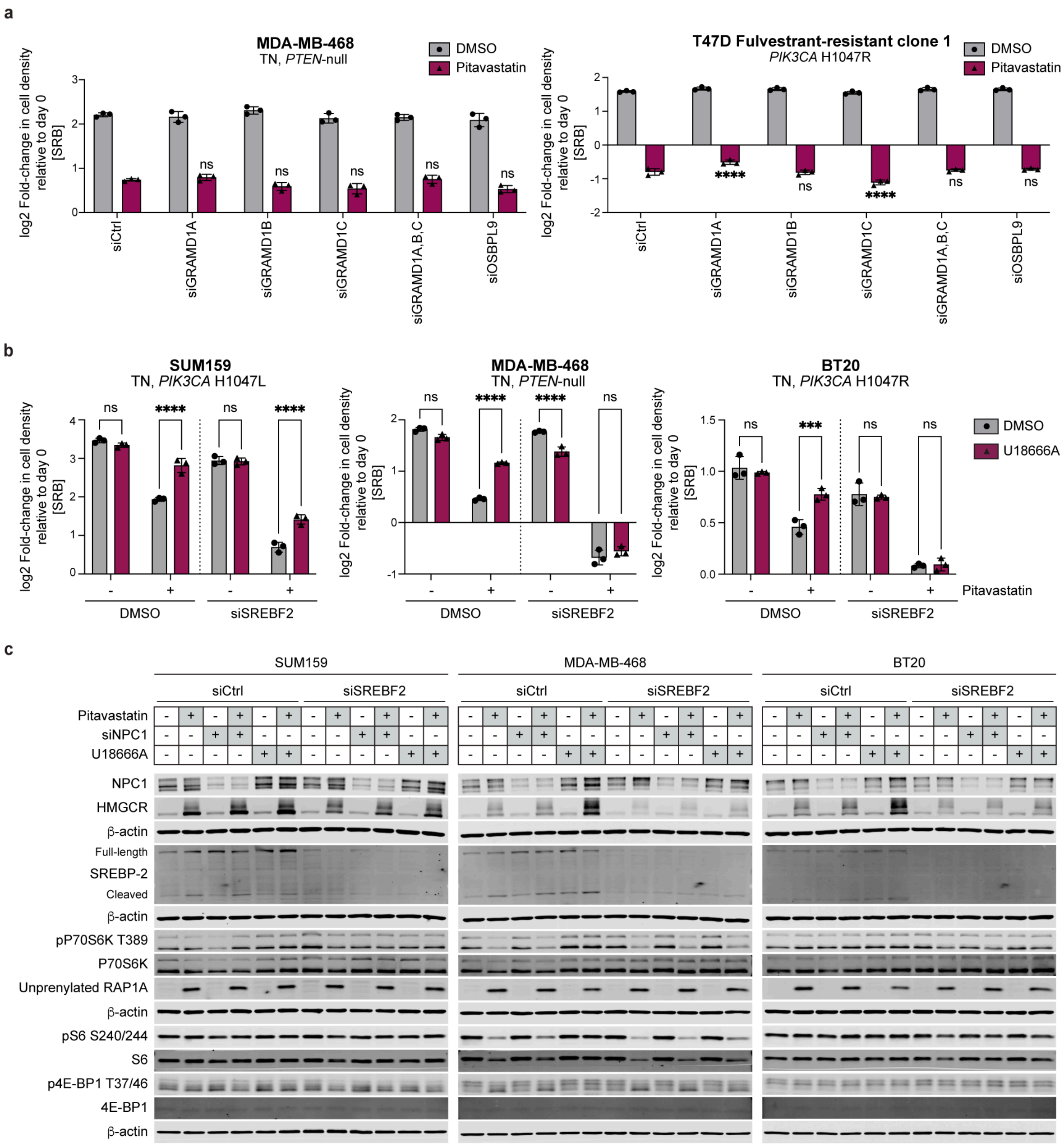


**Extended Data Fig.21 | Data supporting Fig.6 Depletion or inhibition of the cholesterol trafficking protein NPC1 rescues SREBP-2 activation after pitavastatin treatment in TNBC. a,** TNBC cell lines (MDA-MB-468, fulvestrant-resistant T47D clone 1) were transfected with siControl (siCtrl), siGRAMD1A, siGRAMD1B, siGRAMD1C, all 3 siGRAMD1s or siOSBPL9 for 24 hours and then treated with DMSO or 2 μM pitavastatin for 72 hours, and cell density was measured by SRB assay. Data are represented as mean ± SD (N=3 technical replicates). Statistical analysis was performed using two-way analysis of variance (ANOVA) with Dunnett's multiple comparison test; asterisks (*) indicate significant differences compared to the pitavastatin-treated siControl (siCtrl) condition for each cell line (****, $p < 0.0001$). **b,** TNBC cells (SUM159, MDA-MB-468, BT20) were transfected with siControl (siCtrl) or siSREBF2 and then treated with DMSO or 1 μM U18666A and DMSO or 2 μM pitavastatin for 72 hours, and cell density was measured by SRB assay. Data are represented as mean ± SD (N=3 technical replicates). Statistical analysis was performed using two-way analysis of variance (ANOVA) with Šidák's multiple comparison test (***, $p = 0.0002$, ****, $p < 0.0001$). **c,** Immunoblots of NPC1, HMGCR, SREBP-2, pP70S6K$^{Thr389}$, unprenylated RAP1A, pS6$^{Ser240/244}$, p4E-BP1$^{Thr37/46}$ and β-actin in a panel of TNBC cells (SUM159, MDA-MB-468, BT20) transfected with siControl (siCtrl) or siSREBF2 and siCtrl or siNPC1 for 24 hours and then treated with DMSO or 1 μM U18666A and DMSO or 2 μM pitavastatin for 24 hours.
